# The Contribution of Environmental Enrichment to Phenotypic Variation in Mice and Rats

**DOI:** 10.1101/2020.07.11.198705

**Authors:** Amanda C. Kentner, Amanda V. Speno, Joanne Doucette, Ryland C. Roderick

## Abstract

The reproducibility and translation of neuroscience research is assumed to be undermined by introducing environmental complexity and heterogeneity. Rearing laboratory animals with minimal (if any) environmental stimulation is thought to control for biological variability but may not adequately test the robustness of our animal models. Standard laboratory housing is associated with reduced demonstrations of species typical behaviors and changes in neurophysiology that may impact the translation of research results. Modest increases in environmental enrichment (EE), mitigate against insults used to induce animal models of disease, directly calling into question the translatability of our work. This may in part underlie the disconnect between preclinical and clinical research findings. Enhancing environmental stimulation for our model organisms promotes ethological natural behaviors but may simultaneously increase phenotypic trait variability. To test this assumption, we conducted a systematic review and evaluated coefficients of variation between EE and standard housed mice and rats. Given findings of suboptimal reporting of animal laboratory housing conditions, we also developed a methodological reporting table for enrichment use in neuroscience research. Our data show that animals housed in environmental enrichment were not more variable than those in standard housing. Therefore, environmental heterogeneity introduced into the laboratory, in the form of enrichment, does not compromise data integrity. Overall, human life is complicated and by embracing such nuanced complexity into our laboratories we may paradoxically improve upon the rigor and reproducibility of our research.

**Significance Statement:** Environmental complexity is thought to increase phenotypic variability, undermining research translation. We conducted a systematic review and compared coefficients of variation between environmentally enriched and standard housed laboratory animals. Despite there being no differences in variability across several phenotypic traits, there are stark contrasts in the display of ethological natural behaviors between these housing conditions. Environmental enrichment is recognized as being beneficial for animal welfare and mitigates against insults used to induce animal models of disease. In contrast, standard laboratory cages are recognized as being impoverished and ‘unnatural’. From these observations, it is apparent that our current “gold standard” caging system is not a true control condition as it does not adequately test the robustness of our animal models.

## Introduction

Contributions to phenotypic variation are thought to derive not only from genotype but from multiple environmental factors that range from feeding and microbiology, to variables as seemingly simple as housing condition. In experimental research, scientists attempt to control factors presumed to have an impact on biological variation and consequently the reproducibility of their data. One way to control for phenotypic variability in the laboratory is to standardize animal caging systems and limit environmental complexity. Environmental enrichment (EE) is one form of complexity that includes physical, sensory, cognitive, and/or social stimulation which provides an enhanced living experience to laboratory animals, relative to standard housing conditions. The use of EE has become prominent in neuroscience, due to substantial evidence that EE influences structural and functional changes in the brain, in addition to engendering enduring effects on behavior (Kemperman, 2019; Nithianantharajah & Hannan, 2006). Provisioning supplementary resources to animals not only maintains their welfare but promotes more naturalistic species typical behavioral repertoires (Bloomsmith et al., 2018). Moreover, this enhanced rearing condition has been used to study the mitigative potential of the environment in a variety of animal disease models (Nithianantharajah & Hannan, 2006).

Regardless of the purpose of its use, there are questions about potential within- and between-experiment variability that may accompany the addition of environmental complexity to animal laboratory cages (Kempermann, 2019; Bayne & Würbel, 2014, Grimm, 2018; Toth, 2015; Toth et al., 2011; Sparling et al., 2020). It is thought that the diverse phenotypes promoted by EE may lead to data variation within a study. Moreover, the variety in enrichment protocols used may create data variability between studies and laboratories, compromising data reproducibility. Together, these concerns foster arguments to maintain barren cages as the ‘gold’ standard housing condition (Bayne & Würbel, 2014; Voelkl et al., 2020). Importantly, similar justifications (of increased variation) have been used to support the exclusion of studying females in research, due to hormonal fluctuations across the reproductive cycle. However scientific evidence has since shown this perspective to be incorrect (Becker et al., 2016; Beery, 2018).

Given the shifting attention of the scientific community to the topic of rigor and reproducibility (Toth, 2015; Voelkl et al., 2020), this is the perfect time to reconsider our assumptions about variation due to environmental complexity. Standardization of the environment intuitively falls in line with the scientific method. Parsing out contributors of extraneous variation (Phenotype (*P*) = Gene x Environmental interactions; *G* × *E*) is thought to increase statistical power and reproducibility between experiments. On the other hand, such standardization leads to homogeneity in a population and may undermine the robustness of the potential treatment being studied (Kentner et al., 2018; see Voelkl et al., 2020 for an excellent recent review), a crucial concern given the disconnect between preclinical and clinical research outcomes (Berk, 2012; Hyman, 2012; Munos, 2013).

Still, to control for potential variability, efforts to standardize the environment continues. These efforts have been complicated by varying definitions of what is enriching to animals of each species, strain, and sex (Simpson & Kelly, 2011; Toth, 2015; Toth et al., 2011), even for standard laboratory housing where only minimal EE is recommended or required. Moreover, a lack of reporting on what types of enrichment protocols are used (e.g., shelters, nesting materials, cage mates, music, food/treats; Toth, 2015) make this task even more difficult. Overall, the differential implementation of EE in experimental design has provoked discussion over the inconsistent definitions and reporting methodology of enrichment use in the neuroscience literature, and whether standardization and minimization of laboratory caging is necessary to prevent further extraneous biological variation (Bayne & Würbel, 2014; Toth, 2015).

Outside of theoretical debates, data on whether EE contributes to the replication crisis, by increasing phenotypic variability and undermining research findings, is mixed (Toth, 2015; Toth et al., 2011; Walsh & Cummins, 1979; Wolfer et al., 2004; Würbel, 2007) and concerns about its use persist (Grimm, 2018). Recently, there has been a call to action suggesting that the question of biological variation and its impact on rigor and reproducibility be extended to the diversification of environmental conditions or “controlled heterogenization” (Voelkl et al., 2020). For example, diversification may be implemented by using different sexes, animal strains, ages, and even housing conditions (e.g., EE) within a study. One way to address the question of variability due to the implementation of EE is to utilize the methods of others who have conducted large scale evaluations comparing between male and female animals (Becker et al., 2016) and inbred versus outbred strains of mice (Tuttle et al., 2018). Indeed, it has been noted that the EE literature has typically focused on mean 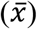 differences between groups, rather than evaluating whether EE increases variability specifically (Kempermann, 2019). Of the small subset that have studied variation directly (e.g., Wolfer et al., 2004; Würbel, 2007; André et al., 2018) they have so far focused on mice and on a limited number of strains within the confines of their own experiments. To our knowledge, there has been no systematic literature-wide evaluation of multiple traits comparing EE to standard housed groups across species.

## Materials and Methods

To evaluate whether EE housed rats or mice display increased phenotypic variability in neuroscience research, we conducted a systematic review and compared the coefficient of variation (*CV*), a measure of trait-specific variability, extracted from data where EE animals were directly compared to a standard (control) housed condition on the same trait. First, to determine the general scientific interest in EE protocols, the proportion of articles published each year, using the search term “environmental enrichment” was identified in PubMed (Sperr, 2016).

### Search Strategy

Both PubMed and EMBASE were searched from the period of January 1^st^, 2013 to September 5, 2018, the date when these searches were initiated. The period evaluated is comparable to other important systematic reviews that assessed phenotypic variability (Becker et al., 2016). We used the search terms (1) environmental enrichment AND (2) electrophysiology OR (3) brain OR (4) behavior OR (5) “nervous system physiological phenomena”, which yielded 3,650 articles (*Figure 1*).

**Figure 1.**
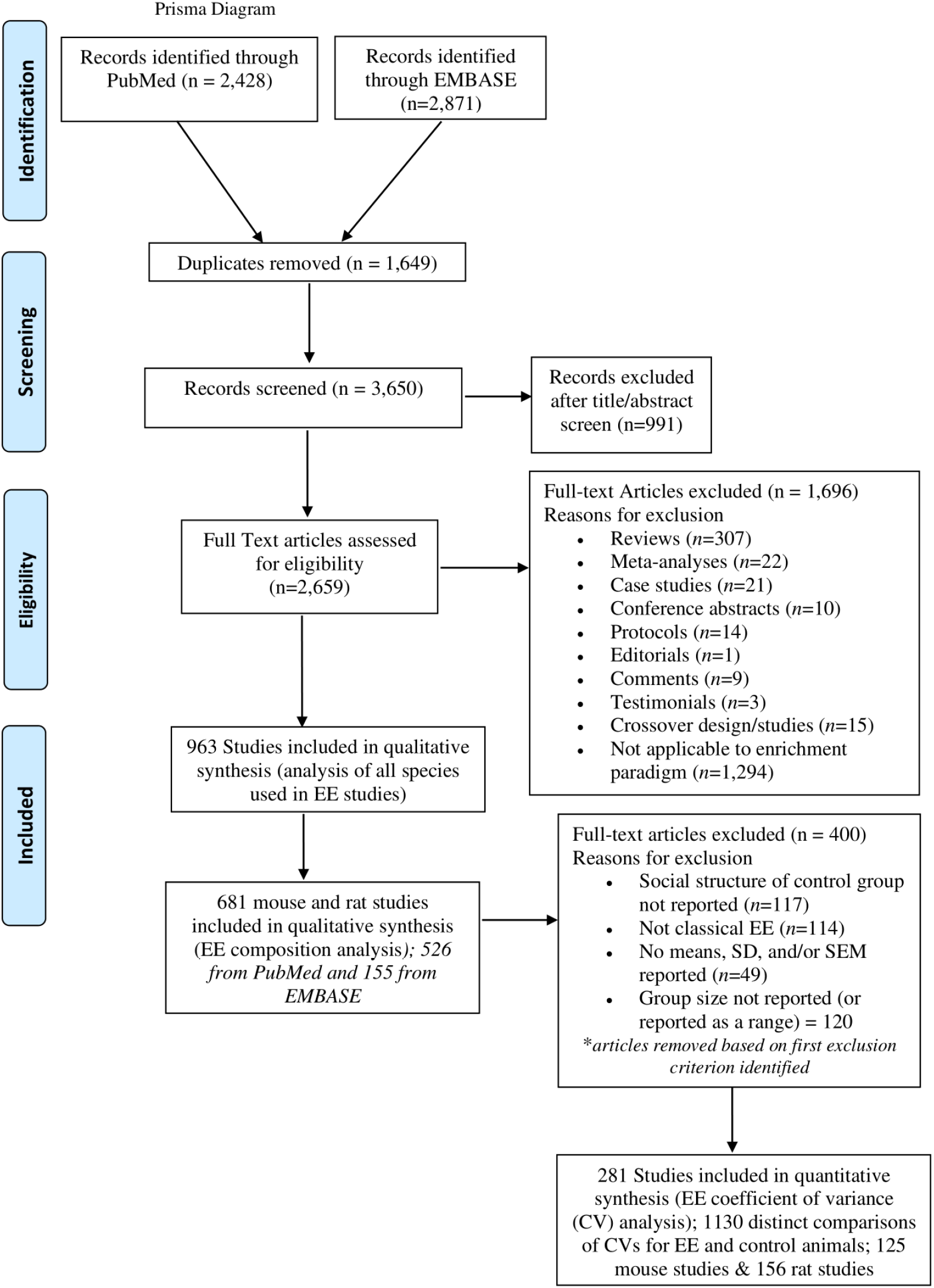
Prisma Flow Diagram.

### Study Selection

After duplicates were removed, evaluators independently identified studies eligible for inclusion in a 2-step process. First, we conducted an abstract and title search. If insufficient details were provided in the titles and abstracts, then the study was selected for full text review. Eligibility was based on (1) article relevance to the subject matter of interest (EE), (2) studies using any animal species including humans, (3) observational and experimental studies, and (4) English-written articles only. Exclusion criteria consisted of reviews, meta-analyses, case studies, conference abstracts, protocols, editorials, comments, and non-English articles. Overall, the articles included in this systematic review were primarily from the fields of neuroscience and animal welfare (see Figure 2 Extended Data Tables 2-1, 2-2).

**Figure 2.**
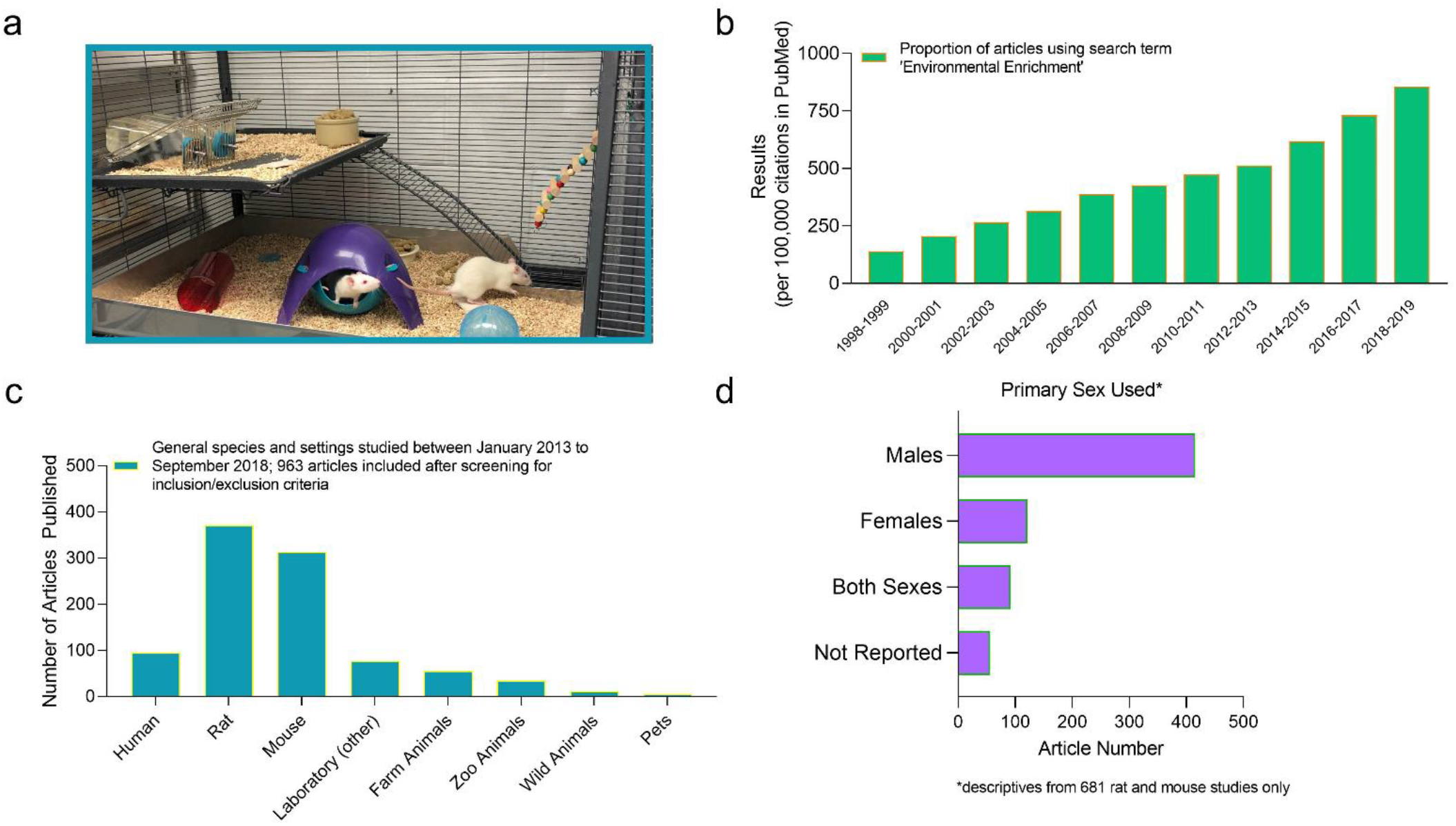
Descriptive analysis of common environmental enrichment (EE) use in research. (a) Picture of a classic EE cage set-up for rodents. (b) Proportion per 100,000 citations of PubMed articles returned when searching “environmental enrichment”. Graph depicts articles published between 1998 and 2019. Includes both primary and secondary sources (an update from Simpson & Kelly, 2011). Graphs depict the (c) general species and settings and (d) primary sex studied using EE between January 2013 and September 2018. The selected articles used in this study are primarily from the areas of neuroscience and animal welfare (Extended Data Tables 2-1, 2-2).

### Data Extraction

Of the 963 articles identified as using EE in any species, a subset of 681 articles were identified as using mice or rats and were further evaluated on their use of several methodological variables including sex, types of enrichment devices employed, in addition to social structure of the EE group and composition of the control conditions used (e.g., running wheel, isolated, social/group housing). Phenotypic variability was also evaluated on the rat and mouse studies identified as using traditional EE caging systems (*Figure 2A*). For these analyses, 281 studies were evaluated based on meeting the inclusion criteria of providing means and standard deviations (or standard errors) that could be extracted from the article, and sample sizes for at least one EE and one control group (*Figure 1*). We also identified whether EE and control groups were naïve or ‘treated/manipulated’ (e.g., drug treated, knockout models, surgery etc.). Studies with parental exposure to EE were excluded to control for potential confounds of parental care (Connors et al., 2015), as were studies where it was unclear if control animals were singly or socially housed. To avoid oversampling (Tuttle et al., 2018), we limited data collection to the first three reported measures where data and error bars were clearly legible. Each measure was categorized similarly to how others had done previously (Becker et al., 2016; Tuttle et al., 2018) by using behavior/CNS, behavior/other, anatomy, immune system function, organ function, molecules, and electrophysiology as traits. Generally, the behavior/CNS category included measures where animals demonstrated some type of learning, discrimination, or what could be considered more complex sequences of behaviors. Examples from our dataset include time spent with a novel object or social conspecific, sniffing duration, and duration of social contact (e.g., discrimination and preference tests). Number of lever responses, conditioned place preference scores, latency to locate a platform in the Morris Water Maze, % fear generalization, % freezing time, % sucrose preference, number of reference memory errors etc. were also included in this category. In contrast, the behavior/other category represented measures such as time spent in the center of the open field, frequency of crossings into the open center or periphery, and distance traveled in the open field. Anatomy included measures like the length or volume of brain regions (e.g., dendritic length, corpus callosum thickness). Immune system function as a category included measures such as flow cytometric analysis of CD40 on peritoneal macrophage, tumor volume or weight. We also placed plasma cytokine levels into this category. The organ function category included heart rate, changes in arterial blood P02, PC02, and pH, as well as fasting blood glucose levels. Molecules included any other measures of protein or mRNA, for example. These latter measures were primarily localized to the brain in our dataset.

In total there were 1130 direct comparisons of coefficients of variation (*CV)*s between EE and control animals included here (618 naïve pair comparisons and 512 manipulated/treated pair comparisons; *Figure 1*). The number of articles included, and direct comparisons made, in our analyses surpassed other excellent systematic reviews evaluating phenotypic variability (Becker et al., 2016; Tuttle et al., 2018). Therefore, we have an adequate sample size to make appropriate conclusions. Data were extracted from graphs provided on digital PDF articles (using http://rhig.physics.yale.edu/~ullrich/software/xyscan/), or directly from tables. Graphical data extractions were performed by two trained researchers. Inter-rater reliability was assessed, and Pearson *r* correlation was determined to range from 0.912-0.997.

### Statistical Analyses

*CV*s were calculated as standard deviation divided by the mean and compared using paired t-tests (for individual trait evaluations), or ANOVA (for multiple trait evaluations). Pairwise comparisons were done using the Tukey’s multiple comparisons test (Becker et al., 2016; Howell, 2001). The partial eta-squared (η^2^) is also reported as an index of effect size for the ANOVAs (the range of values being 0.02 = small effect, 0.13 = moderate effect, 0.26 = large effect; Miles and Shevlin, 2001). To determine whether the distribution of variation differed by environmental complexity, we calculated EE to control ratios of *CV* = [(*CV_EE_)/(CV_EE_ + CV_control_*)]. *CV* ratios for each trait were tested as a function of housing complexity against the theoretical mean 0.5 by t test (Becker et al., 2016; Beery, 2018). Data were considered significant if p<0.05.

## Results

Using the term “environmental enrichment” we identified the proportion of articles indexed in PubMed each year from 1998 to 2019 (Sperr, 2016). One report has previously evaluated the number of articles published from 1960-2009 (Simpson & Kelly, 2011). In this work, it was demonstrated that an increased interest in environmental enrichment emerged between 1990-1999 and 2000-2009. Here, we provide a replication and extension of those data from 1998 until 2019. Our search, including both review and empirical research articles, highlights a continuation of the increasing interest on this topic, relative to the number of total articles published (*Figure 2B*).

The results of our analyses demonstrate patterns of experimental biases, specifically a heavy reliance on the use of rats and mice over other laboratory species (*Figure 2C*), and the continued exclusion of females in EE research (*Figure 2D*; Simpson & Kelly, 2011). Our findings also show a range in the definition of EE used across laboratories in that the frequency of enrichment types, timing, and the social structures implemented varied widely (*Figure 3A-F*). The use of toys (including plastic or wooden), bones/chews, house hideaways or tubes/pipes and tunnels, in addition to a larger cage space and social conspecifics were more frequently used in the enrichment housing conditions. Supplementary bedding/nesting materials and ramps/ladders or perches were less commonly used, as were swings, ropes and chains (*Figure 3A*).

**Figure 3.**
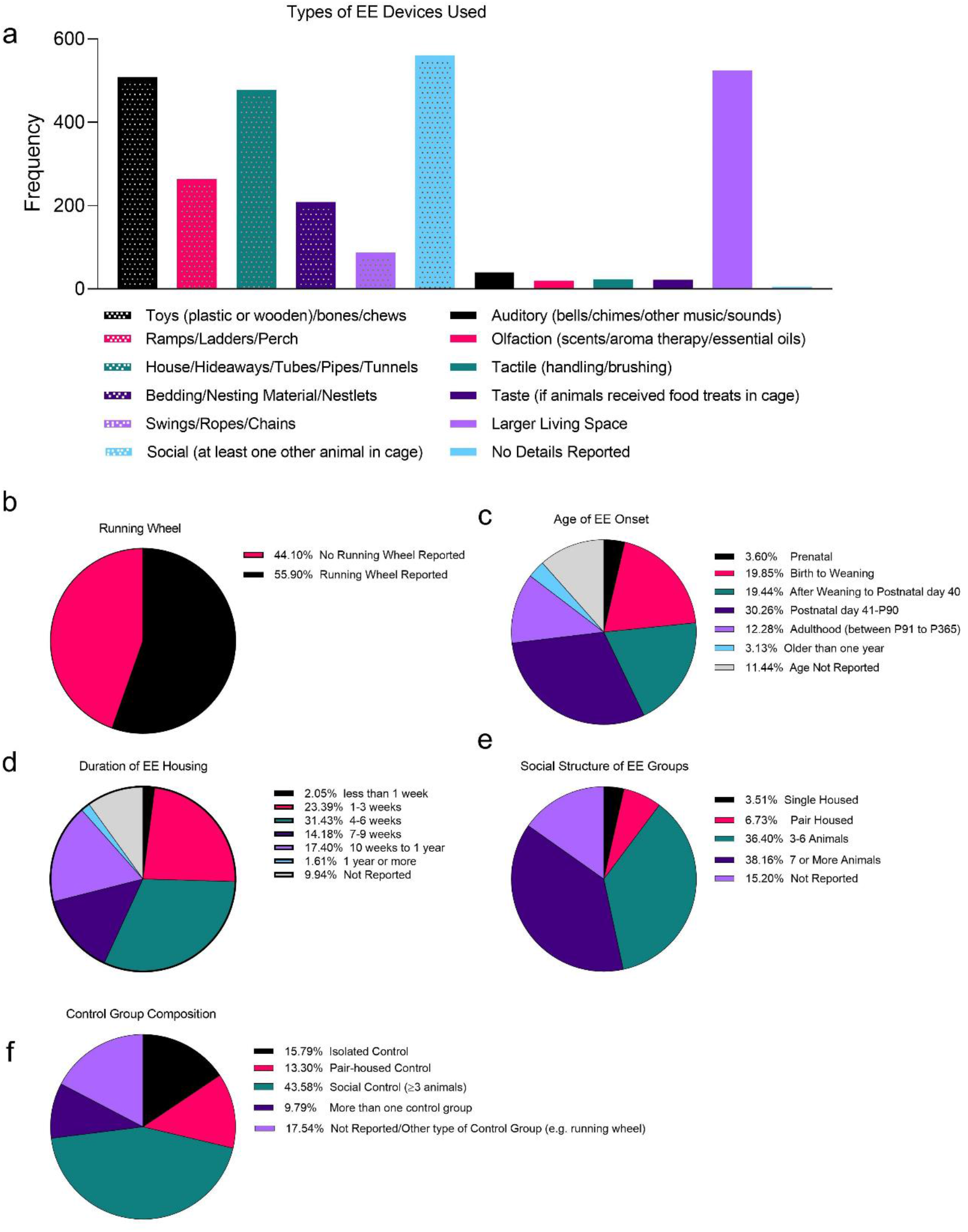
Descriptive analysis of common environmental enrichment (EE) methodology. All descriptive data are from rat and mice studies where animals are housed in a classic EE design. Data outline the (a) frequency of types of EE devices used, in addition to the percentage of EE studies using (b) running wheels, or a particular (d) age of EE onset, (d) duration of EE housing, and general/social structure of (e) EE and (f) control groups. Data derived from a total of 681 research articles published between January 2013 and September 2018.

One issue that arose was a significant lack of reporting on several variables. This prompted us to develop a reporting table for describing key aspects of enrichment use in research (*Table 1*), following suit with other initiatives to improve on animal model reporting (Kentner et al., 2019). As part of this table, we suggest authors report if they are providing EE animals with manufactured/artificial enrichment devices or more natural stimuli as there are differences in animal phenotypes depending on these devices (Lambert et al., 2015; Hess et al., 2008).

**Table 1.** Environmental Enrichment Reporting Guidelines Checklist. The recommended use of this reporting form is to fill it out and include it as supplemental material for each of your laboratory’s environmental enrichment research publications. This document can also be used as a guide for including details of cage enrichment for studies utilizing only standard laboratory housing. If there are difficulties utilizing/adapting this form, please contact one of the corresponding authors to request a copy.

| Reporting Item | Description |
| --- | --- |
| *EE cage size (LxWxH) in cm* |  |
| *Control cage size (LxWxH) in cm* |  |
| *Material of EE cages* |  |
| *Material of control cages* |  |
| *Type of EE caging system (e.g. open top/individually ventilated)* |  |
| *Type of control caging system (e.g. open top/individually ventilated)* |  |
| *Number & sex of animals housed in each EE cage* |  |
| *Number & sex of animals housed in each control cage* |  |
| *Enrichment devices (e.g., toys, shelters, ramps, running wheels, perches, or different levels/shelves) in EE cages. What type of materials were devices made of (e.g., wood, plastic, are they manufactured/artificial, or are the natural stimuli). How many enrichment devices were typically in the cage?* |  |
| *Enrichment devices e.g., toys, shelters, ramps, running wheels, perches or different levels/shelves) in control cages. What type of materials were devices made of (e.g., wood, plastic, are they manufactured/artificial, or are the natural stimuli). How many enrichment devices were typically in the cage?* |  |
| *How frequently were enrichment devices changed in EE cages?* |  |
| *How frequently were enrichment devices changed in control cages?* |  |
| *Bedding type and other nesting materials in EE cages* |  |
| *Bedding type and other nesting materials in control cages* |  |
| *Age of animals when placed into EE housing* |  |
| *Duration housed in EE cages* |  |
| *Frequency of cage cleanings for EE animals* |  |
| *Frequency of cage cleanings for control animals* |  |
| *Were EE and control animals housed in the same room or different?* |  |
| *Specific number of EE and control animals used for each measure reported* |  |
| *Other notes/comments (e.g., other types of enrichment given, for example music in animal holding room, food treats)* |  |

Using paired t-tests, we found no differences between EE and standard housed mice or rats on *CVs* across traits (p>0.05), regardless of control housing type (e.g., running wheel, isolated, social/group housing) or whether animals were naïve or manipulated/treated (e.g., drug treated, knockout models, surgery). Therefore, we collapsed and analyzed both species together. When species were combined, the treated/manipulated social/group housed controls (0.65 ±0.073) were more variable than their manipulated/treated EE counterparts (0.59 ± 0.050; t(46) = 2.211, p = 0.032) on the “behavior other” trait only. Isolated control animals (0.24 ± 0.079) had higher *CVs* than treated/manipulated EE animals on the anatomy trait (0.019 ± 0.072; (t(4) = 4.720, p = 0.009). However, for the anatomy trait the number of available comparisons between these two groups was not sufficiently powered (n = 5 comparisons based on 3 articles). In general, we did not find EE to increase trait variability compared to any control housing type in either naïve or manipulated/treated animals (p>0.05).

To increase the power in our analyses, we collapsed the control group types together and analyzed across species and traits, both separately and together. Again, we found that EE does not make animals more variable than controls (p> 0.05; *Figure 4A-D*; Extended Data Tables 4-1, 4-2, 4-3, 4-4, 4-5, 4-6, 4-7, 4-8, 4-9, 4-10, 4-11, 4-12, 4-13, 4-14, 4-15, 4-16, 4-17, 4-18, 4-19, 4-20, 4-21, 4-22, 4-23, 4-24, 4-25). When species were combined, we found that controls were more variable (had higher *CVs*) than EE housed animals under treated/manipulated conditions. However, this was only found on the “overall behavior” (main effect of housing: t(290) = 2.120, p = 0.035; Control *CV*: 0.67 ± 0.06/EE *CV*: 0.56 ± 0.04) and “behavior other” traits (main effect of housing: t(46) = 2.211, p = 0.032; Control *CV*: 0.73 ± 0.07/EE *CV*: 0.60 ± 0.05, based on 21 articles; *Figure 4BD*; Extended Data Table 4-4).

**Figure 4.**
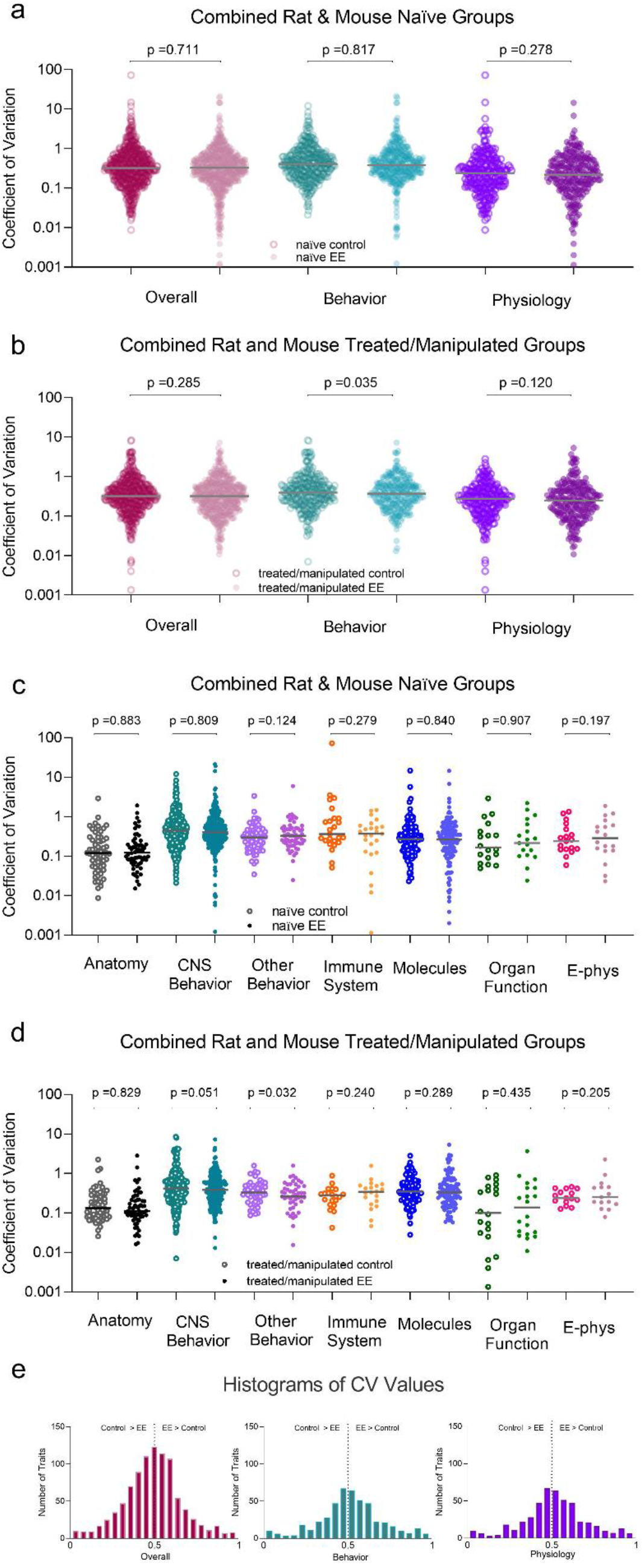
Coefficient of variance for all studies where control and environmentally enriched (EE) mice or rats were directly compared. All data presented as the overall trait variance and further separated into subcategories of behavior and physiology as well as seven specific trait measures for (a, c) naïve/untreated and (b, d) treated/manipulated animals (mean ±SEM). Each data point represents a single control or EE measure from a single experiment along with the mean for each respective trait. Coefficient of variance was calculated as the standard deviation divided by the mean for each data point. (e) Histogram of distribution of CV ratios (EE CV/(EE CV + control CV) collapsed across naïve and treated/manipulated mice and rats. To determine whether the variance from the mean was normally distributed for the different traits, we evaluated the CV ratios (P values from Extended Data Tables 4-1, 4-2, 4-3, 4-4, 4-5, 4-6, 4-7, 4-8, 4-9, 4-10, 4-11, 4-12, 4-13, 4-14, 4-15, 4-16, 4-17, 4-18, 4-19, 4-20, 4-21, 4-22, 4-23, 4-24, 4-25). A value of 0.5 (black dotted line) indicates that EE and control animals are similar. Values to the right suggest that EE animals are more variable than controls.

There were no main effects of housing, nor significant housing by trait interactions on the two-way ANOVAs (p> 0.05, Figure 4 Extended Data Table 4-13). However, there were significant main effects of trait, indicating that “behavior” was more variable than “anatomical” traits for both rats (main effect of trait: F(5, 542) = 4.015, p = 0.001, η^2^ = 0.036; Tukey HSD: p = 0.004) and mice (main effect of trait: F(6, 460) = 4.953, p = 0.0001, η^2^ = 0.057; Tukey HSD: p = 0.001; Figure 4 Extended Data Table 4-13). Of special note, partial η^2^ values were indicative of small effect sizes for these comparisons.

Although the inclusion of female animals was demonstrably lower than males to be able to make adequately powered comparisons on many traits (*Figure 2D*), we conducted some preliminary sex difference analyses. Our sub-analyses revealed that naïve male EE rats (0.60 ± 0.10) had higher *CVs* than their naïve social/group housed controls (0.39 ± 0.18; t(32) = −2.266, p = 0.030, based on 18 articles) on the “behavior other” trait, but were not more variable on any other trait (p>0.05). There were no further differences in variability between EE and control animals across any combination of sex, strain, control type, or naïve vs treated/manipulated animals (where n> 5 direct EE vs control comparisons).

When comparing *CV* ratios, the data did not support the premise that environmental complexity increases variability in neuroscience research (p >0.05; *Figure 4E*; Extended Data Tables 4-14, 4-15, 4-16, 4-17, 4-18, 4-19, 4-20, 4-21, 4-22, 4-23, 4-24, 4-25).

## Discussion

Our findings should resonate well with neuroscientists who would like to increase complexity in laboratory caging systems, promoting more naturalistic species typical behaviors and brain functioning, but who have been concerned about compromising data integrity and their control over environmental conditions. This should be especially salient given that lack of enrichment in laboratory cages leads to suppression of behavioral repertoires, increased stereotypies, and a reduction of general activity level, even during an animals’ active phase (Hurst et al., 1997). Indeed, deprivation in the environment is known to impact the structure and functioning of the brain, affecting cognition and behavior (McLaughlin et al., 2017; Lahvis, 2017). This underscores the view that our current standard laboratory housing condition is not a true control condition. Cage enrichment is recommended in the Guide for the Care and Use of Laboratory Animals (NRC, 2011), and for standard housed rodents typically takes the form of sanitizable polyvinyl chloride (PVC) tubes, a chew bone, or a piece of nesting material. If the animal is lucky, they may receive a combination of two or three pieces of these enrichment devices. To be frank, the composition of this housing condition needs a major renovation. Seldom do these cage enrichment objects change across the course of the study; novelty and increased stimulation are luxuries afforded to animals reared in classic EE (see *Figure 2A*). This latter housing condition is rarely utilized as a standard in the laboratory; when employed, EE is usually for the purpose of exploring mechanisms underlying neural plasticity, or to mitigate some type of toxic insult (Nithianantharajah & Hannan, 2006). The availability of resources is a major restriction to increasing stimulation in the animal laboratory. It will require a change in the mindsets of institutions, scientists, and funding bodies to make this housing condition, or an adapted version, the new “gold standard”. Some solutions to address cost, physical space, as well as personnel constraints to implementing higher levels of enrichment have been discussed elsewhere (Kentner et al., 2018) and are outlined below. Still, the direction of funds to establish more complex housing conditions for laboratory animals should be part of the movement to improve scientific rigor and reproducibility.

Another important hurdle to the implementation of EE is concerns about phenotypic variability due to increased heterogeneity. While we identified some increased variability in naïve male EE rats on measures such as distance traveled and open field, most studies evaluated utilized some type of experimental treatment/manipulation which did not affect phenotypic variability on any trait. Moreover, others have reported no differences in variability on these types of measures, when associated with EE use, at least in mice (André et al., 2018; Würbel, 2007; Wolfer et al., 2004). In general, this species and sex specific effect suggests that researchers may need to identify the appropriate EE devices to use for male rats in some experimental designs, to resolve potential issues in variability. Notably, we observed increased phenotypic variability on the ‘overall behavior’ and “behavior other” traits in control housed animals. Therefore, complex housing does not make animals any more variable in comparison to standard laboratory housing. One consideration with respect to our data is that a larger proportion of our analyses were behavioral measures, versus cellular or molecular. However, these latter measures were also equally unaffected by housing condition. Still, our interpretations are limited by the fact that we summarized many studies and that these overall findings may not be applicable to individual experiments. Moreover, factors such as age of EE onset, animal age at endpoint evaluation, strain differences, as well as other species differences are important contributions to EE that may affect our interpretations. Another potential contributor to the shaping of phenotype could be the shared experiences in EE, resulting in within-group differences. Individual animals influence their environment, and each other, affecting phenotypes and preventing full control of the environment. Therefore, EE could be considered not just as *P* = *G x E*, but as *G* x (*E_shared_* + *E_nonshared_*); see Kempermann, 2019 for an excellent review. This equation is also relevant to pair and grouped standard laboratory cage housing, which do not increase phenotypic trait variability (Becker et al., 2016), similarly to what we show here with more naturalistic settings.

Together, the main complaints against the implementation of EE have been about feasibility and associated financial costs, in addition to arguments of increased phenotypic variability as a result of modeling more naturalistic settings in the laboratory environment (Grimm, 2018). However, EE may not need to be extravagant or require larger caging systems and space but may be as simple as regularly changing enrichment devices (Kentner et al., 2018). Notably, investigators often group house their animals to reduce stress (Hurst et al., 1997); and consequentially save on laboratory caging costs. One option is to use bigger cages that take up the same space/area as multiple smaller cages. Animals can then be grouped together in larger colonies. While this can serve to increase social enrichment, its implementation must also keep the needs of each species and sex in mind. For example, issues of social hierarchy and dominance are more likely to occur in males of some species and social stress experiences can greatly affect overall health and disposition (Beery et al., 2020; Larrieu et al., 2017; Zhou et al., 2018). Species such as CD-1 mice will become territorial when enrichment devices are introduced into the environment, disrupting their established hierarchy (McQuaid et al., 2012). These types of species may otherwise live cooperatively in a larger group when the social hierarchy is firmly established (McQuaid et al., 2012; see Beery et al., 2020 for an excellent review on groups and non-traditional housing models). Importantly, most EE studies begin to offer increased stimulation at weaning, or shortly after puberty (see *Figure 3C*). In at least some animal models, when higher levels of stimulation are the norm across the entire lifespan, versus being introduced after adolescence, fighting has been reported to be non-existent in both male and female mice and rats. This has allowed for the peaceful use of enrichment devices among these groups (Zhao et al., 2020; Kentner et al., 2016; Connors et al., 2014).

From a purely scientific perspective, EE can mitigate the effects of several experimental treatments and animal models of disease (Nithianantharajah & Hannan, 2006) and is often interpreted as a beneficial intervention (Sparling et al., 2020). However, this calls into question the external validity of these apparent context specific effects (Bernard, 2019; Manouze et al., 2019) and the robustness of our animal models; a clear example of fallacious reasoning (Bernard, 2020). Indeed, incorporating more environmental heterogeneity into neuroscience research, and testing our findings against such complexity, should increase the robustness of our experimental designs and the fidelity of biomedical treatments (Kentner et al., 2018; Voelkl et al., 2020), without compromising the underlying stability of data. Our study supports this idea given that traditional EE caging systems are dynamic environments where devices are being replaced or are changing location as animals interact and move them. Moreover, social experiences are varied for each animal. Specifically, experiences both between and within EE cages are unique, yet complex housing does not make animals any more variable compared to standard laboratory housed rats or mice. Importantly, the increased use of EE and improved robustness of experimental design should be less costly in the long run. This contrasts with a continued reliance on standard laboratory housing, which is clearly not a true control condition and appears to impede the translation of research results.

Going forward, it will be necessary to identify appropriate enrichment types for the species, sex, and age of the model organism of interest, in addition to the animal model/paradigm being used, and to accurately report their use (Kentner et al., 2018; Simpson & Kelly, 2011; Toth, 2015). Importantly, there are proposed methodologies for how to implement and account for such environmental variation (Voelkl et al., 2020). Overall, human life is complicated and by embracing such nuanced complexity into our laboratories we may paradoxically improve upon the rigor and reproducibility of our research.

## Supporting information

Extended Table 2-1. List of PubMed References Used

Extended Table 2-2. List of EMBASE References Used

Extended Data File 4-1. Pairwise Comparisons

Extended Data File 4-2. Pairwise Comparisons

Extended Data File 4-3. Pairwise Comparisons

Extended Data File 4-4. Pairwise Comparisons

Extended Data File 4-5. Pairwise Comparisons

Extended Data File 4-6. Pairwise Comparisons

Extended Data File 4-7. Pairwise Comparisons

Extended Data File 4-8. Pairwise Comparisons

Extended Data File 4-9. Pairwise Comparisons

Extended Data File 4-10. Pairwise Comparisons

Extended Data File 4-11. Pairwise Comparisons

Extended Data File 4-12. Pairwise Comparisons

Extended Data File 4-13. Two Way ANOVAs

Extended Data File 4-14. Coefficient of Variation (CV)s

Extended Data File 4-15. Coefficient of Variation (CV)s

Extended Data File 4-16. Coefficient of Variation (CV)s

Extended Data File 4-17. Coefficient of Variation (CV)s

Extended Data File 4-18. Coefficient of Variation (CV)s

Extended Data File 4-19. Coefficient of Variation (CV)s

Extended Data File 4-20. Coefficient of Variation (CV)s

Extended Data File 4-21. Coefficient of Variation (CV)s

Extended Data File 4-22. Coefficient of Variation (CV)s

Extended Data File 4-23. Coefficient of Variation (CV)s

Extended Data File 4-24. Coefficient of Variation (CV)s

Extended Data File 4-25. Coefficient of Variation (CV)s

Extended Data File 4-26. Coefficient of Variation (CV)s

## Data Availability

All data are available upon request.

## Code Availability

There is no code associated with this work.

---

**Extended Data Table 2-1.** List of PubMed References Used.

**Extended Data Table 2-2.** List of EMBASE References Used.

**Extended Data Table 4-1**. Pairwise comparisons for naïve controls and naïve enriched rats and mice in which all behavior, physiology, and anatomy traits are combined.

**Extended Data Table 4-2**. Pairwise comparisons for treated/manipulated controls and treated/manipulated enriched rats and mice in which all behavior, physiology, and anatomy traits are combined.

**Extended Data Table 4-3**. Pairwise comparisons for naïve controls and naïve enriched rats and mice by each individual trait.

**Extended Data Table 4-4**. Pairwise comparisons for treated/manipulated controls and treated/manipulated enriched rats and mice by each individual trait.

**Extended Data Table 4-5**. Pairwise comparisons for naïve control and naïve enriched rats in which all behavior, physiology, and anatomy traits are combined.

**Extended Data Table 4-6**. Pairwise comparisons for treated/manipulated control and treated/manipulated enriched rats in which all behavior, physiology, and anatomy traits are combined.

**Extended Data Table 4-7**. Pairwise comparisons for naïve controls and naïve enriched rats by each individual trait.

**Extended Data Table 4-8**. Pairwise comparisons for treated/manipulated controls and treated/manipulated enriched rats by each individual trait.

**Extended Data Table 4-9**. Pairwise comparisons for naïve controls and naïve enriched mice in which all behavior, physiology, and anatomy traits are combined.

**Extended Data Table 4-10**. Pairwise comparisons for treated/manipulated controls and treated/manipulated enriched mice in which all behavior, physiology, and anatomy traits are combined.

**Extended Data Table 4-11**. Pairwise comparisons for naïve controls and naïve enriched mice by each individual trait.

**Extended Data Table 4-12**. Pairwise comparisons for treated/manipulated controls and treated/manipulated enriched mice by each individual trait.

**Extended Data Table 4-13**. Two-way ANOVAs comparing multiple traits (all behavior, physiology, anatomy) by housing condition (environmental enrichment, standard housing) for the independent variable coefficient of variation (CV). Data presented for both rats and mice combined and separately.

**Extended Data Table 4-14**. Coefficient of variation (CV) distributions for naïve standard housed (controls) and naïve environmental enriched (EE) rats and mice combined with treated/manipulated controls and treated/manipulated EE rats and mice in which all behavior, physiology, and anatomy traits are combined. CV ratios were used to determine whether the distribution of variation differed by environmental complexity. Calculated EE to control ratios of *CV* = [(*CV_EE_)/(CV_EE_ + CV_control_*)]. CV ratios tested as a function of housing complexity against the theoretical mean of 0.5 by a one-sample t-test.

**Extended Data Table 4-15**. Coefficient of variation (CV) distributions for naïve standard housed (controls) and naïve environmental enriched (EE) rats and mice in which all behavior, physiology, and anatomy traits are combined. CV ratios were used to determine whether the distribution of variation differed by environmental complexity. Calculated EE to control ratios of *CV* = [(*CV_EE_)/(CV_EE_ + CV_control_*)]. CV ratios tested as a function of housing complexity against the theoretical mean of 0.5 by a one-sample t-test.

**Extended Data Table 4-16**. Coefficient of variation (CV) distributions for treated/manipulated standard housed (controls) and treated/manipulated environmental enriched (EE) rats and mice in which all behavior, physiology, and anatomy traits are combined. CV ratios were used to determine whether the distribution of variation differed by environmental complexity. Calculated EE to control ratios of *CV* = [(*CV_EE_)/(CV_EE_ + CV_control_*)]. CV ratios tested as a function of housing complexity against the theoretical mean of 0.5 by a one-sample t-test.

**Extended Data Table 4-17**. Coefficient of variation (CV) distributions for naïve standard housed (controls) and naïve environmental enriched (EE) rats and mice by each individual trait. CV ratios were used to determine whether the distribution of variation differed by environmental complexity. Calculated EE to control ratios of *CV* = [(*CV_EE_)/(CV_EE_ + CV_control_*)]. CV ratios tested as a function of housing complexity against the theoretical mean of 0.5 by a one-sample t-test.

**Extended Data Table 4-18**. Coefficient of variation (CV) distributions for treated/manipulated standard housed (controls) and treated/manipulated environmental enriched (EE) rats and mice by each individual trait. CV ratios were used to determine whether the distribution of variation differed by environmental complexity. Calculated EE to control ratios of *CV* = [(*CV_EE_)/(CV_EE_ + CV_control_*)]. CV ratios tested as a function of housing complexity against the theoretical mean of 0.5 by a one-sample t-test.

**Extended Data Table 4-19**. Coefficient of variation (CV) distributions for naïve standard housed (controls) and naïve environmental enriched (EE) rats in which all behavior, physiology, and anatomy traits are combined. CV ratios were used to determine whether the distribution of variation differed by environmental complexity. Calculated EE to control ratios of *CV*= [(*CV_EE_)/(CV_EE_ + CV_control_*)]. CV ratios tested as a function of housing complexity against the theoretical mean of 0.5 by a one-sample t-test.

**Extended Data Table 4-20**. Coefficient of variation (CV) distributions for treated/manipulated standard housed (controls) and treated/manipulated environmental enriched (EE) rats in which all behavior, physiology, and anatomy traits are combined. CV ratios were used to determine whether the distribution of variation differed by environmental complexity. Calculated EE to control ratios of *CV* = [(*CV_EE_)/(CV_EE_ + CV_control_*)]. CV ratios tested as a function of housing complexity against the theoretical mean of 0.5 by a one-sample t-test.

**Extended Data Table 4-21**. Coefficient of variation (CV) distributions for naïve standard housed (controls) and naïve environmental enriched (EE) rats by each individual trait. CV ratios were used to determine whether the distribution of variation differed by environmental complexity. Calculated EE to control ratios of *CV* = [(*CV_EE_)/(CV_EE_ + CV_control_*)]. CV ratios tested as a function of housing complexity against the theoretical mean of 0.5 by a one-sample t-test.

**Extended Data Table 4-22**. Coefficient of variation (CV) distributions for treated/manipulated standard housed (controls) and treated/manipulated environmental enriched (EE) rats by each individual trait. CV ratios were used to determine whether the distribution of variation differed by environmental complexity. Calculated EE to control ratios of [(*CV_EE_)/(CV_EE_ + CV_control_*)]. CV ratios tested as a function of housing complexity against the theoretical mean of 0.5 by a one-sample t-test.

**Extended Data Table 4-23**. Coefficient of variation (CV) distributions for naïve standard housed (controls) and naïve environmental enriched (EE) mice in which all behavior, physiology, and anatomy traits are combined. CV ratios were used to determine whether the distribution of variation differed by environmental complexity. Calculated EE to control ratios of *CV* = [(*CV_EE_)/(CV_EE_ + CV_control_*)]. CV ratios tested as a function of housing complexity against the theoretical mean of 0.5 by a one-sample t-test.

**Extended Data Table 4-24**. Coefficient of variation (CV) distributions for treated/manipulated standard housed (controls) and treated/manipulated environmental enriched (EE) mice in which all behavior, physiology, and anatomy traits are combined. CV ratios were used to determine whether the distribution of variation differed by environmental complexity. Calculated EE to control ratios of *CV* = [(*CV_EE_)/(CV_EE_ + CV_control_*)]. CV ratios tested as a function of housing complexity against the theoretical mean of 0.5 by a one-sample t-test.

**Extended Data Table 4-25**. Coefficient of variation (CV) distributions for naïve standard housed (controls) and naïve environmental enriched (EE) mice by each individual trait. CV ratios were used to determine whether the distribution of variation differed by environmental complexity. Calculated EE to control ratios of *CV* = [(*CV_EE_)/(CV_EE_ + CV_control_*)]. CV ratios tested as a function of housing complexity against the theoretical mean of 0.5 by a one-sample t-test.

**Extended Data Table 4-26**. Coefficient of variation (CV) distributions for treated/manipulated standard housed (controls) and treated/manipulated environmental enriched (EE) mice by each individual trait. CV ratios were used to determine whether the distribution of variation differed by environmental complexity. Calculated EE to control ratios of *CV* = [(*CV_EE_)/(CV_EE_ + CV_control_*)]. CV ratios tested as a function of housing complexity against the theoretical mean of 0.5 by a one-sample t-test.

## Acknowledgments

This project was funded by NIMH under Award Number R15MH114035 (to ACK) and a MCPHS Summer Undergraduate Research Fellowship (SURF) awarded to R.C.R. The authors are grateful to Dominic Rainone, Jie Yi Tan, Victoria Perez, Alexandra Best, Madeline Puracchio, and Yvonne Zheng for their help with data collection. The authors would also like to thank the MCPHS University School of Pharmacy and School of Arts & Sciences for their continual support. The content is solely the responsibility of the authors and does not necessarily represent the official views of any of the financial supporters.

## Conflict of Interest

Authors report no conflict of interest.

## Author Contributions

A.C.K. designed and supervised the study and wrote the manuscript. The study was carried out and analyzed by A.V.P, R.C.R., J.D, and A.C.K.

## Notes

### Competing Interest Statement

The authors have declared no competing interest.

### Summary of Updates

We have revised the manuscript by adding more information to the discussion. Supplemental files have been updated.

