## Extended Table 2-1. List of PubMed References Used for "The Contribution of Environmental Enrichment to Phenotypic Variation in Mice and Rats"

**Extended Data Table 3-1.** List of PubMed References Used

| **Study** | **Year** | **Author** | **Journal** |
| --- | --- | --- | --- |
| 1 | 2018 | Ahmadalipour, Ali; Ghodrati-Jaldbakhan, Shahrbanoo; Samaei, Seyed Afshin; Rashidy-Pour, Ali | Neurobiology of learning and memory |
| 2 | 2018 | Anstotz, Max; Lee, Sun Kyong; Neblett, Tamra I.; Rune, Gabriele M.; Maccaferri, Gianmaria | Cerebral cortex |
| 3 | 2018 | Ashokan, Archana; Hegde, Akshaya; Balasingham, Anushanthy; Mitra, Rupshi | Brain research |
| 4 | 2018 | Ashokan, Archana; Lim, Jamien Wee Han; Hang, Nicholas; Mitra, Rupshi | Scientific reports |
| 5 | 2018 | Aujnarain, Amiirah B.; Luo, Owen D.; Taylor, Natalie; Lai, Jonathan K. Y.; Foster, Jane A. | Behavioural brain research |
| 6 | 2018 | Bailoo, Jeremy D.; Murphy, Eimear; Varholick, Justin A.; Novak, Janja; Palme, Rupert; Wurbel, Hanno | Scientific reports |
| 7 | 2018 | Bates, M. L. Shawn; Hofford, Rebeca S.; Emery, Michael A.; Wellman, Paul J.; Eitan, Shoshana | Drug and alcohol dependence |
| 8 | 2018 | Bator, Ewelina; Latusz, Joachim; Wedzony, Krzysztof; Mackowiak, Marzena | European neuropsychopharmacology : the journal of the European College of Neuropsychopharmacology |
| 9 | 2018 | Bengoetxea, Harkaitz; Rico-Barrio, Irantzu; Ortuzar, Naiara; Murueta-Goyena, Ane; Lafuente, Jose V. | Molecular neurobiology |
| 10 | 2018 | Birch, Amy M.; Kelly, Aine M. | Neuropharmacology |
| 11 | 2018 | Bockman, Charles S.; Zeng, Wanyun; Hall, Jamie; Mittelstet, Beth; Schwarzkopf, Liz; Stairs, Dustin J. | Psychopharmacology |
| 12 | 2018 | Bonfiglio, T.; Olivero, G.; Vergassola, M.; Di Cesare Mannelli, L.; Pacini, A.; Iannuzzi, F.; Summa, M.; Bertorelli, R.; Feligioni, M.; Ghelardini, C.; Pittaluga, A. | Neuropharmacology |
| 13 | 2018 | Borniger, Jeremy C.; Ungerleider, Kyra; Zhang, Ning; Karelina, Kate; Magalang, Ulysses J.; Weil, Zachary M. | Neuroscience |
| 14 | 2018 | Brown, Russell W.; Schlitt, Marjorie A.; Owens, Alex S.; DePreter, Caitlynn C.; Cummins, Elizabeth D.; Kirby, Seth L.; Gill, W. Drew; Burgess, Katherine C. | Developmental neuroscience |
| 15 | 2018 | Bures, Zbynek; Pysanenko, Kateryna; Lindovsky, Jiri; Syka, Josef | Neural plasticity |
| 16 | 2018 | Campolongo, Marcos; Kazlauskas, Nadia; Falasco, German; Urrutia, Leandro; Salgueiro, Natali; Hocht, Christian; Depino, Amaicha Mara | Molecular autism |
| 17 | 2018 | Cintoli, Simona; Cenni, Maria Cristina; Pinto, Bruno; Morea, Silvia; Sale, Alessandro; Maffei, Lamberto; Berardi, Nicoletta | Neural plasticity |
| 18 | 2018 | Cleary-Gaffney, Michael; Coogan, Andrew N. | Physiology & behavior |
| 19 | 2018 | Cope, Elise C.; Opendak, Maya; LaMarca, Elizabeth A.; Murthy, Sahana; Park, Christin Y.; Olson, Lyra B.; Martinez, Susana; Leung, Jacqueline M.; Graham, Andrea L.; Gould, Elizabeth | Hippocampus |
| 20 | 2018 | Cortese, Giuseppe P.; Olin, Andrew; O'Riordan, Kenneth; Hullinger, Rikki; Burger, Corinna | Neurobiology of aging |
| 21 | 2018 | Cutuli, Debora; Berretta, Erica; Caporali, Paola; Sampedro-Piquero, Patricia; De Bartolo, Paola; Laricchiuta, Daniela; Gelfo, Francesca; Pesoli, Matteo; Foti, Francesca; Farioli Vecchioli, Stefano; Petrosini, Laura | Neuropharmacology |
| 22 | 2018 | Dandi, Epsilonvgenia; Kalamari, Aikaterini; Touloumi, Olga; Lagoudaki, Rosa; Nousiopoulou, Evangelia; Simeonidou, Constantina; Spandou, Evangelia; Tata, Despina A. | International journal of developmental neuroscience : the official journal of the International Society for Developmental Neuroscience |
| 23 | 2018 | Faraji, Jamshid; Karimi, Mitra; Soltanpour, Nabiollah; Rouhzadeh, Zahra; Roudaki, Shabnam; Hosseini, S. Abedin; Jafari, S. Yaghoob; Abdollahi, Ali-Akbar; Soltanpour, Nasrin; Moeeini, Reza; Metz, Gerlinde A. S. | Scientific reports |
| 24 | 2018 | Fernandez-Montoya, Julia; Martin, Yasmina B.; Negredo, Pilar; Avendano, Carlos | Brain structure & function |
| 25 | 2018 | Freese, Luana; Almeida, Felipe Borges; Heidrich, Nubia; Hansen, Alana Witt; Steffens, Luiza; Steinmetz, Aline; Moura, Dinara Jaqueline; Gomez, Rosane; Barros, Helena Maria Tannhauser | Pharmacology, biochemistry, and behavior |
| 26 | 2018 | Goncalves, Lara Vezula; Herlinger, Alice Laschuk; Ferreira, Tamara Andrea Alarcon; Coitinho, Juliana Barbosa; Pires, Rita Gomes Wanderley; Martins-Silva, Cristina | Behavioural brain research |
| 27 | 2018 | Gong, Xingrui; Chen, Yongmei; Chang, Jing; Huang, Yue; Cai, Meihau; Zhang, Mazhong | Journal of neuroinflammation |
| 28 | 2018 | Green, Amanda; Esser, Michael J.; Perrot, Tara S. | Behavioural brain research |
| 29 | 2018 | Gregoire, Catherine-Alexandra; Tobin, Stephanie; Goldenstein, Brianna L.; Samarut, Eric; Leclerc, Andreanne; Aumont, Anne; Drapeau, Pierre; Fulton, Stephanie; Fernandes, Karl J. L. | Frontiers in molecular neuroscience |
| 30 | 2018 | Grimm, Jeffrey W.; Glueck, Edwin; Ginder, Darren; Hyde, Jeff; North, Katherine; Jiganti, Kyle | Scientific reports |
| 31 | 2018 | Grinan-Ferre, Christian; Izquierdo, Vanesa; Otero, Eduard; Puigoriol-Illamola, Dolors; Corpas, Ruben; Sanfeliu, Coral; Ortuno-Sahagun, Daniel; Pallas, Merce | Frontiers in cellular neuroscience |
| 32 | 2018 | Hakon, Jakob; Quattromani, Miriana Jlenia; Sjolund, Carin; Tomasevic, Gregor; Carey, Leeanne; Lee, Jin-Moo; Ruscher, Karsten; Wieloch, Tadeusz; Bauer, Adam Q. | NeuroImage. Clinical |
| 33 | 2018 | Hase, Yoshiki; Craggs, Lucinda; Hase, Mai; Stevenson, William; Slade, Janet; Chen, Aiqing; Liang, Di; Ennaceur, Abdel; Oakley, Arthur; Ihara, Masafumi; Horsburgh, Karen; Kalaria, Raj N. | Journal of cerebral blood flow and metabolism : official journal of the International Society of Cerebral Blood Flow and Metabolism |
| 34 | 2018 | Hernan, Amanda E.; Mahoney, J. Matthew; Curry, Willie; Richard, Greg; Lucas, Marcella M.; Massey, Andrew; Holmes, Gregory L.; Scott, Rod C. | PloS one |
| 35 | 2018 | Hofford, Rebecca S.; Prendergast, Mark A.; Bardo, Michael T. | Behavioural brain research |
| 36 | 2018 | Jones, Samantha; Neville, Vikki; Higgs, Laura; Paul, Elizabeth S.; Dayan, Peter; Robinson, Emma S. J.; Mendl, Michael | Scientific reports |
| 37 | 2018 | Kentner, Amanda C.; Scalia, Stephanie; Shin, Junyoung; Migliore, Mattia M.; Rondon-Ortiz, Alejandro N. | Psychoneuroendocrinology |
| 38 | 2018 | Kim, Hyun-Wook; Oh, Seunghak; Lee, Seung Hwan; Lee, Sanghoon; Na, Ji-Eun; Lee, Kea Joo; Rhyu, Im Joo | Microscopy research and technique |
| 39 | 2018 | Lajud, Naima; Diaz-Chavez, Arturo; Radabaugh, Hannah L.; Cheng, Jeffrey P.; Rojo-Soto, Georgina; Valdez-Alarcon, Juan J.; Bondi, Corina O.; Kline, Anthony E. | Journal of neurotrauma |
| 40 | 2018 | Lewis, M. H.; Lindenmaier, Z.; Boswell, K.; Edington, G.; King, M. A.; Muehlmann, A. M. | Genes, brain, and behavior |
| 41 | 2018 | Lopes, Danielle A.; Souza, Thaissa M. O.; de Andrade, Jose S.; Silva, Mariana F. S.; Antunes, Hanna K. M.; Sueur-Maluf, Luciana Le; Cespedes, Isabel C.; Viana, Milena B. | Behavioural brain research |
| 42 | 2018 | Matsuda, Wakoto; Ehara, Ayuka; Nakadate, Kazuhiko; Yoshimoto, Kanji; Ueda, Shuichi | Congenital anomalies |
| 43 | 2018 | McQuaid, Robyn Jane; Dunn, Roderick; Jacobson-Pick, Shlomit; Anisman, Hymie; Audet, Marie-Claude | Frontiers in behavioral neuroscience |
| 44 | 2018 | Moraes, Michele M.; Rabelo, Patricia C. R.; Pinto, Valeria A.; Pires, Washington; Wanner, Samuel P.; Szawka, Raphael E.; Soares, Danusa D. | Neuroscience letters |
| 45 | 2018 | Murueta-Goyena, Ane; Ortuzar, Naiara; Gargiulo, Pascual Angel; Lafuente, Jose Vicente; Bengoetxea, Harkaitz | Molecular neurobiology |
| 46 | 2018 | Nawaz, Amber; Batool, Zehra; Shazad, Sidrah; Rafiq, Sahar; Afzal, Asia; Haider, Saida | Life sciences |
| 47 | 2018 | Neal, Steven; Kent, Molly; Bardi, Massimo; Lambert, Kelly G. | Frontiers in behavioral neuroscience |
| 48 | 2018 | Park, Esther; Tjia, Michelle; Zuo, Yi; Chen, Lu | The Journal of neuroscience : the official journal of the Society for Neuroscience |
| 49 | 2018 | Preston, Kerry E.; Corwin, Rebecca L.; Bader, Julia O.; Crimmins, Stephen L. | Physiology & behavior |
| 50 | 2018 | Prounis, George S.; Thomas, Kyle; Ophir, Alexander G. | The Journal of comparative neurology |
| 51 | 2018 | Pysanenko, Kateryna; Bures, Zbynek; Lindovsky, Jiri; Syka, Josef | Neuroscience |
| 52 | 2018 | Qi, Fangfang; Zuo, Zejie; Hu, Saisai; Xia, Yucen; Song, Dan; Kong, Jiechen; Yang, Yang; Wu, Yingying; Wang, Xiao; Yang, Junhua; Hu, Dandan; Yuan, Qunfang; Zou, Juntao; Guo, Kaihua; Xu, Jie; Yao, Zhibin | Brain, behavior, and immunity |
| 53 | 2018 | Qian, Hai-Zhou; Zhang, Hong; Yin, Lin-Ling; Zhang, Jun-Jian | Current medical science |
| 54 | 2018 | Quattromani, Miriana Jlenia; Pruvost, Mathilde; Guerreiro, Carla; Backlund, Fredrik; Englund, Elisabet; Aspberg, Anders; Jaworski, Tomasz; Hakon, Jakob; Ruscher, Karsten; Kaczmarek, Leszek; Vivien, Denis; Wieloch, Tadeusz | Molecular neurobiology |
| 55 | 2018 | Rae, Mariana; Zanos, Panos; Georgiou, Polymnia; Chivers, Priti; Bailey, Alexis; Camarini, Rosana | Neuropharmacology |
| 56 | 2018 | Rapley, Susan A.; Prickett, Timothy C. R.; Dalrymple-Alford, John C.; Espiner, Eric A. | Frontiers in behavioral neuroscience |
| 57 | 2018 | Requejo, C.; Ruiz-Ortega, J. A.; Cepeda, H.; Sharma, A.; Sharma, H. S.; Ozkizilcik, A.; Tian, R.; Moessler, H.; Ugedo, L.; Lafuente, J. V. | Molecular neurobiology |
| 58 | 2018 | Rico-Barrio, Irantzu; Penasco, Sara; Puente, Nagore; Ramos, Almudena; Fontaine, Christine J.; Reguero, Leire; Giordano, Maria Elvira; Buceta, Ianire; Terradillos, Itziar; Lekunberri, Leire; Mendizabal-Zubiaga, Juan; Rodriguez de Fonseca, Fernando; Gerrikagoitia, Inmaculada; Elezgarai, Izaskun; Grandes, Pedro | Addiction biology |
| 59 | 2018 | Rodriguez-Ortega, Elisa; de la Fuente, Leticia; de Amo, Enedina; Cubero, Inmaculada | Frontiers in behavioral neuroscience |
| 60 | 2018 | Rountree-Harrison, Darius; Burton, Thomas J.; Leamey, Catherine A.; Sawatari, Atomu | Frontiers in behavioral neuroscience |
| 61 | 2018 | Sampedro-Piquero, P.; Alvarez-Suarez, P.; Moreno-Fernandez, R. D.; Garcia-Castro, G.; Cuesta, M.; Begega, A. | Neuroscience |
| 62 | 2018 | Scala, Federico; Nenov, Miroslav N.; Crofton, Elizabeth J.; Singh, Aditya K.; Folorunso, Oluwarotimi; Zhang, Yafang; Chesson, Brent C.; Wildburger, Norelle C.; James, Thomas F.; Alshammari, Musaad A.; Alshammari, Tahani K.; Elfrink, Hannah; Grassi, Claudio; Kasper, James M.; Smith, Ashley E.; Hommel, Jonathan D.; Lichti, Cheryl F.; Rudra, Jai S.; D'Ascenzo, Marcello; Green, Thomas A.; Laezza, Fernanda | Cell reports |
| 63 | 2018 | Seo, Jung Hwa; Pyo, Soonil; Shin, Yoon-Kyum; Nam, Bae-Geun; Kang, Jeong Won; Kim, Kwang Pyo; Lee, Hoo Young; Cho, Sung-Rae | Frontiers in neurology |
| 64 | 2018 | Seong, Ho-Hyun; Park, Jong-Min; Kim, Youn-Jung | Biological research for nursing |
| 65 | 2018 | Shin, Samuel S.; Krishnan, Vijai; Stokes, William; Robertson, Courtney; Celnik, Pablo; Chen, Yanrong; Song, Xiaolei; Lu, Hanzhang; Liu, Peiying; Pelled, Galit | Brain stimulation |
| 66 | 2018 | Sikora, Magdalena; Nicolas, Celine; Istin, Marine; Jaafari, Nematollah; Thiriet, Nathalie; Solinas, Marcello | Behavioural brain research |
| 67 | 2018 | Sikora, Magdalena; Nicolas, Celine; Istin, Marine; Jaafari, Nematollah; Thiriet, Nathalie; Solinas, Marcello | Behavioural brain research |
| 68 | 2018 | Smith, Bryon M.; Yao, Xinyue; Chen, Kelly S.; Kirby, Elizabeth D. | Frontiers in aging neuroscience |
| 69 | 2018 | Sparling, Jessica E.; Baker, Stephanie L.; Bielajew, Catherine | Behavioural brain research |
| 70 | 2018 | Suemaru, Katsuya; Yoshikawa, Misato; Aso, Hiroaki; Watanabe, Masahiko | Epilepsy & behavior : E&B |
| 71 | 2018 | Tarasova, A. Yu; Perepelkina, O. V.; Lil'p, I. G.; Revishchin, A. V.; Pavlova, G. V.; Poletaeva, I. I. | Bulletin of experimental biology and medicine |
| 72 | 2018 | Ulrich, K.; Spriggs, M. J.; Abraham, W. C.; Dalrymple-Alford, J. C.; McNaughton, N. | Hippocampus |
| 73 | 2018 | Wang, Ruixiang; Hausknecht, Kathryn A.; Haj-Dahmane, Samir; Shen, Roh-Yu; Richards, Jerry B. | Behavioural brain research |
| 74 | 2018 | Wassouf, Zinah; Hentrich, Thomas; Samer, Sebastian; Rotermund, Carola; Kahle, Philipp J.; Ehrlich, Ingrid; Riess, Olaf; Casadei, Nicolas; Schulze-Hentrich, Julia M. | Frontiers in cellular neuroscience |
| 75 | 2018 | Wi, Soohyun; Lee, Jang Woo; Kim, MinGi; Park, Chang-Hwan; Cho, Sung-Rae | Cell transplantation |
| 76 | 2018 | Wu, Xiaoying; Liu, Shengqun; Hu, Zhenhua; Zhu, Guosong; Zheng, Gaifang; Wang, Guangzhi | Brain research bulletin |
| 77 | 2018 | Yamada, Jun; Nadanaka, Satomi; Kitagawa, Hiroshi; Takeuchi, Kosei; Jinno, Shozo | The Journal of neuroscience : the official journal of the Society for Neuroscience |
| 78 | 2018 | Zarif, Hadi; Hosseiny, Salma; Paquet, Agnes; Lebrigand, Kevin; Arguel, Marie-Jeanne; Cazareth, Julie; Lazzari, Anne; Heurteaux, Catherine; Glaichenhaus, Nicolas; Chabry, Joelle; Guyon, Alice; Petit-Paitel, Agnes | Frontiers in synaptic neuroscience |
| 79 | 2018 | Zarif, Hadi; Nicolas, Sarah; Guyot, Melanie; Hosseiny, Salma; Lazzari, Anne; Canali, Maria Magdalena; Cazareth, Julie; Brau, Frederic; Golzne, Valentine; Dourneau, Elisa; Maillaut, Maud; Luci, Carmelo; Paquet, Agnes; Lebrigand, Kevin; Arguel, Marie-Jeanne; Daoudlarian, Douglas; Heurteaux, Catherine; Glaichenhaus, Nicolas; Chabry, Joelle; Guyon, Alice; Petit-Paitel, Agnes | Brain, behavior, and immunity |
| 80 | 2018 | Zhang, Tie-Yuan; Keown, Christopher L.; Wen, Xianglan; Li, Junhao; Vousden, Dulcie A.; Anacker, Christoph; Bhattacharyya, Urvashi; Ryan, Richard; Diorio, Josie; O'Toole, Nicholas; Lerch, Jason P.; Mukamel, Eran A.; Meaney, Michael J. | Nature communications |
| 81 | 2018 | Zhang, Yujing; Xu, Dan; Qi, Hong; Yuan, Yin; Liu, Hong; Yao, Shanglong; Yuan, Shiying; Zhang, Jiancheng | Brain research |
| 82 | 2018 | Ziegler-Waldkirch, Stephanie; d'Errico, Paolo; Sauer, Jonas-Frederic; Erny, Daniel; Savanthrapadian, Shakuntala; Loreth, Desiree; Katzmarski, Natalie; Blank, Thomas; Bartos, Marlene; Prinz, Marco; Meyer-Luehmann, Melanie | The EMBO journal |
| 83 | 2018 | Ziegler-Waldkirch, Stephanie; Marksteiner, Karin; Stoll, Johannes; d'Errico, Paolo; Friesen, Marina; Eiler, Denise; Neudel, Lea; Sturn, Verena; Opper, Isabel; Datta, Moumita; Prinz, Marco; Meyer-Luehmann, Melanie | Acta neuropathologica communications |
| 84 | 2017 | Ahmadalipour, Ali; Sadeghzadeh, Jafar; Samaei, Seyed Afshin; Rashidy-Pour, Ali | Basic and clinical neuroscience |
| 85 | 2017 | Ali, Mohamad; Cholvin, Thibault; Muller, Marc Antoine; Cosquer, Brigitte; Kelche, Christian; Cassel, Jean-Christophe; Pereira de Vasconcelos, Anne | Neurobiology of learning and memory |
| 86 | 2017 | Arai, Mitsunori D.; Zhan, Bo; Maruyama, Atsuko; Matsui-Harada, Akiko; Horinouchi, Kazuhiro; Komai, Shoji | Journal of the neurological sciences |
| 87 | 2017 | Aranda, Marcos L.; Gonzalez Fleitas, Maria F.; Dieguez, Hernan H.; Milne, Georgia A.; Devouassoux, Julian D.; Keller Sarmiento, Maria I.; Chianelli, Monica; Sande, Pablo H.; Dorfman, Damian; Rosenstein, Ruth E. | Neuropharmacology |
| 88 | 2017 | Aranda, Marcos L.; Gonzalez Fleitas, Maria F.; Dieguez, Hernan H.; Milne, Georgia A.; Devouassoux, Julian D.; Keller Sarmiento, Maria I.; Chianelli, Monica; Sande, Pablo H.; Dorfman, Damian; Rosenstein, Ruth E. | Neuropharmacology |
| 89 | 2017 | Bahi, Amine | Progress in neuro-psychopharmacology & biological psychiatry |
| 90 | 2017 | Baroncelli, Laura; Cenni, Maria Cristina; Melani, Riccardo; Deidda, Gabriele; Landi, Silvia; Narducci, Roberta; Cancedda, Laura; Maffei, Lamberto; Berardi, Nicoletta | Neuropharmacology |
| 91 | 2017 | Bechard, Allison R.; Bliznyuk, Nikolay; Lewis, Mark H. | Developmental psychobiology |
| 92 | 2017 | Bento-Torres, J.; Sobral, L. L.; Reis, R. R.; de Oliveira, R. B.; Anthony, D. C.; Vasconcelos, P. F. C.; Picanco Diniz, Cristovam Wanderley | Oxidative medicine and cellular longevity |
| 93 | 2017 | Bilkey, David K.; Cheyne, Kirsten R.; Eckert, Michael J.; Lu, Xiaodong; Chowdhury, Shoaib; Worley, Paul F.; Crandall, James E.; Abraham, Wickliffe C. | Hippocampus |
| 94 | 2017 | Brenneis, Christian; Westhof, Andreas; Holschbach, Jeannine; Michaelis, Martin; Guehring, Hans; Kleinschmidt-Doerr, Kerstin | Journal of the American Association for Laboratory Animal Science : JAALAS |
| 95 | 2017 | Buschler, Arne; Manahan-Vaughan, Denise | Neuropharmacology |
| 96 | 2017 | Campelo, Clarissa L. C.; Santos, Jose R.; Silva, Anatildes F.; Dierschnabel, Aline L.; Pontes, Andre; Cavalcante, Jeferson S.; Ribeiro, Alessandra M.; Silva, Regina H. | Behavioural brain research |
| 97 | 2017 | Cao, Min; Pu, Tinglin; Wang, Linmei; Marshall, Charles; He, Hongliang; Hu, Gang; Xiao, Ming | Brain, behavior, and immunity |
| 98 | 2017 | Cao, Wen-Yu; Hu, Zhao-Lan; Xu, Yang; Zhang, Wen-Juan; Huang, Fu-Lian; Qiao, Xiao-Qing; Cui, Yan-Hui; Wan, Wei; Wang, Xue-Qin; Liu, Dan; Dai, Ru-Ping; Li, Fang; Li, Chang-Qi | Psychopharmacology |
| 99 | 2017 | Catalao, Carlos Henrique Rocha; Shimizu, Glaucia Yuri; Tida, Jacqueline Atsuko; Garcia, Camila Araujo Bernardino; Dos Santos, Antonio Carlos; Salmon, Carlos Ernesto Garrido; Rocha, Maria Jose Alves; da Silva Lopes, Luiza | Child's nervous system : ChNS : official journal of the International Society for Pediatric Neurosurgery |
| 100 | 2017 | Chan, Jackie N.-M.; Lee, Jada C.-D.; Lee, Sylvia S. P.; Hui, Katy K. Y.; Chan, Alan H. L.; Fung, Timothy K.-H.; Sanchez-Vidana, Dalinda I.; Lau, Benson W.-M.; Ngai, Shirley P.-C. | Frontiers in behavioral neuroscience |
| 101 | 2017 | Chan, Wilson; Singh, Sanmeet; Keshav, Taj; Dewan, Ramita; Eberly, Christian; Maurer, Robert; Nunez-Parra, Alexia; Araneda, Ricardo C. | Frontiers in synaptic neuroscience |
| 102 | 2017 | Chen, Jia-Yi; Yu, Yuan; Yuan, Yin; Zhang, Yu-Jing; Fan, Xue-Peng; Yuan, Shi-Ying; Zhang, Jian-Cheng; Yao, Shang-Long | Cell death discovery |
| 103 | 2017 | Chen, Xiuping; Zhang, Xin; Liao, Weijing; Wan, Qi | Neurochemical research |
| 104 | 2017 | Chen, Xiuping; Zhang, Xin; Xue, Li; Hao, Chizi; Liao, Weijing; Wan, Qi | Cellular physiology and biochemistry : international journal of experimental cellular physiology, biochemistry, and pharmacology |
| 105 | 2017 | Corcoba, Alberto; Gruetter, Rolf; Do, Kim Q.; Duarte, Joao M. N. | Journal of neurochemistry |
| 106 | 2017 | Cox, Conor D.; Palmer, Linda C.; Pham, Danielle T.; Trieu, Brian H.; Gall, Christine M.; Lynch, Gary | Learning & memory (Cold Spring Harbor, N.Y.) |
| 107 | 2017 | de la Tremblaye, Patricia B.; Bondi, Corina O.; Lajud, Naima; Cheng, Jeffrey P.; Radabaugh, Hannah L.; Kline, Anthony E. | Journal of neurotrauma |
| 108 | 2017 | de la Tremblaye, Patricia B.; Wellcome, Jody L.; de Witt, Benjamin Wells; Cheng, Jeffrey P.; Skidmore, Elizabeth R.; Bondi, Corina O.; Kline, Anthony E. | Neurorehabilitation and neural repair |
| 109 | 2017 | Dolivo, Vassilissa; Taborsky, Michael | Journal of comparative psychology (Washington, D.C. : 1983) |
| 110 | 2017 | Du, Lai-Ling; Wang, Lin; Yang, Xi-Fei; Wang, Ping; Li, Xiao-Hong; Chai, Da-Min; Liu, Bing-Jin; Cao, Yun; Xu, Wei-Qi; Liu, Rong; Tian, Qing; Wang, Jian-Zhi; Zhou, Xin-Wen | Molecular neurobiology |
| 111 | 2017 | Folweiler, Kaitlin A.; Bondi, Corina O.; Ogunsanya, Elizabeth A.; LaPorte, Megan J.; Leary, Jacob B.; Radabaugh, Hannah L.; Monaco, Christina M.; Kline, Anthony E. | Journal of neurotrauma |
| 112 | 2017 | Garcia, Erik J.; Haddon, Tara N.; Saucier, Donald A.; Cain, Mary E. | Pharmacology, biochemistry, and behavior |
| 113 | 2017 | Gauthier, Jamie M.; Lin, Amy; Nic Dhonnchadha, Brid A.; Spealman, Roger D.; Man, Heng-Ye; Kantak, Kathleen M. | Addiction biology |
| 114 | 2017 | Glueck, Edwin; Ginder, Darren; Hyde, Jeff; North, Katherine; Grimm, Jeffrey W. | Psychopharmacology |
| 115 | 2017 | Gomez, Carmen; Redolat, Rosa; Carrasco, Carmen | Pharmacological reports : PR |
| 116 | 2017 | Griva, Myrsini; Lagoudaki, Rosa; Touloumi, Olga; Nousiopoulou, Evangelia; Karalis, Filippos; Georgiou, Thomas; Kokaraki, Georgia; Simeonidou, Constantina; Tata, Despina A.; Spandou, Evangelia | Brain research |
| 117 | 2017 | Gualtieri, Fabio; Bregere, Catherine; Laws, Grace C.; Armstrong, Elena A.; Wylie, Nicholas J.; Moxham, Theo T.; Guzman, Raphael; Boswell, Timothy; Smulders, Tom V. | Frontiers in neuroscience |
| 118 | 2017 | Gurfein, Blake T.; Hasdemir, Burcu; Milush, Jeffrey M.; Touma, Chadi; Palme, Rupert; Nixon, Douglas F.; Darcel, Nicholas; Hecht, Frederick M.; Bhargava, Aditi | PloS one |
| 119 | 2017 | Hajheidari, Samira; Miladi-Gorji, Hossein; Bigdeli, Imanollah | Iranian journal of psychiatry |
| 120 | 2017 | Hammami-Abrand Abadi, Arezoo; Miladi-Gorji, Hossein | Canadian journal of physiology and pharmacology |
| 121 | 2017 | Hase, Yoshiki; Craggs, Lucinda; Hase, Mai; Stevenson, William; Slade, Janet; Lopez, Dianne; Mehta, Rubin; Chen, Aiqing; Liang, Di; Oakley, Arthur; Ihara, Masafumi; Horsburgh, Karen; Kalaria, Raj N. | Journal of neuroinflammation |
| 122 | 2017 | He, Chuan; Tsipis, Constantinos P.; LaManna, Joseph C.; Xu, Kui | Advances in experimental medicine and biology |
| 123 | 2017 | Hegde, Preethi; O'Mara, Shane; Laxmi, Thenkanidiyoor Rao | Annals of neurosciences |
| 124 | 2017 | Hilal, Muna L.; Moreau, Maite M.; Racca, Claudia; Pinheiro, Vera L.; Piguel, Nicolas H.; Santoni, Marie-Josee; Dos Santos Carvalho, Steve; Blanc, Jean-Michel; Abada, Yah-Se K.; Peyroutou, Ronan; Medina, Chantal; Doat, Helene; Papouin, Thomas; Vuillard, Laurent; Borg, Jean-Paul; Rachel, Rivka; Panatier, Aude; Montcouquiol, Mireille; Oliet, Stephane H. R.; Sans, Nathalie | Cerebral cortex (New York, N.Y. : 1991) |
| 125 | 2017 | Hofford, Rebecca S.; Chow, Jonathan J.; Beckmann, Joshua S.; Bardo, Michael T. | Psychopharmacology |
| 126 | 2017 | Holgate, Joan Y.; Garcia, Hilary; Chatterjee, Susmita; Bartlett, Selena E. | Brain and behavior |
| 127 | 2017 | Huttenrauch, Melanie; Walter, Susanne; Kaufmann, Margie; Weggen, Sascha; Wirths, Oliver | Molecular neurobiology |
| 128 | 2017 | Jamal, Imran; Kumar, Vipendra; Vatsa, Naman; Singh, Brijesh Kumar; Shekhar, Shashi; Sharma, Ankit; Jana, Nihar Ranjan | Molecular neurobiology |
| 129 | 2017 | Ji, Mu-Huo; Tang, Hui; Luo, Dan; Qiu, Li-Li; Jia, Min; Yuan, Hong-Mei; Feng, Shan-Wu; Yang, Jian-Jun | Oncotarget |
| 130 | 2017 | Ji, Mu-Huo; Wang, Zhong-Yun; Sun, Xiao-Ru; Tang, Hui; Zhang, Hui; Jia, Min; Qiu, Li-Li; Zhang, Guang-Fen; Peng, Yong G.; Yang, Jian-Jun | Molecular neurobiology |
| 131 | 2017 | Jin, Xinhao; Li, Tao; Zhang, Lina; Ma, Jingxi; Yu, Lehua; Li, Changqing; Niu, Lingchuan | Medical science monitor : international medical journal of experimental and clinical research |
| 132 | 2017 | Jungling, Adel; Reglodi, Dora; Karadi, Zsofia Nozomi; Horvath, Gabor; Farkas, Jozsef; Gaszner, Balazs; Tamas, Andrea | International journal of molecular sciences |
| 133 | 2017 | Keiner, Silke; Niv, Fanny; Neumann, Susanne; Steinbach, Tanja; Schmeer, Christian; Hornung, Katrin; Schlenker, Yvonne; Forster, Martin; Witte, Otto W.; Redecker, Christoph | BMC neuroscience |
| 134 | 2017 | Kim, Wha Young; Cho, Bo Ram; Kwak, Myung Ji; Kim, Jeong-Hoon | Scientific reports |
| 135 | 2017 | Kimura, Louise Faggionato; Mattaraia, Vania Gomes de Moura; Picolo, Gisele | Behavioural brain research |
| 136 | 2017 | Klein, C.; Schreyer, S.; Kohrs, F. E.; Elhamoury, P.; Pfeffer, A.; Munder, T.; Steiner, B. | Scientific reports |
| 137 | 2017 | Klein, C.; Schreyer, S.; Kohrs, F. E.; Elhamoury, P.; Pfeffer, A.; Munder, T.; Steiner, B. | Scientific reports |
| 138 | 2017 | Lafragette, Audrey; Bardo, Michael T.; Lardeux, Virginie; Solinas, Marcello; Thiriet, Nathalie | The international journal of neuropsychopharmacology |
| 139 | 2017 | Leary, Jacob B.; Bondi, Corina O.; LaPorte, Megan J.; Carlson, Lauren J.; Radabaugh, Hannah L.; Cheng, Jeffrey P.; Kline, Anthony E. | Journal of neurotrauma |
| 140 | 2017 | Levine, Jared N.; Chen, Hui; Gu, Yu; Cang, Jianhua | The Journal of neuroscience : the official journal of the Society for Neuroscience |
| 141 | 2017 | Liu, Cong; Gu, Jing-Yang; Han, Jin-Hong; Yan, Fu-Lin; Li, Yan; Lv, Ting-Ting; Zhao, Li-Qin; Shao, Qiu-Jing; Feng, Yan-Yan; Zhang, Xiang-Yang; Wang, Chang-Hong | Brain research bulletin |
| 142 | 2017 | Liu, Xixia; Qiu, Jianhua; Alcon, Sasha; Hashim, Jumana; Meehan, William P. 3rd; Mannix, Rebekah | Journal of neurotrauma |
| 143 | 2017 | Lu, Cheng-Qiu; Zhong, Le; Yan, Chong-Huai; Tian, Ying; Shen, Xiao-Ming | Neuroscience letters |
| 144 | 2017 | Madhavadas, S.; Subramanian, S.; Kutty, B. M. | Physiology international |
| 145 | 2017 | Marianno, Priscila; Abrahao, Karina Possa; Camarini, Rosana | PloS one |
| 146 | 2017 | Marmol, F.; Sanchez, J.; Torres, M. N.; Chamizo, V. D. | Behavioural processes |
| 147 | 2017 | Meireles, Andre L. F.; Marques, Marilia R.; Segabinazi, Ethiane; Spindler, Christiano; Piazza, Francele V.; Salvalaggio, Gabriela S.; Augustin, Otavio A.; Achaval, Matilde; Marcuzzo, Simone | Brain research bulletin |
| 148 | 2017 | Melani, Riccardo; Chelini, Gabriele; Cenni, Maria Cristina; Berardi, Nicoletta | Neuroscience |
| 149 | 2017 | Mileva, Guergana R.; Moyes, Carinna; Syed, Shaezeen; Bielajew, Catherine | Neuropsychobiology |
| 150 | 2017 | Montes, Sergio; Solis-Guillen, Rocio Del Carmen; Garcia-Jacome, David; Paez-Martinez, Nayeli | Neurotoxicology and teratology |
| 151 | 2017 | Muhammad, Mustapha Shehu; Magaji, Rabiu Abdussalam; Mohammed, Aliyu; Isa, Ahmed-Sherif; Magaji, Mohammed Garba | Metabolic brain disease |
| 152 | 2017 | Novaes, Leonardo S.; Dos Santos, Nilton Barreto; Batalhote, Rafaela F. P.; Malta, Marilia Brinati; Camarini, Rosana; Scavone, Cristoforo; Munhoz, Carolina Demarchi | Neuropharmacology |
| 153 | 2017 | Park, Jong-Min; Seong, Ho-Hyun; Jin, Han-Byeol; Kim, Youn-Jung | Biological research for nursing |
| 154 | 2017 | Pasquarelli, Noemi; Voehringer, Patrizia; Henke, Julia; Ferger, Boris | Behavioural brain research |
| 155 | 2017 | Pautassi, Ricardo Marcos; Suarez, Andrea B.; Hoffmann, Lucas Barbosa; Rueda, Andre Veloso; Rae, Mariana; Marianno, Priscila; Camarini, Rosana | Scientific reports |
| 156 | 2017 | Pereira-Caixeta, Ana Raquel; Guarnieri, Leonardo O.; Pena, Roberta R.; Dias, Thomaz L.; Pereira, Grace Schenatto | Molecular neurobiology |
| 157 | 2017 | Pinto, Hyorrana Priscila Pereira; Carvalho, Vinicius Rezende; Medeiros, Daniel de Castro; Almeida, Ana Flavia Santos; Mendes, Eduardo Mazoni Andrade Marcal; Moraes, Marcio Flavio Dutra | Neuroscience |
| 158 | 2017 | Pooriamehr, Alireza; Sabahi, Parviz; Miladi-Gorji, Hossein | Neuroscience letters |
| 159 | 2017 | Qi, Fangfang; Zuo, Zejie; Yang, Junhua; Hu, Saisai; Yang, Yang; Yuan, Qunfang; Zou, Juntao; Guo, Kaihua; Yao, Zhibin | Journal of neuroinflammation |
| 160 | 2017 | Radabaugh, Hannah L.; LaPorte, Megan J.; Greene, Anna M.; Bondi, Corina O.; Lajud, Naima; Kline, Anthony E. | Experimental neurology |
| 161 | 2017 | Rogers, Jake; Li, Shanshan; Lanfumey, Laurence; Hannan, Anthony J.; Renoir, Thibault | Behavioural brain research |
| 162 | 2017 | Ross, Amy P.; Norvelle, Alisa; Choi, Dennis C.; Walton, James C.; Albers, H. Elliott; Huhman, Kim L. | Physiology & behavior |
| 163 | 2017 | Ruitenberg, Marc J.; Wells, Julia; Bartlett, Perry F.; Harvey, Alan R.; Vukovic, Jana | Brain research bulletin |
| 164 | 2017 | Sakalem, Marna Eliana; Seidenbecher, Thomas; Zhang, Mingyue; Saffari, Roja; Kravchenko, Mykola; Wordemann, Stephanie; Diederich, Kai; Schwamborn, Jens C.; Zhang, Weiqi; Ambree, Oliver | Hippocampus |
| 165 | 2017 | Sakata, Kazuko; Overacre, Abigail E. | Journal of neurochemistry |
| 166 | 2017 | Salmin, Vladimir V.; Komleva, Yulia K.; Kuvacheva, Natalia V.; Morgun, Andrey V.; Khilazheva, Elena D.; Lopatina, Olga L.; Pozhilenkova, Elena A.; Shapovalov, Konstantin A.; Uspenskaya, Yulia A.; Salmina, Alla B. | Frontiers in aging neuroscience |
| 167 | 2017 | Selvi, Yavuz; Gergerlioglu, Hasan Serdar; Akbaba, Nursel; Oz, Mehmet; Kandeger, Ali; Demir, Enver Ahmet; Yerlikaya, Fatma Humeyra; Nurullahoglu-Atalik, Kismet Esra | Acta neuropsychiatrica |
| 168 | 2017 | Shilpa, B. M.; Bhagya, V.; Harish, G.; Srinivas Bharath, M. M.; Shankaranarayana Rao, B. S. | Progress in neuro-psychopharmacology & biological psychiatry |
| 169 | 2017 | Shojaei, Amir; Anaraki, Afsaneh Kamali; Mirnajafi-Zadeh, Javad; Atapour, Nafiseh | International journal of developmental neuroscience : the official journal of the International Society for Developmental Neuroscience |
| 170 | 2017 | Soares, Roberto O.; Horiquini-Barbosa, Everton; Almeida, Sebastiao S.; Lachat, Joao-Jose | Behavioural brain research |
| 171 | 2017 | Sta Maria, Naomi S.; Reger, Maxine L.; Cai, Yan; Baquing, Mary Anne T.; Buen, Floyd; Ponnaluri, Aditya; Hovda, David A.; Harris, Neil G.; Giza, Christopher C. | Journal of neurotrauma |
| 172 | 2017 | Stamenkovic, V.; Milenkovic, I.; Galjak, N.; Todorovic, V.; Andjus, P. | Behavioural brain research |
| 173 | 2017 | Stamenkovic, Vera; Stamenkovic, Stefan; Jaworski, Tomasz; Gawlak, Maciej; Jovanovic, Milos; Jakovcevski, Igor; Wilczynski, Grzegorz M.; Kaczmarek, Leszek; Schachner, Melitta; Radenovic, Lidija; Andjus, Pavle R. | Brain structure & function |
| 174 | 2017 | Stuart, Kimberley E.; King, Anna E.; Fernandez-Martos, Carmen M.; Dittmann, Justin; Summers, Mathew J.; Vickers, James C. | The Journal of comparative neurology |
| 175 | 2017 | Tanaka, Mika; Wang, Xiaowen; Mikoshiba, Katsuhiko; Hirase, Hajime; Shinohara, Yoshiaki | The Journal of physiology |
| 176 | 2017 | Vrinda, Marigowda; Sasidharan, Arun; Aparna, Sahajan; Srikumar, Bettadapura N.; Kutty, Bindu M.; Shankaranarayana Rao, Byrathnahalli S. | Epilepsia |
| 177 | 2017 | Wang, Maya Zhe; Marshall, Andrew T.; Kirkpatrick, Kimberly | Behavioural brain research |
| 178 | 2017 | Wang, Qi; Shen, Feng-Yan; Zou, Rong; Zheng, Jing-Jing; Yu, Xiang; Wang, Ying-Wei | Molecular brain |
| 179 | 2017 | Westenbroek, Christel; Perry, Adam N.; Jagannathan, Lakshmikripa; Becker, Jill B. | Physiology & behavior |
| 180 | 2017 | Whitehouse, Cristina M.; Curry-Pochy, Lisa S.; Shafer, Robin; Rudy, Joseph; Lewis, Mark H. | Behavioural brain research |
| 181 | 2017 | Wohr, Markus; Engelhardt, K. Alexander; Seffer, Dominik; Sungur, A. Ozge; Schwarting, Rainer K. W. | Current topics in behavioral neurosciences |
| 182 | 2017 | Yagishita, Kaori; Suzuki, Ritsuko; Mizuno, Shota; Katoh-Semba, Ritsuko; Sadakata, Tetsushi; Sano, Yoshitake; Furuichi, Teiichi; Shinoda, Yo | Neuroscience letters |
| 183 | 2017 | Yamaguchi, Hiroshi; Hara, Yuta; Ago, Yukio; Takano, Erika; Hasebe, Shigeru; Nakazawa, Takanobu; Hashimoto, Hitoshi; Matsuda, Toshio; Takuma, Kazuhiro | Behavioural brain research |
| 184 | 2017 | Yeshurun, Shlomo; Short, Annabel K.; Bredy, Timothy W.; Pang, Terence Y.; Hannan, Anthony J. | Psychoneuroendocrinology |
| 185 | 2017 | Zhang, Xin; Chen, Xiu-Ping; Lin, Jun-Bin; Xiong, Yu; Liao, Wei-Jing; Wan, Qi | Brain research |
| 186 | 2017 | Zheng, Jie; Jiang, Ying-Ying; Xu, Ling-Chi; Ma, Long-Yu; Liu, Feng-Yu; Cui, Shuang; Cai, Jie; Liao, Fei-Fei; Wan, You; Yi, Ming | The Journal of neuroscience : the official journal of the Society for Neuroscience |
| 187 | 2017 | Zhou, Zhike; Liu, Tingting; Sun, Xiaoyu; Mu, Xiaopeng; Zhu, Gang; Xiao, Ting; Zhao, Mei; Zhao, Chuansheng | Behavioural brain research |
| 188 | 2016 | Alwis, Dasuni Sathsara; Yan, Edwin Bingbing; Johnstone, Victoria; Carron, Simone; Hellewell, Sarah; Morganti-Kossmann, Maria Cristina; Rajan, Ramesh | Journal of neurotrauma |
| 189 | 2016 | Ashokan, Archana; Hegde, Akshaya; Mitra, Rupshi | Psychoneuroendocrinology |
| 190 | 2016 | Barbosa, Everton Horiquini; Soares, Roberto Oliveira; Braga, Natalia Nassif; Almeida, Sebastiao de Sousa; Lachat, Joao-Jose | Nutritional neuroscience |
| 191 | 2016 | Bardi, M.; Kaufman, C.; Franssen, C.; Hyer, M. M.; Rzucidlo, A.; Brown, M.; Tschirhart, M.; Lambert, K. G. | Journal of neuroendocrinology |
| 192 | 2016 | Baroncelli, Laura; Scali, Manuela; Sansevero, Gabriele; Olimpico, Francesco; Manno, Ilaria; Costa, Mario; Sale, Alessandro | The Journal of neuroscience : the official journal of the Society for Neuroscience |
| 193 | 2016 | Bechard, Allison R.; Cacodcar, Nadia; King, Michael A.; Lewis, Mark H. | Behavioural brain research |
| 194 | 2016 | Berardo, Luciana R.; Fabio, Maria C.; Pautassi, Ricardo M. | Frontiers in behavioral neuroscience |
| 195 | 2016 | Borniger, Jeremy C.; Cisse, Yasmine M.; Cantemir-Stone, Carmen Z.; Bolon, Brad; Nelson, Randy J.; Marsh, Clay B. | Behavioural brain research |
| 196 | 2016 | Botanas, Chrislean Jun; Lee, Hyelim; de la Pena, June Bryan; Dela Pena, Irene Joy; Woo, Taeseon; Kim, Hee Jin; Han, Doug Hyun; Kim, Bung-Nyun; Cheong, Jae Hoon | Physiology & behavior |
| 197 | 2016 | Brenes, Juan C.; Lackinger, Martin; Hoglinger, Gunter U.; Schratt, Gerhard; Schwarting, Rainer K. W.; Wohr, Markus | The Journal of comparative neurology |
| 198 | 2016 | Browne, Caleb J.; Fletcher, Paul J.; Zeeb, Fiona D. | Psychopharmacology |
| 199 | 2016 | Buschert, Jens; Sakalem, Marna E.; Saffari, Roja; Hohoff, Christa; Rothermundt, Matthias; Arolt, Volker; Zhang, Weiqi; Ambree, Oliver | Progress in neuro-psychopharmacology & biological psychiatry |
| 200 | 2016 | Catanzaro, G.; Pucci, M.; Viscomi, M. T.; Lanuti, M.; Feole, M.; Angeletti, S.; Grasselli, G.; Mandolesi, G.; Bari, M.; Centonze, D.; D'Addario, C.; Maccarrone, M. | Journal of neuroimmunology |
| 201 | 2016 | Catuara-Solarz, Silvina; Espinosa-Carrasco, Jose; Erb, Ionas; Langohr, Klaus; Gonzalez, Juan Ramon; Notredame, Cedric; Dierssen, Mara | eNeuro |
| 202 | 2016 | Chamizo, V. D.; Rodriguez, C. A.; Sanchez, J.; Marmol, F. | Learning & behavior |
| 203 | 2016 | Cho, Sung-Rae; Suh, Hwal; Yu, Ji Hea; Kim, Hyongbum Henry; Seo, Jung Hwa; Seo, Cheong Hoon | International journal of molecular sciences |
| 204 | 2016 | Cordner, Z. A.; Tamashiro, K. L. K. | Translational psychiatry |
| 205 | 2016 | Dezsi, Gabi; Ozturk, Ezgi; Salzberg, Michael R.; Morris, Margaret; O'Brien, Terence J.; Jones, Nigel C. | Neurobiology of disease |
| 206 | 2016 | Diaz, Ramiro; Miguel, Patricia Maidana; Deniz, Bruna Ferrary; Confortim, Heloisa Deola; Barbosa, Silvia; Mendonca, Monique Culturato Padilha; da Cruz-Hofling, Maria Alice; Pereira, Lenir Orlandi | International journal of developmental neuroscience : the official journal of the International Society for Developmental Neuroscience |
| 207 | 2016 | Diniz, Daniel Guerreiro; de Oliveira, Marcus Augusto; de Lima, Camila Mendes; Foro, Cesar Augusto Raiol; Sosthenes, Marcia Consentino Kronka; Bento-Torres, Joao; da Costa Vasconcelos, Pedro Fernando; Anthony, Daniel Clive; Diniz, Cristovam Wanderley Picanco | Behavioral and brain functions : BBF |
| 208 | 2016 | Dow-Edwards, Diana; Frank, Ashley; Wade, Dean; Weedon, Jeremy; Izenwasser, Sari | Neurotoxicology and teratology |
| 209 | 2016 | Fan, Dan; Li, Jun; Zheng, Bin; Hua, Lei; Zuo, Zhiyi | Molecular neurobiology |
| 210 | 2016 | Foglesong, Grant D.; Huang, Wei; Liu, Xianglan; Slater, Andrew M.; Siu, Jason; Yildiz, Vedat; Salton, Stephen R. J.; Cao, Lei | Endocrinology |
| 211 | 2016 | Fuchs, Fanny; Cosquer, Brigitte; Penazzi, Lorene; Mathis, Chantal; Kelche, Christian; Majchrzak, Monique; Barbelivien, Alexandra | Behavioural brain research |
| 212 | 2016 | Fureix, Carole; Walker, Michael; Harper, Laura; Reynolds, Kathryn; Saldivia-Woo, Amanda; Mason, Georgia | Behavioural brain research |
| 213 | 2016 | Gapp, Katharina; Bohacek, Johannes; Grossmann, Jonas; Brunner, Andrea M.; Manuella, Francesca; Nanni, Paolo; Mansuy, Isabelle M. | Neuropsychopharmacology : official publication of the American College of Neuropsychopharmacology |
| 214 | 2016 | Garbugino, Luciana; Centofante, Eleonora; D'Amato, Francesca R. | Neural plasticity |
| 215 | 2016 | Garthe, Alexander; Roeder, Ingo; Kempermann, Gerd | Hippocampus |
| 216 | 2016 | Gelfo, Francesca; Florenzano, Fulvio; Foti, Francesca; Burello, Lorena; Petrosini, Laura; De Bartolo, Paola | Brain structure & function |
| 217 | 2016 | Gergerlioglu, Hasan Serdar; Oz, Mehmet; Demir, Enver Ahmet; Nurullahoglu-Atalik, Kismet Esra; Yerlikaya, Fatma Humeyra | Life sciences |
| 218 | 2016 | Greifzu, Franziska; Kalogeraki, Evgenia; Lowel, Siegrid | Neurobiology of aging |
| 219 | 2016 | Grimm, Jeffrey W.; Barnes, Jesse L.; Koerber, Jonathon; Glueck, Edwin; Ginder, Darren; Hyde, Jeff; Eaton, Laura | Brain structure & function |
| 220 | 2016 | Grinan-Ferre, Christian; Perez-Caceres, David; Gutierrez-Zetina, Sofia Martinez; Camins, Antoni; Palomera-Avalos, Veronica; Ortuno-Sahagun, Daniel; Rodrigo, M. Teresa; Pallas, M. | Molecular neurobiology |
| 221 | 2016 | Grinan-Ferre, Christian; Puigoriol-Illamola, Dolors; Palomera-Avalos, Veronica; Perez-Caceres, David; Companys-Alemany, Julia; Camins, Antonio; Ortuno-Sahagun, Daniel; Rodrigo, M. Teresa; Pallas, Merce | Frontiers in aging neuroscience |
| 222 | 2016 | Hammami-Abrand Abadi, Arezoo; Miladi-Gorji, Hossein; Bigdeli, Imanollah | Behavioural pharmacology |
| 223 | 2016 | Hendershott, Taylor R.; Cronin, Marie E.; Langella, Stephanie; McGuinness, Patrick S.; Basu, Alo C. | Behavioural brain research |
| 224 | 2016 | Hilario, Willyan Franco; Herlinger, Alice Laschuk; Areal, Lorena Bianchine; de Moraes, Livia Silveira; Ferreira, Tamara Andrea Alarcon; Andrade, Tassiane Emanuelle Servane; Martins-Silva, Cristina; Pires, Rita Gomes Wanderley | Journal of molecular neuroscience : MN |
| 225 | 2016 | Hofford, Rebecca S.; Beckmann, Joshua S.; Bardo, Michael T. | Drug and alcohol dependence |
| 226 | 2016 | Hong, S. Lee; Estrada-Sanchez, Ana Maria; Barton, Scott J.; Rebec, George V. | Behavioural brain research |
| 227 | 2016 | Huttenrauch, Melanie; Salinas, Gabriela; Wirths, Oliver | Frontiers in molecular neuroscience |
| 228 | 2016 | Islas-Preciado, D.; Lopez-Rubalcava, C.; Gonzalez-Olvera, J.; Gallardo-Tenorio, A.; Estrada-Camarena, E. | Neuroscience |
| 229 | 2016 | Jha, S.; Dong, B. E.; Xue, Y.; Delotterie, D. F.; Vail, M. G.; Sakata, K. | Translational psychiatry |
| 230 | 2016 | Jiang, Congyu; Yu, Kewei; Wu, Yi; Xie, Hongyu; Liu, Gang; Wu, Junfa; Jia, Jie; Kuang, Shenyi | Journal of stroke and cerebrovascular diseases : the official journal of National Stroke Association |
| 231 | 2016 | Kamakura, Remi; Kovalainen, Miia; Leppaluoto, Juhani; Herzig, Karl-Heinz; Makela, Kari A. | Physiological reports |
| 232 | 2016 | Kang, Hee; Choi, Dong-Hee; Kim, Su-Kang; Lee, Jongmin; Kim, Youn-Jung | Developmental neuroscience |
| 233 | 2016 | Kapgal, Vijayakumar; Prem, Neethi; Hegde, Preethi; Laxmi, T. R.; Kutty, Bindu M. | Neurobiology of learning and memory |
| 234 | 2016 | Kentner, Amanda C.; Khoury, Antoine; Lima Queiroz, Erika; MacRae, Molly | Brain, behavior, and immunity |
| 235 | 2016 | Kim, Myung-Sun; Yu, Ji Hea; Kim, Chul Hoon; Choi, Jae Yong; Seo, Jung Hwa; Lee, Min-Young; Yi, Chi Hoon; Choi, Tae Hyun; Ryu, Young Hoon; Lee, Jong Eun; Lee, Bae Hwan; Kim, Hyongbum; Cho, Sung-Rae | Journal of cerebral blood flow and metabolism : official journal of the International Society of Cerebral Blood Flow and Metabolism |
| 236 | 2016 | Koe, A. S.; Ashokan, A.; Mitra, R. | Translational psychiatry |
| 237 | 2016 | Kondo, Hiroko; Kurahashi, Minori; Mori, Daisuke; Iinuma, Mitsuo; Tamura, Yasuo; Mizutani, Kenmei; Shimpo, Kan; Sonoda, Shigeru; Azuma, Kagaku; Kubo, Kin-ya | Archives of oral biology |
| 238 | 2016 | Kondo, Mari A.; Gray, Laura J.; Pelka, Gregory J.; Leang, Sook-Kwan; Christodoulou, John; Tam, Patrick P. L.; Hannan, Anthony J. | Developmental neurobiology |
| 239 | 2016 | Kreilaus, Fabian; Spiro, Adena S.; Hannan, Anthony J.; Garner, Brett; Jenner, Andrew M. | Journal of Huntington's disease |
| 240 | 2016 | Lach, Gilliard; Bicca, Maira Assuncao; Hoeller, Alexandre Ademar; Santos, Evelyn Cristina da Silva; Costa, Ana Paula Ramos; de Lima, Thereza Christina Monteiro | Neuropeptides |
| 241 | 2016 | Lambert, Kelly; Hyer, Molly; Bardi, Massimo; Rzucidlo, Amanda; Scott, Samantha; Terhune-Cotter, Brennan; Hazelgrove, Ashley; Silva, Ilton; Kinsley, Craig | Neuroscience |
| 242 | 2016 | Lauterborn, Julie C.; Palmer, Linda C.; Jia, Yousheng; Pham, Danielle T.; Hou, Bowen; Wang, Weisheng; Trieu, Brian H.; Cox, Conor D.; Kantorovich, Svetlana; Gall, Christine M.; Lynch, Gary | The Journal of neuroscience : the official journal of the Society for Neuroscience |
| 243 | 2016 | Li, Ki Angel; Lund, Emilie Torp; Voigt, Jorg-Peter W. | Behavioural processes |
| 244 | 2016 | Li, Yu-Wang; Li, Qing-Yun; Wang, Jin-Hua; Xu, Xiao-Lin | Cellular physiology and biochemistry : international journal of experimental cellular physiology, biochemistry, and pharmacology |
| 245 | 2016 | Link, Andrea S.; Kurinna, Svitlana; Havlicek, Steven; Lehnert, Sandra; Reichel, Martin; Kornhuber, Johannes; Winner, Beate; Huth, Tobias; Zheng, Fang; Werner, Sabine; Alzheimer, Christian | Molecular neurobiology |
| 246 | 2016 | Lopez-Luengo, Beatriz; Muela-Martinez, Jose A. | Cognitive neuropsychiatry |
| 247 | 2016 | Ma, Yao-Ying; Wang, Xiusong; Huang, Yanhua; Marie, Helene; Nestler, Eric J.; Schluter, Oliver M.; Dong, Yan | Proceedings of the National Academy of Sciences of the United States of America |
| 248 | 2016 | Mahati, K.; Bhagya, V.; Christofer, T.; Sneha, A.; Shankaranarayana Rao, B. S. | Neurobiology of learning and memory |
| 249 | 2016 | Makowska, I. Joanna; Weary, Daniel M. | PloS one |
| 250 | 2016 | McCreary, J. Keiko; Erickson, Zachary T.; Hao, YongXin; Ilnytskyy, Yaroslav; Kovalchuk, Igor; Metz, Gerlinde A. S. | Scientific reports |
| 251 | 2016 | McCreary, J. Keiko; Erickson, Zachary T.; Metz, Gerlinde A. S. | Neuroscience letters |
| 252 | 2016 | Mesa-Gresa, Patricia; Ramos-Campos, Marta; Redolat, Rosa | Physiology & behavior |
| 253 | 2016 | Meyers, Emily A.; Gobeske, Kevin T.; Bond, Allison M.; Jarrett, Jennifer C.; Peng, Chian-Yu; Kessler, John A. | Neurobiology of aging |
| 254 | 2016 | Molina, S. J.; Capani, F.; Guelman, L. R. | Brain research |
| 255 | 2016 | Morozova, Anna; Zubkov, Eugene; Strekalova, Tatyana; Kekelidze, Zurab; Storozeva, Zinaida; Schroeter, Careen A.; Bazhenova, Nataliia; Lesch, Klaus-Peter; Cline, Brandon H.; Chekhonin, Vladimir | Progress in neuro-psychopharmacology & biological psychiatry |
| 256 | 2016 | Mustroph, M. L.; Pinardo, H.; Merritt, J. R.; Rhodes, J. S. | Behavioural brain research |
| 257 | 2016 | Neidl, Romain; Schneider, Anne; Bousiges, Olivier; Majchrzak, Monique; Barbelivien, Alexandra; de Vasconcelos, Anne Pereira; Dorgans, Kevin; Doussau, Frederic; Loeffler, Jean-Philippe; Cassel, Jean-Christophe; Boutillier, Anne-Laurence | The Journal of neuroscience : the official journal of the Society for Neuroscience |
| 258 | 2016 | Nobre, Manoel Jorge | International journal of developmental neuroscience : the official journal of the International Society for Developmental Neuroscience |
| 259 | 2016 | Ortiz-Perez, A.; Espinosa-Raya, J.; Picazo, O. | Cognitive processing |
| 260 | 2016 | Perez-Martin, Margarita; Rivera, Patricia; Blanco, Eduardo; Lorefice, Clara; Decara, Juan; Pavon, Francisco J.; Serrano, Antonia; Rodriguez de Fonseca, Fernando; Suarez, Juan | Frontiers in neuroscience |
| 261 | 2016 | Pusic, Kae M.; Pusic, Aya D.; Kraig, Richard P. | Cellular and molecular neurobiology |
| 262 | 2016 | Radabaugh, Hannah L.; Carlson, Lauren J.; O'Neil, Darik A.; LaPorte, Megan J.; Monaco, Christina M.; Cheng, Jeffrey P.; de la Tremblaye, Patricia B.; Lajud, Naima; Bondi, Corina O.; Kline, Anthony E. | Experimental neurology |
| 263 | 2016 | Rahati, M.; Nozari, M.; Eslami, H.; Shabani, M.; Basiri, M. | Neuroscience |
| 264 | 2016 | Rahmeier, Francine Luciano; Zavalhia, Lisiane Silveira; Tortorelli, Lucas Silva; Huf, Fernanda; Gea, Luiza Paul; Meurer, Rosalva Thereza; Machado, Aryadne Cardoso; Gomez, Rosane; Fernandes, Marilda da Cruz | Neuroscience letters |
| 265 | 2016 | Reichmann, Florian; Wegerer, Vanessa; Jain, Piyush; Mayerhofer, Raphaela; Hassan, Ahmed M.; Frohlich, Esther E.; Bock, Elisabeth; Pritz, Elisabeth; Herzog, Herbert; Holzer, Peter; Leitinger, Gerd | Scientific reports |
| 266 | 2016 | Rodriguez, Carlos I.; Magcalas, Christy M.; Barto, Daniel; Fink, Brandi C.; Rice, James P.; Bird, Clark W.; Davies, Suzy; Pentkowski, Nathan S.; Savage, Daniel D.; Hamilton, Derek A. | Behavioural brain research |
| 267 | 2016 | Rogers, J.; Vo, U.; Buret, L. S.; Pang, T. Y.; Meiklejohn, H.; Zeleznikow-Johnston, A.; Churilov, L.; van den Buuse, M.; Hannan, A. J.; Renoir, T. | Translational psychiatry |
| 268 | 2016 | Salois, Garrick; Smith, Jeffrey S. | Neural plasticity |
| 269 | 2016 | Sampedro-Piquero, P.; Castilla-Ortega, E.; Zancada-Menendez, C.; Santin, L. J.; Begega, A. | Neuroscience |
| 270 | 2016 | Santos, Camila Mauricio; Peres, Fernanda Fiel; Diana, Mariana Cepollaro; Justi, Veronica; Suiama, Mayra Akimi; Santana, Marcela Goncalves; Abilio, Vanessa Costhek | Schizophrenia research |
| 271 | 2016 | Schuch, Clarissa Pedrini; Diaz, Ramiro; Deckmann, Iohanna; Rojas, Joseane Jimenez; Deniz, Bruna Ferrary; Pereira, Lenir Orlandi | Neuroscience letters |
| 272 | 2016 | Schuch, Clarissa Pedrini; Jeffers, Matthew Strider; Antonescu, Sabina; Nguemeni, Carine; Gomez-Smith, Mariana; Pereira, Lenir Orlandi; Morshead, Cindi M.; Corbett, Dale | Behavioural brain research |
| 273 | 2016 | Scichilone, John M.; Yarraguntla, Kalyan; Charalambides, Ana; Harney, Jacob P.; Butler, David | Journal of molecular neuroscience : MN |
| 274 | 2016 | Slaker, Megan; Barnes, Jesse; Sorg, Barbara A.; Grimm, Jeffrey W. | PloS one |
| 275 | 2016 | Stein, Liana R.; O'Dell, Kazuko A.; Funatsu, Michiyo; Zorumski, Charles F.; Izumi, Yukitoshi | Neuroscience |
| 276 | 2016 | Sun, Xiao R.; Zhang, Hui; Zhao, Hong T.; Ji, Mu H.; Li, Hui H.; Wu, Jing; Li, Kuan Y.; Yang, Jian J. | Behavioural brain research |
| 277 | 2016 | Thanos, Panayotis K.; Hamilton, John; O'Rourke, Joseph R.; Napoli, Anthony; Febo, Marcelo; Volkow, Nora D.; Blum, Kenneth; Gold, Mark | Oncotarget |
| 278 | 2016 | Tomiga, Yuki; Ito, Ai; Sudo, Mizuki; Ando, Soichi; Maruyama, Akino; Nakashima, Shihoko; Kawanaka, Kentaro; Uehara, Yoshinari; Kiyonaga, Akira; Tanaka, Hiroaki; Higaki, Yasuki | Biochemical and biophysical research communications |
| 279 | 2016 | Vega-Rivera, Nelly Maritza; Ortiz-Lopez, Leonardo; Gomez-Sanchez, Ariadna; Oikawa-Sala, Julian; Estrada-Camarena, Erika Monserrat; Ramirez-Rodriguez, Gerardo Bernabe | Behavioural brain research |
| 280 | 2016 | Wang, Xin; Chen, Aiguo; Wu, Honghai; Ye, Min; Cheng, Hong; Jiang, Xinfeng; Wang, Xiaohong; Zhang, Xiaobin; Wu, Di; Gu, Xin; Shen, Feiyang; Shan, Chunlei; Yu, Duonan | Brain research |
| 281 | 2016 | Weaver, S. R.; Cronick, C. M.; Prichard, A. P.; Laporta, J.; Benevenga, N. J.; Hernandez, L. L. | Laboratory animals |
| 282 | 2016 | Weil, Zachary M.; Karelina, Kate; Gaier, Kristopher R.; Corrigan, Timothy E. D.; Corrigan, John D. | Journal of neurotrauma |
| 283 | 2016 | Whitaker, Julia W.; Moy, Sheryl S.; Pritchett-Corning, Kathleen R.; Fletcher, Craig A. | Journal of the American Association for Laboratory Animal Science : JAALAS |
| 284 | 2016 | Winocur, Gordon; Wojtowicz, J. Martin; Merkley, Christina M.; Tannock, Ian F. | Behavioral neuroscience |
| 285 | 2016 | Wren-Dail, Melissa A.; Dauchy, Robert T.; Ooms, Tara G.; Baker, Kate C.; Blask, David E.; Hill, Steven M.; Dupepe, Lynell M.; Bohm, Rudolf P. Jr | Comparative medicine |
| 286 | 2016 | Wu, Yufeng; Gan, Yu; Yuan, Hui; Wang, Qing; Fan, Yingchao; Li, Guohua; Zhang, Jian; Yao, Ming; Gu, Jianren; Tu, Hong | Biochemical and biophysical research communications |
| 287 | 2016 | Xiao, Run; Bergin, Stephen M.; Huang, Wei; Slater, Andrew M.; Liu, Xianglan; Judd, Ryan T.; Lin, En-Ju D.; Widstrom, Kyle J.; Scoville, Steven D.; Yu, Jianhua; Caligiuri, Michael A.; Cao, Lei | Cancer immunology research |
| 288 | 2016 | Xing, Renzhong; Zhang, Yanling; Xu, Hua; Luo, Xiaobin; Chang, Raymond Chuen-Chung; Liu, Jianjun; Yang, Xifei | Oncotarget |
| 289 | 2016 | Xu, Huixin; Gelyana, Eilrayna; Rajsombath, Molly; Yang, Ting; Li, Shaomin; Selkoe, Dennis | The Journal of neuroscience : the official journal of the Society for Neuroscience |
| 290 | 2016 | Yang, Meng; Ozturk, Ezgi; Salzberg, Michael R.; Rees, Sandra; Morris, Margaret; O'Brien, Terence J.; Jones, Nigel C. | Epilepsia |
| 291 | 2016 | Zerwas, Meike; Trouche, Stephanie; Richetin, Kevin; Escude, Timothe; Halley, Helene; Gerardy-Schahn, Rita; Verret, Laure; Rampon, Claire | Brain structure & function |
| 292 | 2016 | Zhang, Mingqiang; Wu, Jing; Huo, Lan; Luo, Liang; Song, Xi; Fan, Fei; Lu, Yiming; Liang, Dong | Molecular neurobiology |
| 293 | 2016 | Zhang, Xiao Qian; Mu, Jing Wei; Wang, Hui Bin; Jolkkonen, Jukka; Liu, Ting Ting; Xiao, Ting; Zhao, Mei; Zhang, Chao Dong; Zhao, Chuan Sheng | Molecular medicine reports |
| 294 | 2016 | Zhang, Yafang; Crofton, Elizabeth J.; Fan, Xiuzhen; Li, Dingge; Kong, Fanping; Sinha, Mala; Luxon, Bruce A.; Spratt, Heidi M.; Lichti, Cheryl F.; Green, Thomas A. | Neuroscience |
| 295 | 2016 | Zhang, Yafang; Kong, Fanping; Crofton, Elizabeth J.; Dragosljvich, Steven N.; Sinha, Mala; Li, Dingge; Fan, Xiuzhen; Koshy, Shyny; Hommel, Jonathan D.; Spratt, Heidi M.; Luxon, Bruce A.; Green, Thomas A. | Frontiers in molecular neuroscience |
| 296 | 2016 | Zou, Chengyu; Shi, Yuan; Ohli, Jasmin; Schuller, Ulrich; Dorostkar, Mario M.; Herms, Jochen | Acta neuropathologica |
| 297 | 2016 | Zuena, Anna Rita; Zinni, Manuela; Giuli, Chiara; Cinque, Carlo; Alema, Giovanni Sebastiano; Giuliani, Alessandro; Catalani, Assia; Casolini, Paola; Cozzolino, Roberto | Physiology & behavior |
| 298 | 2015 | Ahmadalipour, A.; Sadeghzadeh, J.; Vafaei, A. A.; Bandegi, A. R.; Mohammadkhani, R.; Rashidy-Pour, A. | Neuroscience |
| 299 | 2015 | Arndt, David L.; Peterson, Christy J.; Cain, Mary E. | PloS one |
| 300 | 2015 | Bayat, Mahnaz; Sharifi, Mohammad Davood; Haghani, Masoud; Shabani, Mohammad | Brain research bulletin |
| 301 | 2015 | Begenisic, Tatjana; Sansevero, Gabriele; Baroncelli, Laura; Cioni, Giovanni; Sale, Alessandro | Neurobiology of disease |
| 302 | 2015 | Bergami, Matteo; Masserdotti, Giacomo; Temprana, Silvio G.; Motori, Elisa; Eriksson, Therese M.; Gobel, Jana; Yang, Sung Min; Conzelmann, Karl-Klaus; Schinder, Alejandro F.; Gotz, Magdalena; Berninger, Benedikt | Neuron |
| 303 | 2015 | Burrows, Emma L.; McOmish, Caitlin E.; Buret, Laetitia S.; Van den Buuse, Maarten; Hannan, Anthony J. | Neuropsychopharmacology : official publication of the American College of Neuropsychopharmacology |
| 304 | 2015 | Caporali, Paola; Cutuli, Debora; Gelfo, Francesca; Laricchiuta, Daniela; Foti, Francesca; De Bartolo, Paola; Angelucci, Francesco; Petrosini, Laura | Frontiers in behavioral neuroscience |
| 305 | 2015 | Cardenas, Lorena; Garcia-Garcia, Fabio; Santiago-Roque, Isela; Martinez, Armando J.; Coria-Avila, Genaro A.; Corona-Morales, Aleph A. | International journal of developmental neuroscience : the official journal of the International Society for Developmental Neuroscience |
| 306 | 2015 | Catuara-Solarz, Silvina; Espinosa-Carrasco, Jose; Erb, Ionas; Langohr, Klaus; Notredame, Cedric; Gonzalez, Juan R.; Dierssen, Mara | Frontiers in behavioral neuroscience |
| 307 | 2015 | Chabry, Joelle; Nicolas, Sarah; Cazareth, Julie; Murris, Emilie; Guyon, Alice; Glaichenhaus, Nicolas; Heurteaux, Catherine; Petit-Paitel, Agnes | Brain, behavior, and immunity |
| 308 | 2015 | Clemenson, Gregory D.; Lee, Star W.; Deng, Wei; Barrera, Vanessa R.; Iwamoto, Kei S.; Fanselow, Michael S.; Gage, Fred H. | Hippocampus |
| 309 | 2015 | Clipperton-Allen, Amy E.; Ingrao, Joelle C.; Ruggiero, Laura; Batista, Lucas; Ovari, Jelena; Hammermueller, Jutta; Armstrong, John N.; Bienzle, Dorothee; Choleris, Elena; Turner, Patricia V. | Journal of the American Association for Laboratory Animal Science : JAALAS |
| 310 | 2015 | Connors, E. J.; Migliore, M. M.; Pillsbury, S. L.; Shaik, A. N.; Kentner, A. C. | Psychoneuroendocrinology |
| 311 | 2015 | Cunha, A. O. S.; de Oliveira, J. A. C.; Almeida, S. S.; Garcia-Cairasco, N.; Leao, R. M. | Neuroscience |
| 312 | 2015 | Cutuli, Debora; Caporali, Paola; Gelfo, Francesca; Angelucci, Francesco; Laricchiuta, Daniela; Foti, Francesca; De Bartolo, Paola; Bisicchia, Elisa; Molinari, Marco; Farioli Vecchioli, Stefano; Petrosini, Laura | Frontiers in behavioral neuroscience |
| 313 | 2015 | Darna, Mahesh; Beckmann, Joshua S.; Gipson, Cassandra D.; Bardo, Michael T.; Dwoskin, Linda P. | Brain research |
| 314 | 2015 | De Bartolo, P.; Florenzano, F.; Burello, L.; Gelfo, F.; Petrosini, L. | Brain structure & function |
| 315 | 2015 | de Sousa, Aline A.; Dos Reis, Renata R.; de Lima, Camila M.; de Oliveira, Marcus A.; Fernandes, Taiany N.; Gomes, Giovanni F.; Diniz, Daniel G.; Magalhaes, Nara M.; Diniz, Cristovam G.; Sosthenes, Marcia C. K.; Bento-Torres, Joao; Diniz, Jose Antonio P. Jr; Vasconcelos, Pedro F. da C.; Diniz, Cristovam Wanderley P. | The European journal of neuroscience |
| 316 | 2015 | Deats, Sean P.; Adidharma, Widya; Yan, Lily | Neuroscience letters |
| 317 | 2015 | Dorfman, Damian; Aranda, Marcos L.; Rosenstein, Ruth E. | PloS one |
| 318 | 2015 | Dubreucq, Sarah; Marsicano, Giovanni; Chaouloff, Francis | Behavioural brain research |
| 319 | 2015 | Ewin, Sarah E.; Kangiser, Megan M.; Stairs, Dustin J. | Experimental and clinical psychopharmacology |
| 320 | 2015 | Favre, Monica R.; La Mendola, Deborah; Meystre, Julie; Christodoulou, Dimitri; Cochrane, Melissa J.; Markram, Henry; Markram, Kamila | Frontiers in neuroscience |
| 321 | 2015 | Freymann, Jennifer; Tsai, Ping-Ping; Stelzer, Helge; Hackbarth, Hansjoachim | Lab animal |
| 322 | 2015 | Gajhede Gram, Marie; Gade, Louise; Wogensen, Elise; Mogensen, Jesper; Mala, Hana | Brain research |
| 323 | 2015 | Goes, Tiago Costa; Antunes, Fabricio Dias; Teixeira-Silva, Flavia | Neuroscience letters |
| 324 | 2015 | Gomez, Adrian M.; Altomare, Diego; Sun, Wei-Lun; Midde, Narasimha M.; Ji, Hao; Shtutman, Michael; Turner, Jill R.; Creek, Kim E.; Zhu, Jun | The international journal of neuropsychopharmacology |
| 325 | 2015 | Gornicka-Pawlak, Elzbieta; Janowski, Miroslaw; Jablonska, Anna; Sypecka, Joanna; Domanska-Janik, Krystyna | Behavioural brain research |
| 326 | 2015 | Hajheidari, Samira; Miladi-Gorji, Hossein; Bigdeli, Imanollah | Neuroscience letters |
| 327 | 2015 | Heinla, Indrek; Leidmaa, Este; Kongi, Karina; Pennert, Airi; Innos, Jurgen; Nurk, Kaarel; Tekko, Triin; Singh, Katyayani; Vanaveski, Taavi; Reimets, Riin; Mandel, Merle; Lang, Aavo; Lillevali, Kersti; Kaasik, Allen; Vasar, Eero; Philips, Mari-Anne | Frontiers in neuroscience |
| 328 | 2015 | Hofford, Rebecca S.; Prendergast, Mark A.; Bardo, Michael T. | Behavioural brain research |
| 329 | 2015 | Horvath, Gabor; Kiss, Peter; Nemeth, Jozsef; Lelesz, Beata; Tamas, Andrea; Reglodi, Dora | Neuro endocrinology letters |
| 330 | 2015 | Hosseiny, Salma; Pietri, Mariel; Petit-Paitel, Agnes; Zarif, Hadi; Heurteaux, Catherine; Chabry, Joelle; Guyon, Alice | Brain structure & function |
| 331 | 2015 | Huang, Huang; Wang, Linmei; Cao, Min; Marshall, Charles; Gao, Junying; Xiao, Na; Hu, Gang; Xiao, Ming | The international journal of neuropsychopharmacology |
| 332 | 2015 | Hullinger, Rikki; O'Riordan, Kenneth; Burger, Corinna | Neurobiology of learning and memory |
| 333 | 2015 | Hunter, Amy Silvestri | Neurobiology of learning and memory |
| 334 | 2015 | Huzard, Damien; Mumby, Dave G.; Sandi, Carmen; Poirier, Guillaume L.; van der Kooij, Michael A. | Physiology & behavior |
| 335 | 2015 | Iggena, Deetje; Klein, Charlotte; Garthe, Alexander; Winter, York; Kempermann, Gerd; Steiner, Barbara | Scientific reports |
| 336 | 2015 | Jamal, Amanda L.; Walker, Tara L.; Waber Nguyen, Amanda J.; Berman, Robert F.; Kempermann, Gerd; Waldau, Ben | Cell transplantation |
| 337 | 2015 | Ji, Mu-huo; Wang, Xing-ming; Sun, Xiao-ru; Zhang, Hui; Ju, Ling-sha; Qiu, Li-li; Yang, Jiao-jiao; Jia, Min; Wu, Jing; Yang, Jianjun | Journal of molecular neuroscience : MN |
| 338 | 2015 | Jiang, Cuiping; Xu, Xiaoxiao; Yu, Liping; Xu, Jinghong; Zhang, Jiping | The European journal of neuroscience |
| 339 | 2015 | Kop, Willem J.; Galvao, Tatiana F.; Synowski, Stephen J.; Xu, Wenhong; Can, Adem; O'Shea, Karen M.; Gould, Todd D.; Stanley, William C. | Physiology & behavior |
| 340 | 2015 | Kuptsova, Kristina; Kvist, Elisabet; Nitzsche, Franziska; Jolkkonen, Jukka | Romanian journal of morphology and embryology = Revue roumaine de morphologie et embryologie |
| 341 | 2015 | Lauterborn, Julie C.; Jafari, Matiar; Babayan, Alex H.; Gall, Christine M. | Cerebral cortex (New York, N.Y. : 1991) |
| 342 | 2015 | Leger, Marianne; Paizanis, Eleni; Dzahini, Kwamivi; Quiedeville, Anne; Bouet, Valentine; Cassel, Jean-Christophe; Freret, Thomas; Schumann-Bard, Pascale; Boulouard, Michel | Cerebral cortex (New York, N.Y. : 1991) |
| 343 | 2015 | Levone, Brunno R.; Cryan, John F.; O'Leary, Olivia F. | Neurobiology of stress |
| 344 | 2015 | Li, Xinjuan; Meng, Li; Huang, Keyu; Wang, Hua; Li, Dongliang | Neuroscience letters |
| 345 | 2015 | Lockworth, Cynthia R.; Kim, Sun-Jin; Liu, Jun; Palla, Shana L.; Craig, Suzanne L. | Journal of the American Association for Laboratory Animal Science : JAALAS |
| 346 | 2015 | Longo, A.; Oberto, A.; Mele, P.; Mattiello, L.; Pisu, M. G.; Palanza, P.; Serra, M.; Eva, C. | Genes, brain, and behavior |
| 347 | 2015 | Lopez, Marcelo F.; Laber, Kathy | Physiology & behavior |
| 348 | 2015 | Loss, Cassio Morais; Binder, Luisa Bandeira; Muccini, Eduarda; Martins, Wagner Carbolin; de Oliveira, Paulo Alexandre; Vandresen-Filho, Samuel; Prediger, Rui Daniel; Tasca, Carla Ines; Zimmer, Eduardo R.; Costa-Schmidt, Luiz Ernesto; de Oliveira, Diogo Losch; Viola, Giordano Gubert | Neurobiology of learning and memory |
| 349 | 2015 | Maegele, M.; Braun, M.; Wafaisade, A.; Schafer, N.; Lippert-Gruener, M.; Kreipke, C.; Rafols, J.; Schafer, U.; Angelov, D. N.; Stuermer, E. K. | Physiological research |
| 350 | 2015 | Marmol, Frederic; Rodriguez, Clara A.; Sanchez, Juan; Chamizo, Victoria D. | Brain research |
| 351 | 2015 | Meng, C.; Zhang, J.-C.; Shi, R.-L.; Zhang, S.-H.; Yuan, S.-Y. | Neuroscience |
| 352 | 2015 | Meng, Fan-Tao; Zhao, Jun; Ni, Rong-Jun; Fang, Hui; Zhang, Li-Feng; Zhang, Zhi; Liu, Ya-Jing | Neuro endocrinology letters |
| 353 | 2015 | Meyer, Andrew C.; Bardo, Michael T. | Psychopharmacology |
| 354 | 2015 | Mileva, Guergana R.; Bielajew, Catherine | Behavioural brain research |
| 355 | 2015 | Mora-Gallegos, Andrea; Rojas-Carvajal, Mijail; Salas, Sofia; Saborio-Arce, Adriana; Fornaguera-Trias, Jaime; Brenes, Juan C. | Neurobiology of learning and memory |
| 356 | 2015 | Mosaferi, Belal; Babri, Shirin; Ebrahimi, Hadi; Mohaddes, Gisou | Physiology & behavior |
| 357 | 2015 | Mosaferi, Belal; Babri, Shirin; Mohaddes, Gisou; Khamnei, Saeed; Mesgari, Mehran | International journal of developmental neuroscience : the official journal of the International Society for Developmental Neuroscience |
| 358 | 2015 | Nicolas, Sarah; Veyssiere, Julie; Gandin, Carine; Zsurger, Nicole; Pietri, Mariel; Heurteaux, Catherine; Glaichenhaus, Nicolas; Petit-Paitel, Agnes; Chabry, Joelle | Psychoneuroendocrinology |
| 359 | 2015 | Novkovic, T.; Heumann, R.; Manahan-Vaughan, D. | Neuroscience |
| 360 | 2015 | Novkovic, Tanja; Mittmann, Thomas; Manahan-Vaughan, Denise | Hippocampus |
| 361 | 2015 | Nozari, M.; Shabani, M.; Farhangi, A. M.; Mazhari, S.; Atapour, N. | Neuroscience |
| 362 | 2015 | Nudi, Evan T.; Jacqmain, Justin; Dubbs, Kelsey; Geeck, Katalin; Salois, Garrick; Searles, Madeleine A.; Smith, Jeffrey S. | Journal of neurotrauma |
| 363 | 2015 | Oddi, Diego; Subashi, Enejda; Middei, Silvia; Bellocchio, Luigi; Lemaire-Mayo, Valerie; Guzman, Manuel; Crusio, Wim E.; D'Amato, Francesca R.; Pietropaolo, Susanna | Neuropsychopharmacology : official publication of the American College of Neuropsychopharmacology |
| 364 | 2015 | Onaolapo, Olakunle J.; Onaolapo, Adejoke Y.; Akanmu, Moses A.; Olayiwola, Gbola | Annals of neurosciences |
| 365 | 2015 | Pan-Vazquez, Alejandro; Rye, Natasha; Ameri, Mitra; McSparron, Bethan; Smallwood, Gabriella; Bickerdyke, Jordan; Rathbone, Alex; Dajas-Bailador, Federico; Toledo-Rodriguez, Maria | Molecular brain |
| 366 | 2015 | Pan-Vazquez, Alejandro; Rye, Natasha; Ameri, Mitra; McSparron, Bethan; Smallwood, Gabriella; Bickerdyke, Jordan; Rathbone, Alex; Dajas-Bailador, Federico; Toledo-Rodriguez, Maria | Molecular brain |
| 367 | 2015 | Pardo, Marta; King, Margaret K.; Perez-Costas, Emma; Melendez-Ferro, Miguel; Martinez, Ana; Beurel, Eleonore; Jope, Richard S. | Frontiers in behavioral neuroscience |
| 368 | 2015 | Park, C. Sehwan; Valomon, Amandine; Welzl, Hans | PloS one |
| 369 | 2015 | Pascual, Rodrigo; Valencia, Martina; Bustamante, Carlos | Neuropediatrics |
| 370 | 2015 | Peck, Joshua A.; Galaj, Ewa; Eshak, Stephanie; Newman, Kristena L.; Ranaldi, Robert | Pharmacology, biochemistry, and behavior |
| 371 | 2015 | Possamai, Fernanda; dos Santos, Juliano; Walber, Thais; Marcon, Juliana C.; dos Santos, Tiago Souza; Lino de Oliveira, Cilene | Progress in neuro-psychopharmacology & biological psychiatry |
| 372 | 2015 | Ragu Varman, Durairaj; Rajan, Koilmani Emmanuvel | PloS one |
| 373 | 2015 | Ratajczak, P.; Nowakowska, E.; Kus, K.; Danielewicz, R.; Herman, S.; Wozniak, A. | Human & experimental toxicology |
| 374 | 2015 | Reichmann, Florian; Painsipp, Evelin; Holzer, Peter; Kummer, Daniel; Bock, Elisabeth; Leitinger, Gerd | Journal of neuroscience methods |
| 375 | 2015 | Rojas, J. J.; Deniz, B. F.; Schuch, C. P.; Carletti, J. V.; Deckmann, I.; Diaz, R.; Matte, C.; dos Santos, T. M.; Wyse, A. T.; Netto, C. A.; Pereira, L. O. | Neuroscience |
| 376 | 2015 | Rossetti, Maria F.; Varayoud, Jorgelina; Moreno-Piovano, Guillermo S.; Luque, Enrique H.; Ramos, Jorge G. | Molecular and cellular endocrinology |
| 377 | 2015 | Sampedro-Piquero, P.; Zancada-Menendez, C.; Begega, A. | Neuroscience |
| 378 | 2015 | Scholz, Jan; Allemang-Grand, Rylan; Dazai, Jun; Lerch, Jason P. | NeuroImage |
| 379 | 2015 | Soares, Roberto O.; Rorato, Rodrigo C.; Padovan, Diego; Lachat, Joao-Jose; Antunes-Rodrigues, Jose; Elias, Lucila L. K.; Almeida, Sebastiao S. | Brain research |
| 380 | 2015 | Souchet, Benoit; Latour, Alizee; Gu, Yuchen; Daubigney, Fabrice; Paul, Jean-Louis; Delabar, Jean-Maurice; Janel, Nathalie | Journal of molecular neuroscience : MN |
| 381 | 2015 | Tanas, Lukasz; Ostaszewski, Pawel; Iwan, Anna | Acta neurobiologiae experimentalis |
| 382 | 2015 | Torres-Lista, Virginia; Gimenez-Llort, Lydia | Behavioural processes |
| 383 | 2015 | Valero-Aracama, Maria Jesus; Sauvage, Magdalena M.; Yoshida, Motoharu | Behavioural brain research |
| 384 | 2015 | van der Veen, Rixt; Kentrop, Jiska; van der Tas, Liza; Loi, Manila; van IJzendoorn, Marinus H.; Bakermans-Kranenburg, Marian J.; Joels, Marian | Frontiers in behavioral neuroscience |
| 385 | 2015 | Wadowska, Magdalena; Woods, Julie; Rogozinska, Magdalena; Briones, Teresita L. | Neuropathology and applied neurobiology |
| 386 | 2015 | Wheeler, R. R.; Swan, M. P.; Hickman, D. L. | Laboratory animals |
| 387 | 2015 | Wood, Ruth I.; Knoll, Allison T.; Levitt, Pat | Physiology & behavior |
| 388 | 2015 | Xu, Xu-Feng; Li, Ting; Wang, Dong-Dong; Chen, Bing; Wang, Yue; Chen, Zhe-Yu | Scientific reports |
| 389 | 2015 | Yang, Shu; Lu, Wei; Zhou, De-shan; Tang, Yong | Neuroscience letters |
| 390 | 2015 | Yang, Shu; Lu, Wei; Zhou, De-shan; Tang, Yong | Brain structure & function |
| 391 | 2015 | Young, Jennica; Pionk, Timothy; Hiatt, Ivy; Geeck, Katalin; Smith, Jeffrey S. | Brain research bulletin |
| 392 | 2015 | Zeeni, N.; Bassil, M.; Fromentin, G.; Chaumontet, C.; Darcel, N.; Tome, D.; Daher, C. F. | Physiology & behavior |
| 393 | 2015 | Zhang, M. Q.; Ji, M. H.; Zhao, Q. S.; Jia, M.; Qiu, L. L.; Yang, J. J.; Peng, Y. G.; Yang, J. J.; Martynyuk, A. E. | British journal of anaesthesia |
| 394 | 2015 | Zhang, Wen; Zhang, Fangjun; Han, Yu; Liu, Hui; Wang, Ye; Yue, Bo; Chen, Jun; Chen, Yang; Gao, Ya | Neuroreport |
| 395 | 2015 | Zhang, Xiaoqian; Liu, Tingting; Zhou, Zhike; Mu, Xiaopeng; Song, Chengguang; Xiao, Ting; Zhao, Mei; Zhao, Chuansheng | Journal of molecular neuroscience : MN |
| 396 | 2015 | Zhao, Yupeng; Chen, Kaizheng; Shen, Xia | BioMed research international |
| 397 | 2015 | Zubedat, Salman; Aga-Mizrachi, Shlomit; Cymerblit-Sabba, Adi; Ritter, Ami; Nachmani, Maayan; Avital, Avi | Stress (Amsterdam, Netherlands) |
| 398 | 2014 | Akkerman, Sven; Prickaerts, Jos; Bruder, Ann K.; Wolfs, Kevin H. M.; De Vry, Jochen; Vanmierlo, Tim; Blokland, Arjan | PloS one |
| 399 | 2014 | Alvarez, Paula Steffen; Simao, Fabricio; Hemb, Marta; Xavier, Leder Leal; Nunes, Magda Lahorgue | International journal of developmental neuroscience : the official journal of the International Society for Developmental Neuroscience |
| 400 | 2014 | Anderson, Megan R. | Journal of applied animal welfare science : JAAWS |
| 401 | 2014 | Arndt, David L.; Arnold, Jennifer C.; Cain, Mary E. | Experimental and clinical psychopharmacology |
| 402 | 2014 | Barichello, Tatiana; Fagundes, Glauco D.; Generoso, Jaqueline S.; Dagostin, Caroline S.; Simoes, Lutiana R.; Vilela, Marcia C.; Comim, Clarissa M.; Petronilho, Fabricia; Quevedo, Joao; Teixeira, Antonio L. | Revista brasileira de psiquiatria (Sao Paulo, Brazil : 1999) |
| 403 | 2014 | Barone, Ilaria; Novelli, Elena; Strettoi, Enrica | Molecular vision |
| 404 | 2014 | Bates, M. L. Shawn; Emery, Michael A.; Wellman, Paul J.; Eitan, Shoshana | Drug and alcohol dependence |
| 405 | 2014 | Bayod, S.; Mennella, I.; Sanchez-Roige, S.; Lalanza, J. F.; Escorihuela, R. M.; Camins, A.; Pallas, M.; Canudas, A. M. | Brain research |
| 406 | 2014 | Blazquez, Gloria; Canete, Toni; Tobena, Adolf; Gimenez-Llort, Lydia; Fernandez-Teruel, Alberto | Behavioural brain research |
| 407 | 2014 | Boschen, Karen E.; Hamilton, Gillian F.; Delorme, James E.; Klintsova, Anna Y. | Alcohol (Fayetteville, N.Y.) |
| 408 | 2014 | Bures, Zbynek; Bartosova, Jolana; Lindovsky, Jiri; Chumak, Tetyana; Popelar, Jiri; Syka, Josef | The European journal of neuroscience |
| 409 | 2014 | Caliaperumal, Jayalakshmi; Colbourne, Frederick | Neurorehabilitation and neural repair |
| 410 | 2014 | Campos, R. C.; Parfitt, G. M.; Polese, C. E.; Coutinho-Silva, R.; Morrone, F. B.; Barros, D. M. | Neuroscience |
| 411 | 2014 | Cao, Wenyu; Duan, Juan; Wang, Xueqin; Zhong, Xiaolin; Hu, Zhaolan; Huang, Fulian; Wang, Hongtao; Zhang, Juan; Li, Fang; Zhang, Jianyi; Luo, Xuegang; Li, Chang-Qi | Behavioural brain research |
| 412 | 2014 | Caporali, Paola; Cutuli, Debora; Gelfo, Francesca; Laricchiuta, Daniela; Foti, Francesca; De Bartolo, Paola; Mancini, Laura; Angelucci, Francesco; Petrosini, Laura | Frontiers in behavioral neuroscience |
| 413 | 2014 | Cheng, Liang; Wang, Shao-Hui; Jia, Nan; Xie, Min; Liao, Xiao-Mei | Brain & development |
| 414 | 2014 | Clarke, Jared; Langdon, Kristopher D.; Corbett, Dale | Journal of cerebral blood flow and metabolism : official journal of the International Society of Cerebral Blood Flow and Metabolism |
| 415 | 2014 | Cloutier, Sylvie; Wahl, Kim; Baker, Chelsea; Newberry, Ruth C. | Journal of the American Association for Laboratory Animal Science : JAALAS |
| 416 | 2014 | Connors, E. J.; Shaik, A. N.; Migliore, M. M.; Kentner, A. C. | Brain, behavior, and immunity |
| 417 | 2014 | Coombs, Ellen J. | Alternatives to laboratory animals : ATLA |
| 418 | 2014 | Darcy, Michael J.; Trouche, Stephanie; Jin, Shan-Xue; Feig, Larry A. |  |
| 419 | 2014 | Darwish, Hala; Mahmood, Asim; Schallert, Timothy; Chopp, Michael; Therrien, Barbara | Brain injury |
| 420 | 2014 | Doulames, Vanessa; Lee, Sangmook; Shea, Thomas B. | The International journal of neuroscience |
| 421 | 2014 | Dunkerson, Jacob; Moritz, Kasey E.; Young, Jennica; Pionk, Tim; Fink, Kyle; Rossignol, Julien; Dunbar, Gary; Smith, Jeffrey S. | Restorative neurology and neuroscience |
| 422 | 2014 | Durairaj, Ragu Varman; Koilmani, Emmanuvel Rajan | General and comparative endocrinology |
| 423 | 2014 | Febinger, Heidi Y.; George, Amrita; Priestley, Jill; Toth, Linda A.; Opp, Mark R. | Journal of the American Association for Laboratory Animal Science : JAALAS |
| 424 | 2014 | Ferland, Jacqueline-Marie N.; Zeeb, Fiona D.; Yu, Katrina; Kaur, Sukhbir; Taves, Matthew D.; Winstanley, Catharine A. | The European journal of neuroscience |
| 425 | 2014 | Galeano, Pablo; Blanco, Eduardo; Logica Tornatore, Tamara M. A.; Romero, Juan I.; Holubiec, Mariana I.; Rodriguez de Fonseca, Fernando; Capani, Francisco | Frontiers in behavioral neuroscience |
| 426 | 2014 | Geuzaine, Annabelle; Tirelli, Ezio | Behavioural brain research |
| 427 | 2014 | Gill, Kathryn E.; Chappell, Ann M.; Beveridge, Thomas J. R.; Porrino, Linda J.; Weiner, Jeffrey L. | Alcoholism, clinical and experimental research |
| 428 | 2014 | Gill, Margaret J.; Weiss, Mark L.; Cain, Mary E. | Drug and alcohol dependence |
| 429 | 2014 | Gregoire, Catherine-Alexandra; Bonenfant, David; Le Nguyen, Adalie; Aumont, Anne; Fernandes, Karl J. L. | PloS one |
| 430 | 2014 | Greifzu, Franziska; Pielecka-Fortuna, Justyna; Kalogeraki, Evgenia; Krempler, Katja; Favaro, Plinio D.; Schluter, Oliver M.; Lowel, Siegrid | Proceedings of the National Academy of Sciences of the United States of America |
| 431 | 2014 | Hall, J. M.; Vetreno, R. P.; Savage, L. M. | Neuroscience |
| 432 | 2014 | Hamilton, G. F.; Jablonski, S. A.; Schiffino, F. L.; St Cyr, S. A.; Stanton, M. E.; Klintsova, A. Y. | Neuroscience |
| 433 | 2014 | Hamilton, Kristen R.; Elliott, Brenda M.; Berger, Sarah Shafer; Grunberg, Neil E. | Experimental and clinical psychopharmacology |
| 434 | 2014 | Heinla, Indrek; Leidmaa, Este; Visnapuu, Tanel; Philips, Mari-Anne; Vasar, Eero | Behavioural brain research |
| 435 | 2014 | Hunter, Amy Silvestri | Experimental brain research |
| 436 | 2014 | Ignacio, Cherry; Mooney, Sandra M.; Middleton, Frank A. | Frontiers in pediatrics |
| 437 | 2014 | Irier, Hasan; Street, R. Craig; Dave, Ronak; Lin, Li; Cai, Catherine; Davis, Timothy Hayden; Yao, Bing; Cheng, Ying; Jin, Peng | Genomics |
| 438 | 2014 | Jacqmain, Justin; Nudi, Evan T.; Fluharty, Sarah; Smith, Jeffrey S. | Behavioural brain research |
| 439 | 2014 | Jung, Christian K. E.; Herms, Jochen | Cerebral cortex (New York, N.Y. : 1991) |
| 440 | 2014 | Kalogeraki, Evgenia; Greifzu, Franziska; Haack, Franziska; Lowel, Siegrid | The Journal of neuroscience : the official journal of the Society for Neuroscience |
| 441 | 2014 | Kato, Tomokazu; Eriguchi, Takashi; Fujiwara, Norio; Murata, Yoshihiro; Yoshino, Atsuo; Sakatani, Kaoru; Katayama, Yoichi | Advances in experimental medicine and biology |
| 442 | 2014 | Kawano, Takashi; Morikawa, Akihiro; Imori, Satoko; Waki, Sayaka; Tamura, Takahiko; Yamanaka, Daiki; Yamazaki, Fumimoto; Yokoyama, Masataka | Journal of anesthesia |
| 443 | 2014 | Kim, Hyungjun; Cho, Junghun; Kim, Young R.; Song, Youngkyu; Chun, Song-I.; Suh, Ji-Yeon; Kim, Jeong Kon; Ryu, Yeon-Hee; Choi, Sun-Mi; Cho, Hyungjoon; Cho, Gyunggoo | PloS one |
| 444 | 2014 | Korgan, Austin C.; Green, Amanda D.; Perrot, Tara S.; Esser, Michael J. | Behavioural brain research |
| 445 | 2014 | Lay, Christopher C.; Frostig, Ron D. | The European journal of neuroscience |
| 446 | 2014 | Lestaevel, P.; Airault, F.; Racine, R.; Bensoussan, H.; Dhieux, B.; Delissen, O.; Manens, L.; Aigueperse, J.; Voisin, P.; Souidi, M. | Journal of molecular neuroscience : MN |
| 447 | 2014 | Li, Chang-Qi; Luo, Yan-Wei; Bi, Fang-Fang; Cui, Tao-Tao; Song, Ling; Cao, Wen-Yu; Zhang, Jian-Yi; Li, Fang; Xu, Jun-Mei; Hao, Wei; Xing, Xiao-Wei; Zhou, Fiona H.; Zhou, Xin-Fu; Dai, Ru-Ping | Neuropsychopharmacology : official publication of the American College of Neuropsychopharmacology |
| 448 | 2014 | Lichti, Cheryl F.; Fan, Xiuzhen; English, Robert D.; Zhang, Yafang; Li, Dingge; Kong, Fanping; Sinha, Mala; Andersen, Clark R.; Spratt, Heidi; Luxon, Bruce A.; Green, Thomas A. | Frontiers in behavioral neuroscience |
| 449 | 2014 | Lidfors, L.; Wichman, A.; Ewaldsson, B.; Lindh, A.-S. | Laboratory animals |
| 450 | 2014 | Lima, Aline P. A. S.; Silva, Kelly; Padovan, Claudia Maria; Almeida, Sebastiao Sousa; Fukuda, Marisa Tomoe Hebihara | Behavioural brain research |
| 451 | 2014 | Lima, F. B.; Spinelli de Oliveira, E. | Behavioural processes |
| 452 | 2014 | Livneh, Yoav; Adam, Yoav; Mizrahi, Adi | Neuron |
| 453 | 2014 | Madinier, Alexandre; Quattromani, Miriana Jlenia; Sjolund, Carin; Ruscher, Karsten; Wieloch, Tadeusz | PloS one |
| 454 | 2014 | Marriott, Amber L.; Tasker, R. Andrew; Ryan, Catherine L.; Doucette, Tracy A. | Neuroscience letters |
| 455 | 2014 | Matteucci, Andrea; Ceci, Chiara; Mallozzi, Cinzia; Macri, Simone; Malchiodi-Albedi, Fiorella; Laviola, Giovanni | Experimental eye research |
| 456 | 2014 | Mazarakis, Nektarios K.; Mo, Christina; Renoir, Thibault; van Dellen, Anton; Deacon, Robert; Blakemore, Colin; Hannan, Anthony J. | Journal of Huntington's disease |
| 457 | 2014 | Melik, Enver; Babar, Emine; Kocahan, Sayad; Guven, Mustafa; Akillioglu, Kubra | International journal of developmental neuroscience : the official journal of the International Society for Developmental Neuroscience |
| 458 | 2014 | Mesa-Gresa, Patricia; Ramos-Campos, Marta; Redolat, Rosa | Behavioural processes |
| 459 | 2014 | Monaco, Christina M.; Gebhardt, Kory M.; Chlebowski, Sarah M.; Shaw, Kaitlyn E.; Cheng, Jeffrey P.; Henchir, Jeremy J.; Zupa, Margaret F.; Kline, Anthony E. | Journal of neurotrauma |
| 460 | 2014 | Monteiro, Brisa M. M.; Moreira, Fabricio A.; Massensini, Andre R.; Moraes, Marcio F. D.; Pereira, Grace S. | Hippocampus |
| 461 | 2014 | Morelli, Emanuela; Ghiglieri, Veronica; Pendolino, Valentina; Bagetta, Vincenza; Pignataro, Annabella; Fejtova, Anna; Costa, Cinzia; Ammassari-Teule, Martine; Gundelfinger, Eckart D.; Picconi, Barbara; Calabresi, Paolo | Experimental neurology |
| 462 | 2014 | Morgan, Judith L.; Svenson, Karen L.; Lake, Jeffrey P.; Zhang, Weidong; Stearns, Timothy M.; Marion, Michael A.; Peters, Luanne L.; Paigen, Beverly; Donahue, Leah Rae | PloS one |
| 463 | 2014 | Moritz, Kasey E.; Geeck, Katalin; Underly, Robert G.; Searles, Madeleine; Smith, Jeffrey S. | Restorative neurology and neuroscience |
| 464 | 2014 | Mychasiuk, Richelle; Muhammad, Arif; Kolb, Bryan | Synapse (New York, N.Y.) |
| 465 | 2014 | Nader, Joelle; Claudia, Chauvet; El Rawas, Rana; Favot, Laure; Jaber, Mohamed; Thiriet, Nathalie; Solinas, Marcello | Neuropsychopharmacology : official publication of the American College of Neuropsychopharmacology |
| 466 | 2014 | Nowakowska, Elzbieta; Kus, Krzysztof; Ratajczak, Piotr; Cichocki, Michal; Wozniak, Anna | Pharmacological reports : PR |
| 467 | 2014 | Nozari, Masoumeh; Shabani, Mohammad; Hadadi, Mahdieh; Atapour, Nafiseh | Psychopharmacology |
| 468 | 2014 | O'Connor, Angela M.; Burton, Thomas J.; Leamey, Catherine A.; Sawatari, Atomu |  |
| 469 | 2014 | Palm, Sara; Nylander, Ingrid | Alcoholism, clinical and experimental research |
| 470 | 2014 | Pamidi, N.; Nayak, B. Satheesha; Mohandas, K. G.; Rao, S. Srinivasa; Madhav, N. Venu | Bratislavske lekarske listy |
| 471 | 2014 | Pamidi, Narendra; Nayak, Satheesha | Biomedical journal |
| 472 | 2014 | Piazza, Francele Valente; Segabinazi, Ethiane; Centenaro, Ligia Aline; do Nascimento, Patricia Severo; Achaval, Matilde; Marcuzzo, Simone | Metabolic brain disease |
| 473 | 2014 | Pietropaolo, Susanna; Feldon, Joram; Yee, Benjamin K. | Cognitive, affective & behavioral neuroscience |
| 474 | 2014 | Polito, Letizia; Chierchia, Armando; Tunesi, Marta; Bouybayoune, Ihssane; Kehoe, Patrick Gavin; Albani, Diego; Forloni, Gianluigi | Journal of Alzheimer's disease : JAD |
| 475 | 2014 | Pusic, Aya D.; Kraig, Richard P. | Glia |
| 476 | 2014 | Pusic, Kae M.; Pusic, Aya D.; Kemme, Jordan; Kraig, Richard P. | Glia |
| 477 | 2014 | Quattromani, Miriana Jlenia; Cordeau, Pierre; Ruscher, Karsten; Kriz, Jasna; Wieloch, Tadeusz | Neurobiology of disease |
| 478 | 2014 | Ramirez-Rodriguez, G.; Ocana-Fernandez, M. A.; Vega-Rivera, N. M.; Torres-Perez, O. M.; Gomez-Sanchez, A.; Estrada-Camarena, E.; Ortiz-Lopez, L. | Neuroscience |
| 479 | 2014 | Ravenelle, R.; Santolucito, H. B.; Byrnes, E. M.; Byrnes, J. J.; Donaldson, S. T. | Neuroscience |
| 480 | 2014 | Reshef, Ronen; Kreisel, Tirzah; Beroukhim Kay, Dorsa; Yirmiya, Raz | Brain, behavior, and immunity |
| 481 | 2014 | Sampedro-Piquero, P.; Arias, J. L.; Begega, A. | Experimental gerontology |
| 482 | 2014 | Sampedro-Piquero, P.; Begega, A.; Arias, J. L. | Physiology & behavior |
| 483 | 2014 | Sampedro-Piquero, P.; De Bartolo, Paola; Petrosini, Laura; Zancada-Menendez, C.; Arias, J. L.; Begega, A. | Neurobiology of learning and memory |
| 484 | 2014 | Schaeffer, Evelin L.; Cerulli, Fabiana G.; Souza, Helio O. X.; Catanozi, Sergio; Gattaz, Wagner F. | Journal of neural transmission (Vienna, Austria : 1996) |
| 485 | 2014 | Schneider, Armin; Rogalewski, Andreas; Wafzig, Oliver; Kirsch, Friederike; Gretz, Norbert; Kruger, Carola; Diederich, Kai; Pitzer, Claudia; Laage, Rico; Plaas, Christian; Vogt, Gerhard; Minnerup, Jens; Schabitz, Wolf-Rudiger | Experimental & translational stroke medicine |
| 486 | 2014 | Schreiber, S.; Lin, R.; Haim, L.; Baratz-Goldstien, R.; Rubovitch, V.; Vaisman, N.; Pick, C. G. | Behavioural brain research |
| 487 | 2014 | Sequeira-Cordero, A.; Mora-Gallegos, A.; Cuenca-Berger, P.; Fornaguera-Trias, J. | Neuroscience |
| 488 | 2014 | Skillings, Elizabeth A.; Wood, Nigel I.; Morton, A. Jennifer | Brain and behavior |
| 489 | 2014 | Sotnikov, S. V.; Chekmareva, N. Y.; Schmid, B.; Harbich, D.; Malik, V.; Bauer, S.; Kuehne, C.; Markt, P. O.; Deussing, J. M.; Schmidt, M. V.; Landgraf, R. | The European journal of neuroscience |
| 490 | 2014 | Sotnikov, S. V.; Markt, P. O.; Malik, V.; Chekmareva, N. Y.; Naik, R. R.; Sah, A.; Singewald, N.; Holsboer, F.; Czibere, L.; Landgraf, R. | Translational psychiatry |
| 491 | 2014 | Takahashi, Tomohisa; Shimizu, Kunio; Shimazaki, Kuniko; Toda, Hiroyuki; Nibuya, Masashi | Brain research |
| 492 | 2014 | Takuma, Kazuhiro; Maeda, Yuko; Ago, Yukio; Ishihama, Toshihiro; Takemoto, Kosuke; Nakagawa, Akira; Shintani, Norihito; Hashimoto, Hitoshi; Baba, Akemichi; Matsuda, Toshio | Behavioural brain research |
| 493 | 2014 | Teske, Jennifer A.; Perez-Leighton, Claudio E.; Billington, Charles J.; Kotz, Catherine M. | American journal of physiology. Regulatory, integrative and comparative physiology |
| 494 | 2014 | Tipyasang, Rungpiyada; Kunwittaya, Sarun; Mukda, Sujira; Kotchabhakdi, Nittaya J.; Kotchabhakdi, Naiphinich | EXCLI journal |
| 495 | 2014 | Turner, Karly M.; Burne, Thomas H. J. | PloS one |
| 496 | 2014 | Turner, Patricia V.; Sunohara-Neilson, Janet; Ovari, Jelena; Healy, Amanda; Leri, Francesco | Journal of the American Association for Laboratory Animal Science : JAALAS |
| 497 | 2014 | Urakawa, Susumu; Mitsushima, Dai; Shimozuru, Michito; Sakuma, Yasuo; Kondo, Yasuhiko | PloS one |
| 498 | 2014 | Valles, Astrid; Granic, Ivica; De Weerd, Peter; Martens, Gerard J. M. | Learning & memory (Cold Spring Harbor, N.Y.) |
| 499 | 2014 | Venebra-Munoz, Arturo; Corona-Morales, Aleph; Santiago-Garcia, Juan; Melgarejo-Gutierrez, Montserrat; Caba, Mario; Garcia-Garcia, Fabio | Neuroreport |
| 500 | 2014 | Venna, V. R.; Xu, Y.; Doran, S. J.; Patrizz, A.; McCullough, L. D. | Translational psychiatry |
| 501 | 2014 | Venna, Venugopal Reddy; Verma, Rajkumar; O'Keefe, Lena M.; Xu, Yan; Crapser, Joshua; Friedler, Brett; McCullough, Louise D. | Stroke |
| 502 | 2014 | Whiteus, Christina; Freitas, Catarina; Grutzendler, Jaime | Nature |
| 503 | 2014 | Xu, Jia; Sun, Jinling; Xue, Zhaoxia; Li, Xinwang | Neuroreport |
| 504 | 2014 | Yu, Kewei; Wu, Yi; Zhang, Qi; Xie, Hongyu; Liu, Gang; Guo, Zhenzhen; Li, Fang; Jia, Jie; Kuang, Shenyi; Hu, Ruiping | Journal of the neurological sciences |
| 505 | 2014 | Zhang, Xian Geng; Zhang, Hui; Lin, Lin; Yang, Yi Qing; Deng, Ting Ting; Liu, Qin; Liang, Xiao Li; Wang, Mi Qu; Peng, De Zhong | African journal of traditional, complementary, and alternative medicines : AJTCAM |
| 506 | 2014 | Zhang, Yafang; Crofton, Elizabeth J.; Li, Dingge; Lobo, Mary Kay; Fan, Xiuzhen; Nestler, Eric J.; Green, Thomas A. | Frontiers in behavioral neuroscience |
| 507 | 2014 | Zhu, Xiaoqing; Wang, Fang; Hu, Huifang; Sun, Xinde; Kilgard, Michael P.; Merzenich, Michael M.; Zhou, Xiaoming | The Journal of neuroscience : the official journal of the Society for Neuroscience |
| 508 | 2014 | Zigmond, Michael J.; Smeyne, Richard J. | Parkinsonism & related disorders |
| 509 | 2014 | Zlebnik, Natalie E.; Hedges, Valerie L.; Carroll, Marilyn E.; Meisel, Robert L. | Behavioural brain research |
| 510 | 2013 | Ambrogini, P.; Lattanzi, D.; Ciuffoli, S.; Betti, M.; Fanelli, M.; Cuppini, R. | Brain research |
| 511 | 2013 | Avrabos, Charilaos; Sotnikov, Sergey V.; Dine, Julien; Markt, Patrick O.; Holsboer, Florian; Landgraf, Rainer; Eder, Matthias | The Journal of neuroscience : the official journal of the Society for Neuroscience |
| 512 | 2013 | Bengoetxea, Harkaitz; Ortuzar, Naiara; Rico-Barrio, Irantzu; Lafuente, Jose Vicente; Argandona, Enrike G. | Frontiers in cellular neuroscience |
| 513 | 2013 | Bessinis, D. P.; Dalla, C.; Kokras, N.; Pitychoutis, P. M.; Papadopoulou-Daifoti, Z. | Neuroscience |
| 514 | 2013 | Birch, Amy M.; Kelly, Aine M. | Neuropharmacology |
| 515 | 2013 | Castilla-Ortega, Estela; Rosell-Valle, Cristina; Blanco, Eduardo; Pedraza, Carmen; Chun, Jerold; Rodriguez de Fonseca, Fernando; Estivill-Torrus, Guillermo; Santin, Luis J. | Neuroscience research |
| 516 | 2013 | Donato, Flavio; Rompani, Santiago Belluco; Caroni, Pico | Nature |
| 517 | 2013 | Fan, Xiuzhen; Li, Dingge; Zhang, Yafang; Green, Thomas A. | PloS one |
| 518 | 2013 | Franks, Becca; Champagne, Frances A.; Higgins, E. Tory | PloS one |
| 519 | 2013 | Kiss, Peter; Vadasz, Gyongyver; Kiss-Illes, Blanka; Horvath, Gabor; Tamas, Andrea; Reglodi, Dora; Koppan, Miklos | International journal of molecular sciences |
| 520 | 2013 | Nikonenko, Irina; Nikonenko, Alexander; Mendez, Pablo; Michurina, Tatyana V.; Enikolopov, Grigori; Muller, Dominique | Proceedings of the National Academy of Sciences of the United States of America |
| 521 | 2013 | Paez-Martinez, Nayeli; Flores-Serrano, Zoraida; Ortiz-Lopez, Leonardo; Ramirez-Rodriguez, Gerardo | Behavioural brain research |
| 522 | 2013 | Pons-Espinal, Meritxell; Martinez de Lagran, Maria; Dierssen, Mara | Neurobiology of disease |
| 523 | 2013 | Trueba-Saiz, A.; Cavada, C.; Fernandez, A. M.; Leon, T.; Gonzalez, D. A.; Fortea Ormaechea, J.; Lleo, A.; Del Ser, T.; Nunez, A.; Torres-Aleman, I. | Translational psychiatry |
| 524 | 2013 | Yu, Kewei; Wu, Yi; Hu, Yongshan; Zhang, Qi; Xie, Hongyu; Liu, Gang; Chen, Yao; Guo, Zhenzhen; Jia, Jie | Brain research |
| 525 | 2013 | Yu, Kewei; Wu, Yi; Hu, Yongshan; Zhang, Qi; Xie, Hongyu; Liu, Gang; Chen, Yao; Guo, Zhenzhen; Jia, Jie | Neurotoxicology |
| 526 | 2013 | Zhao, Yuan-Yu; Shi, Xiao-Yan; Zhang, Lei; Wu, Hong; Chao, Feng-Lei; Huang, Chun-Xia; Gao, Yuan; Qiu, Xuan; Chen, Lin; Lu, Wei; Tang, Yong | Neuroscience letters |
