## Extended Table 2-2. List of EMBASE References Used for "The Contribution of Environmental Enrichment to Phenotypic Variation in Mice and Rats"

**Extended Data Table 3-2.** List of EMBASE References Used

| **Study** | **Year** | **Author** | **Journal** |
| --- | --- | --- | --- |
| 1 | 2018 | Acosta, J.;Campolongo, M. A.;Hocht, C.;Depino, A. M.;Golombek, D. A.;Agostino, P. V. | European Journal of Neuroscience |
| 2 | 2018 | Benito, E.;Kerimoglu, C.;Ramachandran, B.;Pena-Centeno, T.;Jain, G.;Stilling, R. M.;Islam, M. R.;Capece, V.;Zhou, Q.;Edbauer, D.;Dean, C.;Fischer, A. | Cell Reports |
| 3 | 2018 | Cao, M.;Hu, P. P.;Zhang, Y. L.;Yan, Y. X.;Shields, C. B.;Zhang, Y. P.;Hu, G.;Xiao, M. | CNS Neuroscience and Therapeutics |
| 4 | 2018 | Cattaud, V.;Bezzina, C.;Rey, C. C.;Lejards, C.;Dahan, L.;Verret, L. | Neurobiology of Aging |
| 5 | 2018 | Chen, C. C.;Lu, J.;Yang, R.;Ding, J. B.;Zuo, Y. | Molecular Psychiatry |
| 6 | 2018 | Chun, H.;An, H.;Lim, J.;Woo, J.;Lee, J.;Ryu, H.;Lee, C. J. | Experimental Neurobiology |
| 7 | 2018 | Cinque, C.;Zinni, M.;Zuena, A. R.;Giuli, C.;Alema, S. G.;Catalani, A.;Casolini, P.;Cozzolino, R. | Endocrine Connections |
| 8 | 2018 | Dong, B. E.;Xue, Y.;Sakata, K. | Genes, Brain and Behavior. |
| 9 | 2018 | Duran-Carabali, L. E.;Arcego, D. M.;Odorcyk, F. K.;Reichert, L.;Cordeiro, J. L.;Sanches, E. F.;Freitas, L. D.;Dalmaz, C.;Pagnussat, A.;Netto, C. A. | Molecular Neurobiology |
| 10 | 2018 | Imperio, C. G.;McFalls, A. J.;Hadad, N.;Blanco-Berdugo, L.;Masser, D. R.;Colechio, E. M.;Coffey, A. A.;Bixler, G. V.;Stanford, D. R.;Vrana, K. E.;Grigson, P. S.;Freeman, W. M. | Neuropharmacology |
| 11 | 2018 | Kentner, A. C.;Lima, E.;Migliore, M. M.;Shin, J.;Scalia, S. | Neuroscience |
| 12 | 2018 | Kentrop, J.;Smid, C. R.;Achterberg, E. J. M.;van Ijzendoorn, M. H.;Bakermans-Kranenburg, M. J.;Joels, M.;van der Veen, R. | Frontiers in Behavioral Neuroscience |
| 13 | 2018 | Li, B.;Xu, P.;Wu, S.;Jiang, Z.;Huang, Z.;Li, Q.;Chen, D. | Journal of Alzheimer's Disease |
| 14 | 2018 | Naik, R. R.;Sotnikov, S. V.;Diepold, R. P.;Iurato, S.;Markt, P. O.;Bultmann, A.;Brehm, N.;Mattheus, T.;Lutz, B.;Erhardt, A.;Binder, E. B.;Schmidt, U.;Holsboer, F.;Landgraf, R.;Czibere, L. | Translational Psychiatry |
| 15 | 2018 | Novaes, L. S.;dos Santos, N. B.;Perfetto, J. G.;Goosens, K. A.;Munhoz, C. D. | Psychoneuroendocrinology |
| 16 | 2018 | Otaki, M.;Hirano, T.;Yamaguchi, Y.;Kaida, K.;Koshika, S.;Nagata, K.;Nishimura, M.;Kakinuma, S.;Shimada, Y.;Kobayashi, Y. | International Immunopharmacology |
| 17 | 2018 | Prado Lima, M. G.;Schimidt, H. L.;Garcia, A.;Dare, L. R.;Carpes, F. P.;Izquierdo, I.;Mello-Carpes, P. B. | Proceedings of the National Academy of Sciences of the United States of America |
| 18 | 2018 | Shtoots, L.;Richter-Levin, G.;Hugeri, O.;Anunu, R. | Neurobiology of Learning and Memory |
| 19 | 2018 | Song, S. Y.;Chae, M.;Yu, J. H.;Lee, M. Y.;Pyo, S.;Shin, Y. K.;Baek, A.;Park, J. W.;Park, E. S.;Choi, J. Y.;Cho, S. R. | Frontiers in Neurology |
| 20 | 2018 | Tanichi, M.;Toda, H.;Shimizu, K.;Koga, M.;Saito, T.;Enomoto, S.;Boku, S.;Asai, F.;Mitsui, Y.;Nagamine, M.;Fujita, M.;Yoshino, A. | Biochemical and Biophysical Research Communications |
| 21 | 2018 | Wahl, A. S.;Erlebach, E.;Brattoli, B.;Buchler, U.;Kaiser, J.;Ineichen, B. V.;Mosberger, A. C.;Schneeberger, S.;Imobersteg, S.;Wieckhorst, M.;Stirn, M.;Schroeter, A.;Ommer, B.;Schwab, M. E. | Journal of Cerebral Blood Flow and Metabolism. |
| 22 | 2018 | Xie, H.;Yu, K.;Zhou, N.;Shen, X.;Tian, S.;Zhang, B.;Wang, Y.;Wu, J.;Liu, G.;Jiang, C.;Hu, R.;Ayata, C.;Wu, Y. | Translational Stroke Research |
| 23 | 2018 | Xu, H.;Rajsombath, M. M.;Weikop, P.;Selkoe, D. J. | EMBO Molecular Medicine |
| 24 | 2018 | Zhang, Y.;Wang, G.;Wang, L.;Zhao, J.;Huang, R.;Xiong, Q. | Human Vaccines and Immunotherapeutics |
| 25 | 2017 | Bice, B. D.;Stephens, M. R.;Georges, S. J.;Venancio, A. R.;Bermant, P. C.;Warncke, A. V.;Affolter, K. E.;Hidalgo, J. R.;Angus-Hill, M. L. | Cell Reports |
| 26 | 2017 | Blanco, M. B.;De Souza, R. R.;Canto-De-souza, A. | Psychology and Neuroscience |
| 27 | 2017 | Cutuli, D.;Berretta, E.;Pasqualini, G.;De Bartolo, P.;Caporali, P.;Laricchiuta, D.;Sampedro-Piquero, P.;Gelfo, F.;Pesoli, M.;Foti, F.;Begega, A.;Petrosini, L. | Frontiers in Behavioral Neuroscience |
| 28 | 2017 | Fisch, J.;Oliveira, I. V.;Fank, J.;Paim, L. M.;Zandona, M. R.;Lopes, E. F.;Mello, F. B.;Oliveira, A. T. | Theriogenology |
| 29 | 2017 | Garofalo, S.;Porzia, A.;Mainiero, F.;Di Angelantonio, S.;Cortese, B.;Basilico, B.;Pagani, F.;Cignitti, G.;Chece, G.;Maggio, R.;Tremblay, M. E.;Savage, J.;Bisht, K.;Esposito, V.;Bernardini, G.;Seyfried, T.;Mieczkowski, J.;Stepniak, K.;Kaminska, B.;Santoni, A.;Limatola, C. | eLife |
| 30 | 2017 | Gjendal, K.;Sorensen, D. B.;Kiersgaard, M. K.;Ottesen, J. L. | Animal Welfare |
| 31 | 2017 | Guan, S. Z.;Ji, W. J.;Jiang, Y.;Ning, L.;Lian, Y. L.;Liu, J. W. | International Journal of Clinical and Experimental Medicine |
| 32 | 2017 | Lynch, W. J.;Tan, L.;Narmeen, S.;Beiter, R.;Brunzell, D. H. | Physiology and Behavior. |
| 33 | 2017 | Nawaz, A.;Batool, Z.;Ahmed, S.;Khaliq, S.;Sajid, I.;Anis, L.;Haider, S. | Pakistan Veterinary Journal |
| 34 | 2017 | Peterson, E. K.;Hughes, R. N. | Current Psychopharmacology |
| 35 | 2017 | Rajab, E.;Al-Kafaji, G.;Al Enazi, H.;Al Qassab, N.;Sakusic, A.;Kamal, A. | Bahrain Medical Bulletin |
| 36 | 2017 | Smith, B. L.;Morano, R. L.;Ulrich-Lai, Y. M.;Myers, B.;Solomon, M. B.;Herman, J. P. | Stress |
| 37 | 2017 | Song, Y.;Gan, Y.;Wang, Q.;Meng, Z.;Li, G.;Shen, Y.;Wu, Y.;Li, P.;Yao, M.;Gu, J.;Tu, H. | Cancer Research |
| 38 | 2017 | Stairs, D. J.;Ewin, S. E.;Kangiser, M. M.;Pfaff, M. N. | Experimental and Clinical Psychopharmacology |
| 39 | 2017 | Taffou, M.;Ondrej, J.;O'Sullivan, C.;Warusfel, O.;Dubal, S.;Viaud-Delmon, I. | Psychological research |
| 40 | 2017 | Torres, R. F.;Hidalgo, C.;Kerr, B. | Frontiers in Molecular Neuroscience |
| 41 | 2017 | Zeleznikow-Johnston, A.;Burrows, E. L.;Renoir, T.;Hannan, A. J. | Neuropharmacology |
| 42 | 2017 | Zhang, S.;Xie, H.;Wang, Y.;Li, D.;Du, L.;Wu, Y.;Yang, G. Y. | European Journal of Inflammation |
| 43 | 2016 | Dostes, S.;Dubreucq, S.;Ladeveze, E.;Marsicano, G.;Abrous, D. N.;Chaouloff, F.;Koehl, M. | Hippocampus |
| 44 | 2016 | Fernandez-Montoya, J.;Buendia, I.;Martin, Y. B.;Egea, J.;Negredo, P.;Avendano, C. | Frontiers in Molecular Neuroscience |
| 45 | 2016 | Fuchs, F.;Herbeaux, K.;Aufrere, N.;Kelche, C.;Mathis, C.;Barbelivien, A.;Majchrzak, M. | Learning and Memory |
| 46 | 2016 | Galaj, E.;Manuszak, M.;Ranaldi, R. | Drug and Alcohol Dependence |
| 47 | 2016 | Greifzu, F.;Kalogeraki, E.;Lowel, S. | Neurobiology of Aging |
| 48 | 2016 | Huttenrauch, M.;Brauss, A.;Kurdakova, A.;Borgers, H.;Klinker, F.;Liebetanz, D.;Salinas-Riester, G.;Wiltfang, J.;Klafki, H. W.;Wirths, O. | Translational Psychiatry |
| 49 | 2016 | Mehta-Raghavan, N. S.;Wert, S. L.;Morley, C.;Graf, E. N.;Redei, E. E. | Translational psychiatry |
| 50 | 2016 | Nakamura, Y.;Ueno, A.;Nunomura, Y.;Nakagaki, K.;Takeda, S.;Suzuki, K. | Experimental Animals |
| 51 | 2016 | Pitzer, C.;Kuner, R.;Tappe-Theodor, A. | Molecular Pain |
| 52 | 2016 | Sansevero, G.;Begenisic, T.;Mainardi, M.;Sale, A. | Experimental Neurology |
| 53 | 2015 | Bezzina, C.;Verret, L.;Halley, H.;Dahan, L.;Rampon, C. | Frontiers in Aging Neuroscience |
| 54 | 2015 | Cruz, J. N.;Lima, D. D.;Dal Magro, D. D.;Cruz, J. G. P. | Scandinavian Journal of Laboratory Animal Science |
| 55 | 2015 | During, M. J.;Liu, X.;Huang, W.;Magee, D.;Slater, A.;McMurphy, T.;Wang, C.;Cao, L. | Endocrinology |
| 56 | 2015 | Garofalo, S.;D'Alessandro, G.;Chece, G.;Brau, F.;Maggi, L.;Rosa, A.;Porzia, A.;Mainiero, F.;Esposito, V.;Lauro, C.;Benigni, G.;Bernardini, G.;Santoni, A.;Limatola, C. | Nature Communications |
| 57 | 2015 | Mesa-Gresa, P.;Ramos-Campos, M.;Redolat, R. | Current Topics in Pharmacology |
| 58 | 2015 | Yang, Y.;Zhang, J.;Xiong, L.;Deng, M.;Wang, J.;Xin, J.;Liu, H. | Journal of Molecular Neuroscience |
| 59 | 2015 | Yeung, S. T.;Martinez-Coria, H.;Ager, R. R.;Rodriguez-Ortiz, C. J.;Baglietto-Vargas, D.;LaFerla, F. M. | Brain Research Bulletin |
| 60 | 2014 | Coke-Murphy, C.;Buendia, M. A.;Saborido, T. P.;Stanwood, G. D. | Translational Neuroscience |
| 61 | 2014 | de Oliveira Soares, R.;de Oliveira, L. M.;Almeida, S. S. | Psychology and Neuroscience |
| 62 | 2014 | Dorfman, D.;Aranda, M. L.;Gonzalez Fleitas, M. F.;Chianelli, M. S.;Fernandez, D. C.;Sande, P. H.;Rosenstein, R. E. | PLoS ONE |
| 63 | 2014 | Hofford, R. S.;Darna, M.;Wilmouth, C. E.;Dwoskin, L. P.;Bardo, M. T. | Behavioural Brain Research |
| 64 | 2014 | Kulesskaya, N.;Karpova, N. N.;Ma, L.;Tian, L.;Voikar, V. | Frontiers in Behavioral Neuroscience |
| 65 | 2014 | Marques, M. R.;Stigger, F.;Segabinazi, E.;Augustin, O. A.;Barbosa, S.;Piazza, F. V.;Achaval, M.;Marcuzzo, S. | Behavioural Brain Research |
| 66 | 2014 | Vachon, P. | Scandinavian Journal of Laboratory Animal Science |
| 67 | 2013 | Adams, E.;Klug, J.;Quast, M.;Stairs, D. J. | Drug and Alcohol Dependence |
| 68 | 2013 | Alwis, D. S.;Rajan, R. | Frontiers in Cellular Neuroscience. |
| 69 | 2013 | Ambrogini, P.;Lattanzi, D.;Ciuffoli, S.;Betti, M.;Fanelli, M.;Cuppini, R. | Brain Research |
| 70 | 2013 | Aumann, T. D.;Tomas, D.;Horne, M. K. | Brain and Behavior |
| 71 | 2013 | Avrabos, C.;Sotnikov, S. V.;Dine, J.;Markt, P. O.;Holsboer, F.;Landgraf, R.;Eder, M. | Journal of Neuroscience |
| 72 | 2013 | Baldini, S.;Restani, L.;Baroncelli, L.;Coltelli, M.;Franco, R.;Cenni, M. C.;Maffei, L.;Berardi, N. | Journal of Neuroscience |
| 73 | 2013 | Barak, B.;Shvarts-Serebro, I.;Modai, S.;Gilam, A.;Okun, E.;Michaelson, D. M.;Mattson, M. P.;Shomron, N.;Ashery, U. | Translational psychiatry |
| 74 | 2013 | Baraldi, T.;Schowe, N. M.;Balthazar, J.;Monteiro-Silva, K. C.;Albuquerque, M. S.;Buck, H. S.;Viel, T. A. | Experimental Gerontology |
| 75 | 2013 | Bechara, R. G.;Kelly, T. | Behavioural Brain Research |
| 76 | 2013 | Bengoetxea, H.;Ortuzar, N.;Rico-Barrio, I.;Lafuente, J. V.;Argandona, E. G. | Frontiers in Cellular Neuroscience |
| 77 | 2013 | Birch, A. M.;McGarry, N. B.;Kelly, A. M. | Hippocampus |
| 78 | 2013 | Bonaccorsi, J.;Cintoli, S.;Mastrogiacomo, R.;Baldanzi, S.;Braschi, C.;Pizzorusso, T.;Cenni, M. C.;Berardi, N. | Neural Plasticity |
| 79 | 2013 | Branchi, I.;Santarelli, S.;D'Andrea, I.;Alleva, E. | Hormones and Behavior |
| 80 | 2013 | Cymerblit-Sabba, A.;Lasri, T.;Gruper, M.;Aga-Mizrachi, S.;Zubedat, S.;Avital, A. | Behavioural Brain Research |
| 81 | 2013 | De Jong, T. R.;Harris, B. N.;Perea-Rodriguez, J. P.;Saltzman, W. | Psychoneuroendocrinology |
| 82 | 2013 | Diniz, D. G.;Foro, C. A. R.;Sosthenes, M. C. K.;Demachki, S.;Gomes, G. F.;Malerba, G. A.;Naves, T. B.;Cavalcante, E. A. D.;Sousa, A. M. C.;Ferreira, F. A. B.;Anjos, P. C. S.;Neto, A. L. C.;Pinho, B. G.;Brito, M. V.;Freitas, P. S. L.;Casseb, S. M. M.;Silva, E. V. P.;Nunes, M. R. T.;Diniz, J. A. P.;Cunningham, C.;Perry, V. H.;Vasconcelos, P. F. C.;Diniz, C. W. P. | European Journal of Inflammation |
| 83 | 2013 | Dorfman, D.;Fernandez, D. C.;Chianelli, M.;Miranda, M.;Aranda, M. L.;Rosenstein, R. E. | Experimental Neurology |
| 84 | 2013 | Fan, X.;Li, D.;Lichti, C. F.;Green, T. A. | PLoS ONE |
| 85 | 2013 | Fares, R. P.;Belmeguenai, A.;Sanchez, P. E.;Kouchi, H. Y.;Bodennec, J.;Morales, A.;Georges, B.;Bonnet, C.;Bouvard, S.;Sloviter, R. S.;Bezin, L. | PLoS ONE |
| 86 | 2013 | Freund, J.;Brandmaier, A. M.;Lewejohann, L.;Kirste, I.;Kritzler, M.;Kruger, A.;Sachser, N.;Lindenberger, U.;Kempermann, G. | Science |
| 87 | 2013 | Fuss, J.;Richter, S. H.;Steinle, J.;Deubert, G.;Hellweg, R.;Gass, P. | Behavioural Brain Research |
| 88 | 2013 | Garrido, P.;De Blas, M.;Ronzoni, G.;Cordero, I.;Anton, M.;Gine, E.;Santos, A.;Del Arco, A.;Segovia, G.;Mora, F. | Journal of Neural Transmission |
| 89 | 2013 | Grimm, J. W.;Weber, R.;Barnes, J.;Koerber, J.;Dorsey, K.;Glueck, E. | PLoS ONE |
| 90 | 2013 | Harati, H.;Barbelivien, A.;Herbeaux, K.;Muller, M. A.;Engeln, M.;Kelche, C.;Cassel, J. C.;Majchrzak, M. | Age |
| 91 | 2013 | Harrison, D. J.;Busse, M.;Openshaw, R.;Rosser, A. E.;Dunnett, S. B.;Brooks, S. P. | Experimental Neurology |
| 92 | 2013 | Horvath, G.;Reglodi, D.;Vadasz, G.;Farkas, J.;Kiss, P. | International Journal of Molecular Sciences |
| 93 | 2013 | Hu, Y. S.;Long, N.;Pigino, G.;Brady, S. T.;Lazarov, O. | PLoS ONE |
| 94 | 2013 | Hughes, R. N. | Journal of Caffeine Research |
| 95 | 2013 | Hughes, R. N.;Otto, M. T. | Progress in Neuro-Psychopharmacology and Biological Psychiatry |
| 96 | 2013 | Jain, V.;Baitharu, I.;Prasad, D.;Ilavazhagan, G. | PLoS ONE |
| 97 | 2013 | Jenks, K. R.;Lucas, M. M.;Duffy, B. A.;Robbins, A. A.;Gimi, B.;Barry, J. M.;Scott, R. C. | PLoS ONE |
| 98 | 2013 | Johnson, E. M.;Traver, K. L.;Hoffman, S. W.;Harrison, C. R.;Herman, J. P. | Frontiers in Behavioral Neuroscience. |
| 99 | 2013 | Kajimoto, K.;Allan, A.;Cunningham, L. A. | PloS one |
| 100 | 2013 | Kershaw, M. H.;Westwood, J. A.;Darcy, P. K. | F1000Research |
| 101 | 2013 | Kirkpatrick, K.;Marshall, A. T.;Clarke, J.;Cain, M. E. | Behavioral Neuroscience |
| 102 | 2013 | Kiss, P.;Szabadfi, K.;Horvath, G.;Tamas, A.;Farkas, J.;Gabriel, R.;Reglodi, D. | International Journal of Molecular Sciences |
| 103 | 2013 | Lee, M. Y.;Yu, J. H.;Kim, J. Y.;Seo, J. H.;Park, E. S.;Kim, C. H.;Kim, H.;Cho, S. R. | Neurorehabilitation and Neural Repair |
| 104 | 2013 | Lehmann, M. L.;Brachman, R. A.;Martinowich, K.;Schloesser, R. J.;Herkenham, M. | Journal of Neuroscience |
| 105 | 2013 | Li, S.;Jin, M.;Zhang, D.;Yang, T.;Koeglsperger, T.;Fu, H.;Selkoe, D. J. | Neuron |
| 106 | 2013 | Lobo, M. K.;Zaman, S.;Damez-Werno, D. M.;Koo, J. W.;Bagot, R. C.;DiNieri, J. A.;Nugent, A.;Finkel, E.;Chaudhury, D.;Chandra, R.;Riberio, E.;Rabkin, J.;Mouzon, E.;Cachope, R.;Cheer, J. F.;Han, M. H.;Dietz, D. M.;Self, D. W.;Hurd, Y. L.;Vialou, V.;Nestler, E. J. | Journal of Neuroscience |
| 107 | 2013 | Macri, S.;Ceci, C.;Altabella, L.;Canese, R.;Laviola, G. | Scientific reports |
| 108 | 2013 | Maesako, M.;Uemura, K.;Iwata, A.;Kubota, M.;Watanabe, K.;Uemura, M.;Noda, Y.;Asada-Utsugi, M.;Kihara, T.;Takahashi, R.;Shimohama, S.;Kinoshita, A. | PLoS ONE |
| 109 | 2013 | McQuaid, R. J.;Audet, M. C.;Jacobson-Pick, S.;Anisman, H. | International Journal of Neuropsychopharmacology |
| 110 | 2013 | Mendes, F. D. C. C. D. S.;de Almeida, M. N. F.;Felicio, A. P. G.;Fadel, A. C.;Silva, D. D. J.;Borralho, T. G.;da Silva, R. P.;Bento-Torres, J.;Vasconcelos, P. F. D. C.;Perry, V. H.;Ramos, E. M. L. S.;Picanco-Diniz, C. W.;Sosthenes, M. C. K. | BMC Neuroscience |
| 111 | 2013 | Mesa-Gresa, P.;Perez-Martinez, A.;Redolat, R. | Physiology and Behavior |
| 112 | 2013 | Mesa-Gresa, P.;Perez-Martinez, A.;Redolat, R. | Aggressive Behavior |
| 113 | 2013 | Monaco, C. M.;Mattiola, V. V.;Folweiler, K. A.;Tay, J. K.;Yelleswarapu, N. K.;Curatolo, L. M.;Matter, A. M.;Cheng, J. P.;Kline, A. E. | Experimental Neurology |
| 114 | 2013 | Montarolo, F.;Parolisi, R.;Hoxha, E.;Boda, E.;Tempia, F. | PLoS ONE |
| 115 | 2013 | Okva, K.;Nevalainen, T.;Pokk, P. | Laboratory Animals |
| 116 | 2013 | Ortuzar, N.;Rico-Barrio, I.;Bengoetxea, H.;Argandona, E. G.;Lafuente, J. V. | Behavioural Brain Research |
| 117 | 2013 | Pang, T. Y.;Du, X.;Catchlove, W. A.;Renoir, T.;Lawrence, A. J.;Hannan, A. J. | Frontiers in Pharmacology |
| 118 | 2013 | Pascual, R.;Bustamante, C. | Acta Neurobiologiae Experimentalis |
| 119 | 2013 | Peruzzaro, S. T.;Gallagher, J.;Dunkerson, J.;Fluharty, S.;Mudd, D.;Hoane, M. R.;Smith, J. S. | Restorative Neurology and Neuroscience |
| 120 | 2013 | Pritchard, L. M.;Van Kempen, T. A.;Zimmerberg, B. | Neuroscience Letters |
| 121 | 2013 | Ragu Varman, D.;Marimuthu, G.;Emmanuvel Rajan, K. | Journal of Neuroscience Research |
| 122 | 2013 | Ransome, M. I.;Hannan, A. J. | Molecular and Cellular Neuroscience |
| 123 | 2013 | Ravenelle, R.;Byrnes, E. M.;Byrnes, J. J.;McInnis, C.;Park, J. H.;Donaldson, S. T. | Behavioural Brain Research |
| 124 | 2013 | Raz, S. | Physiology and Behavior |
| 125 | 2013 | Reichmann, F.;Painsipp, E.;Holzer, P. | PLoS ONE |
| 126 | 2013 | Renoir, T.;Pang, T. Y.;Mo, C.;Chan, G.;Chevarin, C.;Lanfumey, L.;Hannan, A. J. | Journal of Physiology |
| 127 | 2013 | Reynolds, S.;Urruela, M.;Devine, D. P. | Autism Research |
| 128 | 2013 | Richter, S. H.;Zeuch, B.;Riva, M. A.;Gass, P.;Vollmayr, B. | Behavioural Brain Research |
| 129 | 2013 | Rojas, J. J.;Deniz, B. F.;Miguel, P. M.;Diaz, R.;Hermel, E. D. E. S.;Achaval, M.;Netto, C. A.;Pereira, L. O. | Experimental Neurology |
| 130 | 2013 | Ruscher, K.;Kuric, E.;Liu, Y.;Walter, H. L.;Issazadeh-Navikas, S.;Englund, E.;Wieloch, T. | Journal of Cerebral Blood Flow and Metabolism |
| 131 | 2013 | Sampedro-Piquero, P.;Begega, A.;Zancada-Menendez, C.;Cuesta, M.;Arias, J. L. | Neuroscience |
| 132 | 2013 | Sampedro-Piquero, P.;Zancada-Menendez, C.;Begega, A.;Rubio, S.;Arias, J. L. | Brain Research Bulletin |
| 133 | 2013 | Sato, Y.;Bernier, F.;Suzuki, I.;Kotani, S.;Nakagawa, M.;Oda, Y. | Journal of Lipid Research |
| 134 | 2013 | Schreiber, W. B.;St. Cyr, S. A.;Jablonski, S. A.;Hunt, P. S.;Klintsova, A. Y.;Stanton, M. E. | Developmental Psychobiology |
| 135 | 2013 | Seo, J. H.;Kim, H.;Park, E. S.;Lee, J. E.;Kim, D. W.;Kim, H. O.;Im, S. H.;Yu, J. H.;Kim, J. Y.;Lee, M. Y.;Kim, C. H.;Cho, S. R. | Cell Transplantation |
| 136 | 2013 | Seo, J. H.;Yu, J. H.;Suh, H.;Kim, M. S.;Cho, S. R. | PLoS ONE |
| 137 | 2013 | Shemesh, Y.;Sztainberg, Y.;Forkosh, O.;Shlapobersky, T.;Chen, A.;Schneidman, E. | eLife |
| 138 | 2013 | Shen, X.;Dong, Y.;Xu, Z.;Wang, H.;Miao, C.;Soriano, S. G.;Sun, D.;Baxter, M. G.;Zhang, Y.;Xie, Z. | Anesthesiology |
| 139 | 2013 | Shinohara, Y.;Hosoya, A.;Hirase, H. | Nature Communications |
| 140 | 2013 | Soares, R. O.;Oliveira, L. M.;Marchini, J. S.;Rodrigues, J. A.;Elias, L. L. K.;Almeida, S. S. | Nutritional Neuroscience |
| 141 | 2013 | Speisman, R. B.;Kumar, A.;Rani, A.;Pastoriza, J. M.;Severance, J. E.;Foster, T. C.;Ormerod, B. K. | Neurobiology of Aging |
| 142 | 2013 | Tanti, A.;Westphal, W. P.;Girault, V.;Brizard, B.;Devers, S.;Leguisquet, A. M.;Surget, A.;Belzung, C. | Hippocampus |
| 143 | 2013 | Tyler, C. R.;Allan, A. M. | PLoS ONE |
| 144 | 2013 | Urakawa, S.;Takamoto, K.;Hori, E.;Sakai, N.;Ono, T.;Nishijo, H. | BMC neuroscience |
| 145 | 2013 | Vachon, P.;Millecamps, M.;Low, L.;Thompsosn, S. J.;Pailleux, F.;Beaudry, F.;Bushnell, C. M.;Stone, L. S. | Behavioral and Brain Functions |
| 146 | 2013 | Vazquez-Sanroman, D.;Sanchis-Segura, C.;Toledo, R.;Hernandez, M. E.;Manzo, J.;Miquel, M. | Behavioural Brain Research |
| 147 | 2013 | Veeraraghavalu, K.;Sisodia, S. S. | Proceedings of the National Academy of Sciences of the United States of America |
| 148 | 2013 | Verret, L.;Krezymon, A.;Halley, H.;Trouche, S.;Zerwas, M.;Lazouret, M.;Lassalle, J. M.;Rampon, C. | Neurobiology of Aging |
| 149 | 2013 | Vivinetto, A. L.;Suarez, M. M.;Rivarola, M. A. | Behavioural Brain Research |
| 150 | 2013 | Wang, B. S.;Feng, L.;Liu, M.;Liu, X.;Cang, J. | Neuron |
| 151 | 2013 | Xie, H.;Wu, Y.;Jia, J.;Liu, G.;Zhang, F.;Zhang, Q.;Yu, K.;Hu, Y.;Bai, Y.;Hu, R. | Brain Research |
| 152 | 2013 | Yang, S.;Li, C.;Qiu, X.;Zhang, L.;Lu, W.;Chen, L.;Zhao, Y. Y.;Shi, X. Y.;Huang, C. X.;Cheng, G. H.;Tang, Y. | Neuroscience |
| 153 | 2013 | Zeeb, F. D.;Wong, A. C.;Winstanley, C. A. | Psychopharmacology |
| 154 | 2013 | Zhang, L.;Zhang, J.;Sun, H.;Zhu, H.;Liu, H.;Yang, Y. | Pharmacology Biochemistry and Behavior |
| 155 | 2013 | Zhu, J.;Bardo, M. T.;Dwoskin, L. P. | Synapse |
