## Extended Data File 4-1. Pairwise Comparisons for "The Contribution of Environmental Enrichment to Phenotypic Variation in Mice and Rats"

**Extended Data Table 4-1**. Pairwise comparisons for naïve controls and naïve enriched rats and mice in which all behavior, physiology, and anatomy traits are combined.

| Description | Trait Category | Mean | Standard  Deviation | Standard Error | 95% confidence interval | | t | df | p-value  (two tailed) |
| --- | --- | --- | --- | --- | --- | --- | --- | --- | --- |
|  |  |  |  |  | Lower | Upper |  |  |  |
| Main effect of housing | all traits combined | .342 | 22.985 | .924 | -1.472 | 2.156 | .370 | 618 | .711 |
