## Extended Data File 4-3. Pairwise Comparisons for "The Contribution of Environmental Enrichment to Phenotypic Variation in Mice and Rats"

**Extended Data Table 4-3**. Pairwise comparisons for naïve controls and naïve enriched rats and mice by each individual trait.

| Description | Trait Category | Mean | Standard  Deviation | Standard Error | 95% confidence interval | | t | df | p-value  (two tailed) |
| --- | --- | --- | --- | --- | --- | --- | --- | --- | --- |
|  |  |  |  |  | Lower | Upper |  |  |  |
| Main effect of housing | Behavior (all) | .357 | 29.538 | 1.538 | -2.667 | 3.381 | .232 | 368 | .817 |
| Main effect of housing | Physiology  (all) | .320 | 4.661 | .295 | -.260 | .901 | 1.086 | 249 | .278 |
| Main effect of housing | Anatomy | .008 | .436 | .055 | -.103 | .119 | .148 | 61 | .883 |
| Main effect of housing | Behavior (CNS) | .445 | 32.34 | 1.84 | -3.18 | 4.070 | .241 | 307 | .809 |
| Main effect of housing | Behavior (other) | -.087 | .434 | .056 | -.198 | .025 | -1.560 | 60 | .124 |
| Main effect of housing | Immune System | 3.213 | 14.513 | 2.90 | -2.78 | 9.203 | 1.107 | 24 | .279 |
| Main effect of housing | Molecules | -.016 | .890 | .079 | -.173 | .141 | -.202 | 125 | .840 |
| Main effect of housing | Organ Function | -.021 | .750 | .177 | -.394 | .352 | -.119 | 17 | .907 |
| Main effect of housing | E-phys | .085 | .276 | .063 | -.048 | .218 | 1.341 | 18 | .197 |
