## Extended Data File 4-4. Pairwise Comparisons for "The Contribution of Environmental Enrichment to Phenotypic Variation in Mice and Rats"

**Extended Data Table 4-4**. Pairwise comparisons for treated/manipulated controls and treated/manipulated enriched rats and mice by each individual trait.

| Description | Trait Category | Mean | Standard  Deviation | Standard Error | 95% confidence interval | | t | df | p-value  (two tailed) |
| --- | --- | --- | --- | --- | --- | --- | --- | --- | --- |
|  |  |  |  |  | Lower | Upper |  |  |  |
| Main effect of housing | Behavior (all) | .110 | .886 | .02 | .007 | .212 | 2.120 | 290 | .035*  Control (0.67 ± 0.06) more variable than EE (0.56 ± 0.04) |
| Main effect of housing | Physiology  (all) | -.060 | .573 | .038 | -.135 | .016 | -1.559 | 222 | .120 |
| Main effect of housing | Anatomy | .0134 | .493 | .0641 | -.115 | .142 | .217 | 58 | .829 |
| Main effect of housing | Behavior (CNS) | .121 | .965 | .062 | -.001 | .243 | 1.959 | 243 | .051 |
| Main effect of housing | Behavior (other) | .054 | .167 | .024 | .005 | .103 | 2.211 | 46 | .032*  Control (0.73 ±  0.07) more variable than EE (0.60 ± 0.05) |
| Main effect of housing | Immune System | -.098 | .363 | .081 | -.2678 | .072 | -1.213 | 19 | .240 |
| Main effect of housing | Molecules | -.063 | .614 | .059 | -.179 | .054 | -1.066 | 108 | .289 |
| Main effect of housing | Organ Function | -.139 | .779 | .174 | -.503 | .225 | -.798 | 19 | .435 |
| Main effect of housing | E-phys | -.172 | .503 | .130 | -.451 | .106 | -1.329 | 14 | .205 |

*Mean ±SEM
