## Extended Data File 4-6. Pairwise Comparisons for "The Contribution of Environmental Enrichment to Phenotypic Variation in Mice and Rats"

**Extended Data Table 4-6**. Pairwise comparisons for treated/manipulated control and treated/manipulated enriched rats in which all behavior, physiology, and anatomy traits are combined.

| Description | Trait Category | Mean | Standard  Deviation | Standard Error | 95% confidence interval | | t | df | p-value  (two tailed) |
| --- | --- | --- | --- | --- | --- | --- | --- | --- | --- |
|  |  |  |  |  | Lower | Upper |  |  |  |
| Main effect of housing | all traits combined | .0496 | .753 | .0452 | -.0394 | .1387 | 1.097 | 276 | .274 |
