## Extended Data File 4-7. Pairwise Comparisons for "The Contribution of Environmental Enrichment to Phenotypic Variation in Mice and Rats"

**Extended Data Table 4-7**. Pairwise comparisons for naïve controls and naïve enriched rats by each individual trait.

| Description | Trait Category | Mean | Standard  Deviation | Standard Error | 95% confidence interval | | t | df | p-value  (two tailed) |
| --- | --- | --- | --- | --- | --- | --- | --- | --- | --- |
|  |  |  |  |  | Lower | Upper |  |  |  |
| Main effect of housing | Behavior (all) | .600 | 37.930 | 2.534 | -4.394 | 5.594 | .237 | 223 | .813 |
| Main effect of housing | Physiology  (all) | -.021 | .911 | .080 | -.179 | .136 | -.267 | 129 | .790 |
| Main effect of housing | Anatomy | -.037 | .336 | .056 | -.150 | .077 | -.658 | 35 | .515 |
| Main effect of housing | Behavior (CNS) | .760 | 41.868 | 3.08 | -5.33 | 6.849 | .246 | 183 | .806 |
| Main effect of housing | Behavior (other) | -.134 | .512 | .081 | -.298 | .030 | -1.659 | 39 | .105 |
| Main effect of housing | Immune System | N/A | N/A | N/A | N/A | N/A | N/A | N/A | N/A |
| Main effect of housing | Molecules | -.045 | 1.238 | .159 | -.362 | .272 | -.286 | 60 | .776 |
| Main effect of housing | Organ Function | -.018 | .773 | .187 | -.415 | .379 | -.096 | 16 | .925 |
| Main effect of housing | E-phys | .101 | .299 | .075 | -.058 | .261 | 1.357 | 15 | .195 |
