## Extended Data File 4-8. Pairwise Comparisons for "The Contribution of Environmental Enrichment to Phenotypic Variation in Mice and Rats"

**Extended Data Table 4-8**. Pairwise comparisons for treated/manipulated controls and treated/manipulated enriched rats by each individual trait.

| Description | Trait Category | Mean | Standard  Deviation | Standard Error | 95% confidence interval | | t | df | p-value  (two tailed) |
| --- | --- | --- | --- | --- | --- | --- | --- | --- | --- |
|  |  |  |  |  | Lower | Upper |  |  |  |
| Main effect of housing | Behavior (all) | .105 | .848 | .065 | -.023 | .233 | 1.621 | 170 | .107 |
| Main effect of housing | Physiology  (all) | -.040 | .560 | .054 | -.148 | .0679 | -.734 | 105 | .464 |
| Main effect of housing | Anatomy | .0392 | .603 | .099 | -.162 | .240 | .395 | 36 | .695 |
| Main effect of housing | Behavior (CNS) | .117 | .931 | .079 | -.038 | .271 | 1.487 | 140 | .139 |
| Main effect of housing | Behavior (other) | .052 | .181 | .033 | -.016 | .119 | 1.558 | 29 | .130 |
| Main effect of housing | Immune System | N/A | N/A | N/A | N/A | N/A | N/A | N/A | N/A |
| Main effect of housing | Molecules | -.065 | .446 | .072 | -.212 | .081 | -.903 | 37 | .372 |
| Main effect of housing | Organ Function | -.145 | .799 | .183 | -.531 | .240 | -.793 | 18 | .438 |
| Main effect of housing | E-phys | -.036 | .208 | .060 | -.168 | .096 | -.608 | 11 | .556 |
