## Extended Data File 4-9. Pairwise Comparisons for "The Contribution of Environmental Enrichment to Phenotypic Variation in Mice and Rats"

**Extended Data Table 4-9**. Pairwise comparisons for naïve controls and naïve enriched mice in which all behavior, physiology, and anatomy traits are combined.

| Description | Trait Category | Mean | Standard  Deviation | Standard Error | 95% confidence interval | | t | df | p-value  (two tailed) |
| --- | --- | --- | --- | --- | --- | --- | --- | --- | --- |
|  |  |  |  |  | Lower | Upper |  |  |  |
| Main effect of housing | all traits combined | .3022 | 4.572 | .280 | -.250 | .855 | 1.076 | 264 | .283 |
