## Extended Data File 4-10. Pairwise Comparisons for "The Contribution of Environmental Enrichment to Phenotypic Variation in Mice and Rats"

**Extended Data Table 4-10**. Pairwise comparisons for treated/manipulated controls and treated/manipulated enriched mice in which all behavior, physiology, and anatomy traits are combined.

| Description | Trait Category | Mean | Standard  Deviation | Standard Error | 95% confidence interval | | t | df | p-value  (two tailed) |
| --- | --- | --- | --- | --- | --- | --- | --- | --- | --- |
|  |  |  |  |  | Lower | Upper |  |  |  |
| Main effect of housing | all traits combined | .021 | .792 | .051 | -.0804 | .122 | .407 | 236 | .684 |
