## Extended Data File 4-11. Pairwise Comparisons for "The Contribution of Environmental Enrichment to Phenotypic Variation in Mice and Rats"

**Extended Data Table 4-11**. Pairwise comparisons for naïve controls and naïve enriched mice by each individual trait.

| Description | Trait Category | Mean | Standard  Deviation | Standard Error | 95% confidence interval | | t | df | p-value  (two tailed) |
| --- | --- | --- | --- | --- | --- | --- | --- | --- | --- |
|  |  |  |  |  | Lower | Upper |  |  |  |
| Main effect of housing | Behavior (all) | -.0187 | 1.228 | .102 | -.220 | .183 | -.183 | 144 | .855 |
| Main effect of housing | Physiology  (all) | .690 | 6.655 | .608 | -.513 | 1.893 | 1.136 | 119 | .258 |
| Main effect of housing | Anatomy | .0701 | .547 | .107 | -.1501 | .293 | .657 | 25 | .517 |
| Main effect of housing | Behavior (CNS) | -.0223 | 1.327 | .119 | -.258 | .213 | -.190 | 123 | .850 |
| Main effect of housing | Behavior (other) | .005 | .197 | .043 | -.085 | .094 | .106 | 20 | .917 |
| Main effect of housing | Immune System | 3.212 | 14.513 | 2.902 | -2.778 | 9.203 | 1.107 | 24 | .279 |
| Main effect of housing | Molecules | .012 | .332 | .041 | -.071 | .094 | .278 | 64 | .782 |
| Main effect of housing | Organ Function | -.004 | .006 | .004 | -.0187 | .011 | -1.051 | 2 | .403 |
| Main effect of housing | E-phys | .0705 | .547 | .107 | -.151 | .292 | .657 | 25 | .517 |
