## Extended Data File 4-12. Pairwise Comparisons for "The Contribution of Environmental Enrichment to Phenotypic Variation in Mice and Rats"

**Extended Data Table 4-12**. Pairwise comparisons for treated/manipulated controls and treated/manipulated enriched mice by each individual trait.

| Description | Trait Category | Mean | Standard  Deviation | Standard Error | 95% confidence interval | | t | df | p-value  (two tailed) |
| --- | --- | --- | --- | --- | --- | --- | --- | --- | --- |
|  |  |  |  |  | Lower | Upper |  |  |  |
| Main effect of housing | Behavior (all) | .117 | .942 | .086 | -.053 | .288 | 1.365 | 119 | .175 |
| Main effect of housing | Physiology  (all) | -.078 | .588 | .054 | -.186 | .030 | -1.435 | 116 | .154 |
| Main effect of housing | Anatomy | -.029 | .210 | .045 | -.122 | .06 | -.641 | 21 | .528 |
| Main effect of housing | Behavior (CNS) | .127 | 1.015 | .101 | -.071 | .326 | 1.271 | 102 | .207 |
| Main effect of housing | Behavior (other) | .058 | .143 | .035 | -.016 | .131 | 1.668 | 16 | .115 |
| Main effect of housing | Immune System | -.098 | .363 | .081 | -.268 | .071 | -1.213 | 19 | .240 |
| Main effect of housing | Molecules | -.061 | .690 | .0812 | -.225 | .102 | -.748 | 70 | .457 |
| Main effect of housing | Organ Function | -.717 | .989 | .571 | -3.17 | 1.74 | -1.257 | 2 | .336 |
| Main effect of housing | E-phys | -.0289 | .209 | .045 | -.122 | .064 | -.641 | 21 | .528 |
