## Extended Data File 4-13. Two Way ANOVAs for "The Contribution of Environmental Enrichment to Phenotypic Variation in Mice and Rats"

**Extended Data Table 4-13**. Two-way ANOVAs comparing multiple traits (all behavior, physiology, anatomy) by housing condition (environmental enrichment, standard housing) for the independent variable coefficient of variation (CV). Data presented for both rats and mice combined and separately.

| **Species** | **Housing** | **ANOVA** | **Tukey HSD** |
| --- | --- | --- | --- |
| **Rat and Mouse Data Combined** | *Naïve Controls and Naïve EE* | housing x trait: F(6, 1014) = 0.790, p = 0.643, ƞ^2^= 0.004  housing: F(1, 1014) = 0.349, p = 0.555, ƞ^2^= 0.0001  trait: F(6, 1014) = 7.336, p = 0.0001, ƞ^2^= 0.042 | *behavior (cns) vs behavior (other)*: p =0.017  *behavior (cns) vs anatomy*: p = 0.0001  *anatomy vs molecules*: p = 0.012 |
|  | *Treated/Manipulated Controls and Treated/Manipulated EE* | housing x trait: F(6, 1024) = 0.078, p =0.998, ƞ^2^=0.0001  housing: F(1, 1024) = 0.129, p =0.719, ƞ^2^= 0.0001  trait: (6, 1024) = 0.513, p = 0.799, ƞ^2^= 0.003 | N/A |
| **Rats** | *Naïve Controls and Naïve EE* | housing x trait: F(5, 542) = 0.422, p = 0.833, ƞ^2^= 0.004  housing: F(1, 542) = 0.007, p = 0.931, ƞ^2^= 0.0001  trait: F(5, 542) = 4.015, p = 0.001, ƞ^2^= 0.036 | *behavior (cns) vs anatomy*: p = 0.004 |
|  | *Treated/Manipulated Controls and Treated/Manipulated EE* | housing x trait: F(5, 696) = 0.013, p = 1.00, ƞ^2^= 0.0001  housing: F(1, 696) = 0.002, p = 0.963, ƞ^2^= 0.0001  trait: F(5, 696) = 0.460, p = 0.806, ƞ^2^= 0.003 | N/A |
| **Mice** | *Naïve Controls and Naïve EE* | housing x trait: F(6, 460) = 0.496, p = 0.811, ƞ^2^= 0.006  housing: F(1, 460) = 0.294, p = 0.588, ƞ^2^= 0.001  trait: F(6, 460) = 4.593, p = 0.0001, ƞ^2^= 0.057 | *behavior (cns) vs anatomy*: p = 0.001 |
|  | *Treated/Manipulated Controls and Treated/Manipulated EE* | housing x trait: F(6, 516) = 1.861, p = 0.086, ƞ^2^= 0.021  housing: F(1, 516) = 0.330, p = 0.566, ƞ^2^= 0.001  trait: F(6, 516) = 2.039, p = 0.059, ƞ^2^= 0.023 | N/A |
