## Extended Data File 4-14. Coefficient of Variation (CV)s for "The Contribution of Environmental Enrichment to Phenotypic Variation in Mice and Rats"

**Extended Data Table 4-14**. Coefficient of variation (CV) distributions for naïve standard housed (controls) and naïve environmental enriched (EE) rats and mice combined with treated/manipulated controls and treated/manipulated EE rats and mice in which all behavior, physiology, and anatomy traits are combined. CV ratios were used to determine whether the distribution of variation differed by environmental complexity. Calculated EE to control ratios of *CV* = [(*CV_EE_)/(CV_EE_ + CV_control_*)]. CV ratios tested as a function of housing complexity against the theoretical mean of 0.5 by a one-sample t-test.

| Description | Trait Category | t | df | p-value  (two tailed) | Mean Difference | 95% confidence interval | |
| --- | --- | --- | --- | --- | --- | --- | --- |
|  |  |  |  |  |  | Lower | Upper |
| Distribution of CV ratios | all traits combined | -1.638 | 1132 | .102 | -.00924 | -.0203 | .0018 |
| Distribution of CV ratios | Behavior | -1.565 | 659 | .118 | -.01157 | -.0261 | .0029 |
| Distribution of CV ratios | Physiology | -.685 | 472 | .493 | -.00598 | -.0231 | .0112 |
