## Extended Data File 4-17. Coefficient of Variation (CV)s for "The Contribution of Environmental Enrichment to Phenotypic Variation in Mice and Rats"

**Extended Data Table 4-17**. Coefficient of variation (CV) distributions for naïve standard housed (controls) and naïve environmental enriched (EE) rats and mice by each individual trait. CV ratios were used to determine whether the distribution of variation differed by environmental complexity. Calculated EE to control ratios of *CV* = [(*CV_EE_)/(CV_EE_ + CV_control_*)]. CV ratios tested as a function of housing complexity against the theoretical mean of 0.5 by a one-sample t-test.

| Description | Trait Category | t | df | p-value  (two tailed) | Mean Difference | 95% confidence interval | |
| --- | --- | --- | --- | --- | --- | --- | --- |
|  |  |  |  |  |  | Lower | Upper |
| Main effect of housing | Behavior  (all) | -.427 | 368 | .670 | -.00460 | -.0258 | .0166 |
| Main effect of housing | Physiology  (all) | -.567 | 249 | .571 | -.00759 | -.0339 | .0188 |
| Main effect of housing | Anatomy | .790 | 61 | .433 | .01742 | -.0267 | .0615 |
| Main effect of housing | Behavior (CNS) | -.881 | 307 | .379 | -.01079 | -.0349 | .0133 |
| Main effect of housing | Behavior (other) | 1.317 | 60 | .193 | .02668 | -.0138 | .0672 |
| Main effect of housing | Immune System | -1.432 | 24 | .165 | -.06355 | -.1551 | .0280 |
| Main effect of housing | Molecules | -.856 | 125 | .393 | -.01792 | -.0593 | .0235 |
| Main effect of housing | Organ Function | .837 | 17 | .414 | .04155 | -.0632 | .1462 |
| Main effect of housing | E-phys | .221 | 18 | .827 | .00642 | -.0545 | .0673 |
