## Extended Data File 4-24. Coefficient of Variation (CV)s for "The Contribution of Environmental Enrichment to Phenotypic Variation in Mice and Rats"

**Extended Data Table 4-24**. Coefficient of variation (CV) distributions for treated/manipulated standard housed (controls) and treated/manipulated environmental enriched (EE) mice in which all behavior, physiology, and anatomy traits are combined. CV ratios were used to determine whether the distribution of variation differed by environmental complexity. Calculated EE to control ratios of *CV* = [(*CV_EE_)/(CV_EE_ + CV_control_*)]. CV ratios tested as a function of housing complexity against the theoretical mean of 0.5 by a one-sample t-test.

| Description | Trait Category | t | df | p-value  (two tailed) | Mean Difference | 95% confidence interval | |
| --- | --- | --- | --- | --- | --- | --- | --- |
|  |  |  |  |  |  | Lower | Upper |
| Main effect of housing | all traits combined | -.840 | 236 | .402 | -.00876 | -.0293 | .0118 |
