## Extended Data File 4-26. Coefficient of Variation (CV)s for "The Contribution of Environmental Enrichment to Phenotypic Variation in Mice and Rats"

**Extended Data Table 4-26**. Coefficient of variation (CV) distributions for treated/manipulated standard housed (controls) and treated/manipulated environmental enriched (EE) mice by each individual trait. CV ratios were used to determine whether the distribution of variation differed by environmental complexity. Calculated EE to control ratios of *CV* = [(*CV_EE_)/(CV_EE_ + CV_control_*)]. CV ratios tested as a function of housing complexity against the theoretical mean of 0.5 by a one-sample t-test.

| Description | Trait Category | t | df | p-value  (two tailed) | Mean Difference | 95% confidence interval | |
| --- | --- | --- | --- | --- | --- | --- | --- |
|  |  |  |  |  |  | Lower | Upper |
| Main effect of housing | Behavior  (all) | -1.230 | 119 | .221 | -.01863 | -.0486 | .0114 |
| Main effect of housing | Physiology  (all) | .096 | 116 | .924 | .00137 | -.0270 | .0297 |
| Main effect of housing | Anatomy | -.448 | 21 | .659 | -.01335 | -.0753 | .0486 |
| Main effect of housing | Behavior (CNS) | -.877 | 102 | .383 | -.01517 | -.0495 | .0191 |
| Main effect of housing | Behavior (other) | -1.841 | 16 | .084 | -.03962 | -.0852 | .0060 |
| Main effect of housing | Immune System | .788 | 19 | .440 | .03536 | -.0586 | .1293 |
| Main effect of housing | Molecules | -.922 | 70 | .360 | -.01514 | -.0479 | .0176 |
| Main effect of housing | Organ Function | 2.193 | 2 | .160 | .18150 | -.1746 | .5376 |
| Main effect of housing | E-phys | -.448 | 21 | .659 | -.01335 | -.0753 | .0486 |
